# Management implications based on diversity patterns under climate change scenarios in a continental island biodiversity hotspot

**DOI:** 10.1101/2020.03.01.971143

**Authors:** Konstantinos Kougioumoutzis, Ioannis P. Kokkoris, Maria Panitsa, Panayiotis Trigas, Arne Strid, Panayotis Dimopoulos

**Affiliations:** Department of Ecology and Systematics, Faculty of Biology, National and Kapodistrian University of Athens, Panepistimiopolis, Greece, GR 15784; Division of Plant Biology, Laboratory of Botany, Department of Biology, University of Patras, Patras, Greece, GR 26504; Laboratory of Systematic Botany, Department of Crop Science, Agricultural University of Athens, Athens, Greece, GR 11855; Bakkevej 6, DK-5853 Ørbæk

**Author notes:** **Corresponding author:** Konstantinos Kougioumoutzis.

**Keywords:** CANAPE, conservation, EDGE, environmental management, extinction risk, Species Distribution Modelling

## Abstract

In the Anthropocene era, climate change poses a great challenge in environmental management and decision-making for species and habitat conservation. To support decision-making, many studies exist regarding the expected vegetation changes and the impacts of climate change on European plants, yet none has investigated how climate change will affect the extinction risk of the entire endemic flora of an island biodiversity hotspot, with intense human disturbance. Our aim is to assess, in an integrated manner, the impact of climate change on the biodiversity and biogeographical patterns of Crete and to provide a case-study upon which a cost-effective and climate-smart conservation planning strategy might be set. We employed a variety of macroecological analyses and estimated the current and future biodiversity, conservation and extinction hotspots in Crete, as well as the factors that may have shaped these distribution patterns. We also evaluated the effectiveness of climate refugia and the NATURA 2000 network (PAs) on protecting the most vulnerable species and identified the taxa that should be of conservation priority based on the Evolutionary Distinct and Globally Endangered (EDGE) index, during any environmental management process. The highlands of Cretan mountain massifs have served as both diversity cradles and museums, due to their stable climate and high topographical heterogeneity. They are also identified as biodiversity hotspots, as well as areas of high conservation and evolutionary value, due their high EDGE scores. Due to the ‘escalator to extinction’ phenomenon and the subsequent biotic homogenization, these areas are projected to become diversity ‘death-zones’ in the near future and should thus be prioritized in terms of conservation efforts and by decision makers. In-situ conservation focusing at micro-reserves and ex-situ conservation practices should be considered as an insurance policy against such biodiversity losses, which constitute cost-effective conservation measures. Scientists and authorities should aim the conservation effort at areas with overlaps among PAs and climate refugia, characterized by high diversity and EDGE scores. These areas may constitute Anthropocene refugia. Thus, this climate-smart, cost-effective conservation-prioritization planning will allow the preservation of evolutionary heritage, trait diversity and future services for human well-being and acts as a pilot for similar regions worldwide.

## 1. Introduction

Earth has entered a new geological epoch, the Anthropocene (Zalasiewicz et al., 2015), characterized by human-induced temperature, thus placing immense impacts upon natural, atmospheric and hydrological processes. Consequently, global ecosystem health is severely challenged, with shifts in biotic composition and ecological linkage disruption being among the major threats of human activity worldwide. Species unable to survive to such a novel adaptive matrix are on an inevitable path to extinction (Urban et al., 2016), often causing a deadlock in nature management and protection practices. Extinction threat might be more prominent on endemic species suffering from habitat loss and fragmented populations (Warren et al., 2013), since they have narrower geographical ranges and environmental niches. Climate change is projected to rapidly change community structure in the near future, likely surpassing the land-use effects by the 2070s (Newbold, 2018) and significantly altering terrestrial biodiversity (Warren et al., 2013). On a global analysis of future climate change impacts, ca. 60% of plant species examined were predicted to lose more than half of their current climatic range in the coming decades, possibly leading to a substantial global biodiversity decrease and ecosystem function degradation (Warren et al., 2013). Even though plants might be more resistant to extinction compared to animals (Wing, 2004), climate change could elevate the estimated extinction rates (Urban, 2015). Nearly 20-30% of species would face high extinction risk if global temperature rises beyond 2-3 °C above pre-industrial levels (Fischlin et al., 2007). In order to avoid a 1.5 °C rise in global temperature until 2052, unprecedented changes should be made at a global scale (IPCC, 2018). In this context, it is important to follow a ‘climate-smart’ conservation planning strategy to efficiently conserve as many species/communities as possible (Groves et al., 2012). This strategy relies on the identification of climate refugia, i.e., relatively climatically stable regions for many species of high conservation value which are characterised by high endemic and genetic diversity (Sandel et al., 2011).

The Mediterranean Basin is a biogeographically complex area, with high levels of endemism and diversification, due to its high topographic and climate heterogeneity (Médail, 2017). It constitutes the second largest terrestrial biodiversity hotspot in the world (Médail and Myers, 2004) with nearly 12500 plant species endemic to the region, most of which have an extremely narrow geographical range (Médail, 2017). The Mediterranean islands have high overall and endemic species richness (Médail, 2017). Plant endemism varies between 9-18% on the largest islands and reaches up to 40% in the high-altitude zone of their mountain ranges (Médail, 2017). Several of these islands constitute biodiversity hotspots and climate refugia (Médail and Diadema, 2009). Crete (S Greece) is rendered the hottest endemism hotspot of the Mediterranean Basin, owing its high species number and endemism levels to long-term geographical isolation and climate stability, as well as to its high environmental and topographical heterogeneity (Médail, 2017).

The current global temperature will probably continue to rise (IPCC, 2018) and has already forced species to migrate into higher altitudes, changed flowering period/duration and in some cases, leading species to extinction (e.g., Thackeray et al., 2016). These impacts will most likely accelerate (Urban, 2015) and are expected to be more severe for species occurring on mountain ranges, due to the ‘escalator to extinction’ phenomenon (Urban, 2018). The Mediterranean is a global hotspot of vulnerable species (Pacifici et al., 2015) and among the regions expected to experience the largest changes in climate (e.g., Giorgi and Lionello, 2008), possibly leading to a ca. 545 m upward vegetation shift and a 50-80 km northward shift in latitude, with these impacts being more prominent on islands and mountain summits (Médail, 2017). However, the Mediterranean Basin has experienced few plant species extinctions (22 taxa – 0.17% of the total; Domina et al., 2015). Most Mediterranean countries have their own Red List of threatened plants. Nevertheless, regarding Crete ca. 30% (58 taxa) of the Cretan single island endemics (SIE_c_) have been assessed and only five are considered facing imminent extinction (Phitos et al., 2009). Several SIE_C_ occur on the island’s mountain ranges and comprise small and isolated populations, thus being prone to climate change impacts.

New light can be shed upon the mechanisms underlying species’ possible extinction and their subsequent conservation needs when integrating phylogenetic diversity metrics into conservation-centred analyses (Morelli and Møller, 2018). Evolutionary distinct species usually have unusual phenotypes, rare ecological roles and increased functional importance (e.g., Cadotte et al., 2008). As such, they have great conservation value, since their loss cannot be readily replaced. Thus, if evolutionary isolated species are to become extinct, this would result into great evolutionary loss. Therefore, any conservation plan needs to take into account the advantage of the conservation prioritization of evolutionary distinct species, as it allows the preservation of evolutionary heritage, trait diversity and future services for human well-being (Veron et al., 2018).

Correlative species distribution models (SDMs) are widely used to identify which species are most vulnerable, as well as to answer how, why, where and when these species are vulnerable (Foden et al., 2019). There is ample evidence that the expectations from correlative SDMs have matched recent population trends focusing on birds, mammals and plants, in decreasing order and they provide appropriate results when the aim is to estimate a species’ extinction risk (Pacifici et al., 2015). To our knowledge, even though continental-wide studies exist regarding the expected vegetation changes and the impacts of climate change on European plants (e.g., Berry et al., 2017), none has investigated how climate change will affect the extinction risk of the entire endemic flora of an island biodiversity hotspot, such as Crete. Our aim is to assess, in an integrated manner, the impact of climate change on the biodiversity and biogeographical patterns of the endemic plants of Crete and to provide a case-study upon which a cost-effective and climate-smart conservation planning strategy might be set.

## 2. Material and Methods

### 2.1 Study area

We focus on Crete (Fig. 1), the fifth largest island in the Mediterranean Basin and the richest island hotspot of Europe in terms of endemic plant species (Médail, 2017).

**Figure 1.**
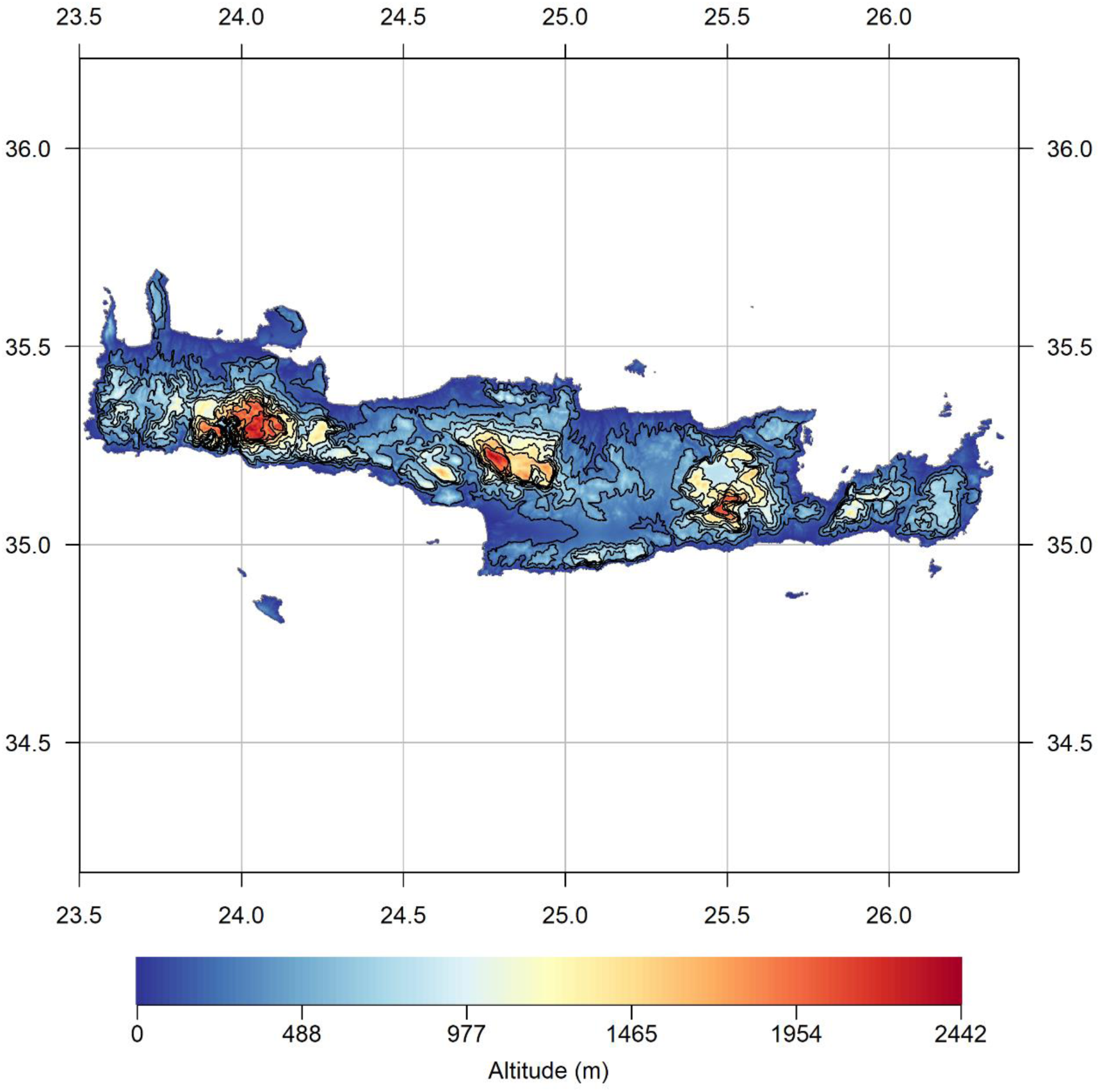
The island of Crete, along with the altitudinal contours at a 100m increment.

Crete is the largest (8836 km^2^) and most topographically complex island in the Aegean archipelago with more than 50 mountain peaks exceeding 2000m a.s.l. The island is climatically compartmentalized in a W-E axis and is characterized by a sharp altitudinal gradient (0-2456 m a.s.l.). There are three main mountain massifs present on Crete (Lefka Ori, Idi and Dikti), with an entirely different climate than the warm and dry lowland plains.

Geologically, Crete comprises mostly limestones, is part of the Hellenic Arc and was formed in response to the subduction of the African plate beneath the Aegean (Higgins, 2009).

Its palaeogeographical history is rather complex and determined by two main geological events that created important dispersal barriers (for a thorough review of the Aegean’s palaeogeographical history see Sakellariou and Galanidou, 2016): i) the isolation of Crete from Karpathos’ and Cyclades’ island groups 12 Mya and ii) the isolation of Crete from Peloponnese after the Messinian salinity crisis (Manzi et al., 2013).

### 2.2 Environmental data

An initial set of 43 predictors has been constructed (see Appendix A in Supporting Information), all at a 30 arc-sec resolution, including:

a. sixteen climatic variables based on the 19 bioclimatic variables from WorldClim for current and future climate conditions.
b. three Global Circulation Models (GCMs) that are rendered more suitable and realistic for the study area’s future climate.
c. two different IPCC scenarios from the Representative Concentration Pathways (RCP) family: the mild RCP2.6 and the severe RCP8.5.
d. seven soil variables providing predicted values for the surface soil layer at varying depths.
e. elevation data from the CGIAR-CSI data-portal (Jarvis et al., 2008) aggregated and resampled to match the resolution of the other environmental variables.

From this set of predictors, only seven were not highly correlated (Spearman rank correlation < 0.7 and VIF < 5 – see Appendix A).

### 2.3 Climate refugia

We located sites in Crete that are climatically stable (sensu Owens and Guralnick, 2019) for the past 4 My, using paleoclimatic data obtained from Brown et al. (2018) and Gamisch (2019), the framework of Owens and Guralnick (2019) and the ‘climateStability’ 0.1.1 package (Owens, 2019). Climate refugia as designated here refer to macro-refugia according to Ashcroft (2010).

### 2.4 Species Occurrence Data

Crete hosts 2240 native plant species, 183 of which are single island endemics (SIE_C_) (Dimopoulos et al., 2013, 2016; Strid, 2016). Based on the most extensive and detailed database [Flora Hellenica Database, Strid (ongoing)], we compiled a dataset of all the SIE_C_ (8773 occurrences). The final dataset includes 172 SIE_C_, since we modelled only those SIE_C_ with at least three locations (following van Proosdij et al., 2016).

Data cleaning and organizing procedure follows Robertson et al. (2016 – see Appendix A).

### 2.5 Phylogenetic tree

Since molecular data are not available for many of the SIE_C_, we used a “supertree” approach to generate our phylogenetic tree (see Appendix A). We subsequently calculated evolutionary distinctiveness (ED) and EDGE (Evolutionary Distinct and Globally Endangered) scores for each taxon, as well as the standardized effect size scores of phylogenetic diversity (PD) for each grid cell (see Appendix A).

### 2.6 Species distribution models

#### 2.6.1 Model parameterization and evaluation

The realized climatic niche of each taxon was modelled by combining the available occurrence data to current environmental predictors in an ensemble modelling scheme (Araújo and New, 2007 - see Appendix A).

#### 2.6.2 Model projections

Calibrated models with a TSS score > 0.8 (to avoid poorly calibrated ones) were used to project the suitable area for each taxon under current and future conditions through an ensemble forecast approach (Araújo and New, 2007 - see Appendix A).

#### 2.6.3 Area range change

We assessed whether the 172 SIE_C_ will experience range contraction or expansion under future conditions (see Appendix A). Taxa were not assumed to have unlimited dispersal capability, since this assumption could be overoptimistic.

### 2.7 IUCN measures

We followed the Preliminary Automated Conservation Assessment (PACA) framework (Stévart et al., 2019), calculated the IUCN measures EOO (Extent of Occurrence) and AOO (Area of Occupancy) and assigned each SIE_C_ to a preliminary IUCN threat category according to the Criteria A and B under current and future conditions (see Appendix A). After taxa were assigned to an IUCN and a PACA category, the total number of taxa recorded and the proportion of taxa assessed under each category were estimated, by combining Criteria A and/or B.

### 2.8 Niche breadth

Levins’ inverse concentration measure of niche breadth (Levins, 1968) was computed for each taxon (see Appendix A). Niche breadth values range from 0 (specialists) to 1 (generalists) being thus comparable among taxa (Levins, 1968). Difference of niche breadth between the different IUCN threat categories, was investigated via a Kruskal-Wallis non-parametric test (KWA), as well as the difference of area range change between SIE_C_ with broad or narrow environmental niches (see Appendix A).

### 2.9 Hotspot analysis

For each climate scenario, we stacked the final binary maps of all 172 SIE_C_. From this overlay, we calculated for each 30-sec grid-cell the number of taxa that would find suitable environmental conditions there. We defined potential SIE_C_ hotspots as the 20% of cells that provide suitable environmental conditions to the highest number of taxa.

### 2.10 Generalized Dissimilarity Modelling analysis

We used Generalised Dissimilarity Modelling (GDM) to model pairwise plant community compositional dissimilarity (Sorensen’s dissimilarity) within map grid cells across Crete. We used the same environmental variables as in the SDM analyses, with the significance of all variables assessed through a Monte Carlo permutation test (see Appendix A).

### 2.11 Significant Differences

To identify sites where predictions of hotspot location differed significantly between current and future environmental conditions, we applied the methodology proposed by Januchowski et al. (2010 - see Appendix A).

### 2.12 Biodiversity indices

We performed Categorical Analyses of Neo- and Paleo-Endemism (CANAPE), using the methods and reasoning described in Mishler et al. (2014) and Thornhill et al. (2016 – see Appendix A).

We assessed the impact of climate change on the distribution of the endemism hotspots by repeating the CANAPE analysis for every GCM/RCP combination included in our study.

We tested the relationships among phylogenetic endemism (PE) and relative phylogenetic endemism (RPE) with elevation, pH, mean diurnal range (MDR) and climate stability by employing spatial autoregressive models with spatially autocorrelated errors (SAR_err_-see Appendix A).

### 2.13 Future diversity and biogeographical patterns

We derived species composition in each grid cell under current and future climatic conditions by stacking the presences from the individual species models. We followed the framework outlined in Menéndez-Guerrero et al. (2019) regarding the assessment of the spatial biodiversity and biogeographical patterns under future climate scenarios (see Appendix A).

### 2.14 Protected areas network and climate refugia overlap

We overlapped current and future CANAPE and hotspot results with the areas recognised as climate refugia, as well as with the protected areas (PA) network retrieved from the World Database on Protected Areas (see Appendix A). Based on each species’ current and future EOO, we calculated the irreplaceability of each PA and climate refugium in Crete for the current and future climate conditions. The irreplaceability index represents biotic uniqueness and quantifies the degree of overlap between each PA/climate refugium and the range of SIE_C_ (Le Saout et al., 2013). We investigated whether the degree of overlap differed between the current and future conditions via a Kruskal-Wallis non-parametric test.

## 3. Results

### 3.1 Model performance

The models for most SIE_C_ had sufficient predictive power (TSS ≥ 0.7 – mean TSS: 0.94; Table S1 in Appendix B). The resulting habitat suitability maps were converted into binary maps and then compared to the binary maps obtained for each GCM, RCP and time-period for each SIE_C_. Precipitation-related variables had the highest contribution among the response variables for most SIE_C_ (53.5%), while temperature-related variables and soil pH were important for a smaller fraction of SIE_C_ (42.4% and 4.1%, respectively). Full details and analyses of model performance are given in Table S1.

### 3.2 Area range change

All SIE_C_ species will experience range contraction, with varying magnitude across taxa, GCMs and RCPs (11.7% - 100.0%; Table S1), but the median range contraction was 98.3% (see Table S2 for the median values for each GCM and RCP).

### 3.3 IUCN measures

For the first time, we provide a preliminary assessment of every SIE_C_ occurring in Crete according to the IUCN Criteria A and B (Fig. 2; Table S1). At present, 58.7% of SIE_C_ are facing imminent extinction (Fig. 2) and 87.8% are characterised as Likely Threatened (LT) under the PACA categories (Table S1). This phenomenon could be greatly deteriorated, since under any GCM and RCP included in our analyses, many of these species are projected to become extinct (median EX%: 48.85% - Fig. 2, Table S1). The distribution and the proportion of taxa assessed as Threatened either under the IUCN or the PACA categories under both Criteria A and B are not uniform across Crete (Fig. 3). These taxa are mostly concentrated at the Cretan mountain massifs, with the highest proportion of them occurring in Lefka Ori (Fig. 3). These high-altitude areas are predicted to become extinction hotspots under any GCM/RCP (Fig. S1).

**Figure 2.**
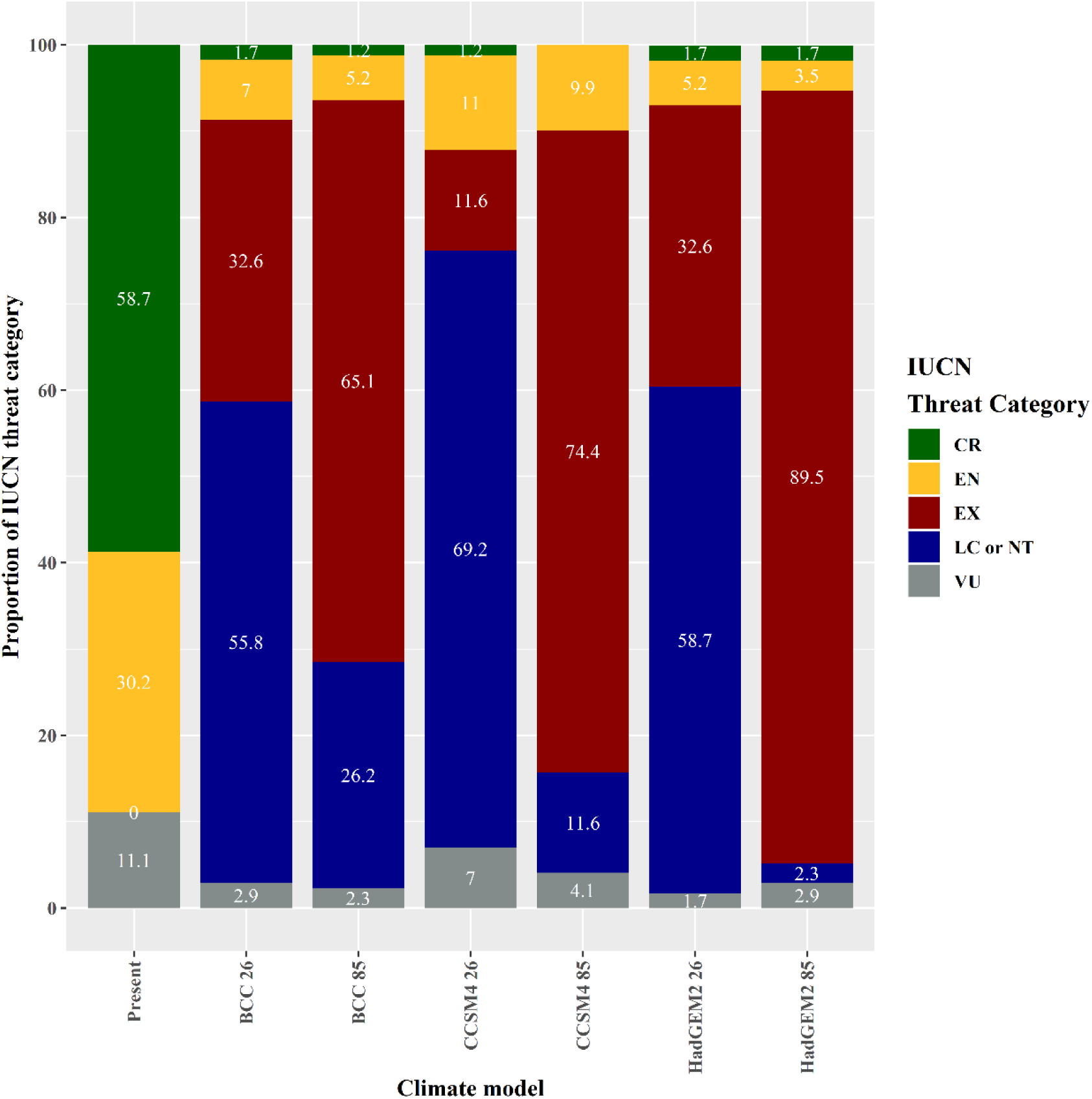
Proportion of the Cretan Single Island Endemics included in our analysis under the IUCN threat categories for the current conditions according to both Criterion A and B, as well as for every GCM and RCP considered in the present study.

**Figure 3.**
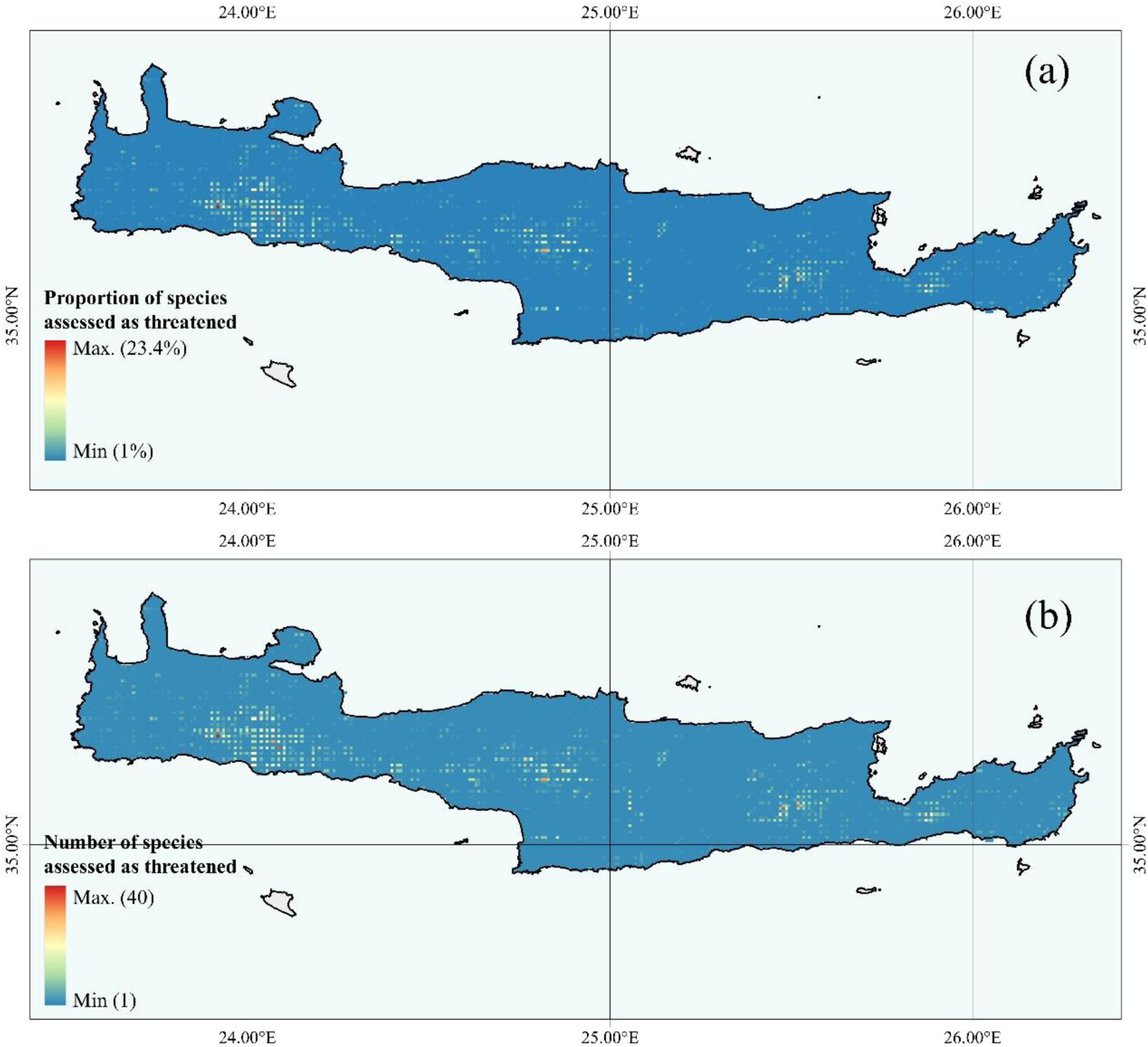
(a) Proportion of species and (b) number of species preliminary assessed as Threatened (either Critically Endangered, Endangered or Vulnerable) following Criteria A and B.

### 3.4 Niche breadth

The median niche breadth was 0.41 for the SIE_C_ and 25% of the SIE_C_ are considered to have a narrow niche breadth (Table S1). Niche breadth differed significantly between all the IUCN categories (KWA: H = 56.6, d.f. = 3, p < 0.05) and species with narrow niches have higher extinction probability for every GCM/RCP (Table S1).

### 3.5 EDGE

Based on both Criteria A and B, the SIE_C_ have EDGE scores in the range 3.57-8.74, with most species (75%) falling in an EDGE score class of 3.57-6.39. Eleven species (Table S1) have an EDGE score exceeding 7.0, belonging to eleven different families and all are considered as CR taxa (Table S1).

### 3.6 Hotspot analysis

The eastern, central and high-altitude parts of Crete are more likely to lose most of the SIE_C_ occurring there (Figs. S2-8). The SIE_C_ hotspots are currently located above 1500 m a.s.l. in Crete (Fig. S2), but all of them are expected to shift downwards – even below 1000 m a.s.l. under any GCM/RCP (Fig. S3-8) and these shifts are significantly different in absolute or relative terms (Figs. S9-20).

### 3.7 GDM analysis

GDM helped disentangle the relative contribution of geographic and environmental drivers of SIE_C_ community composition across time and space. Our model explained 26.6% of deviance in SIE_C_ composition (Figs. S21-22). Under any GCM/RCP, the low-elevation area in W Crete is predicted to exhibit the greater floristic turnover than any other region, followed by the high-altitude Cretan mountain ranges (Figs. S23-28). The most important gradient for determining SIE_C_ turnover was annual potential evapotranspiration (PET), followed by geographical distance (Fig. S21). Soil pH was the weakest predictor of SIE_C_ turnover. The fitted functions describing the turnover rate and magnitude along each gradient were nonlinear, with turnover rate varying with position along gradients (Fig. S21). An acceleration zone in SIE_C_ turnover was evident for PET and short geographical distances, while a sharp compositional transition was apparent along the precipitation of the wettest month gradient (Fig. S21).

### 3.8 Biodiversity indices

Higher and lower PD than expected was found in 21 and 86 sites, respectively (Fig. S29). These sites did not differ significantly in terms of elevation or pH preferences (both occur above 850 m a.s.l.), but the non-significant sites were located in lower elevation areas with higher pH (KWA: H = 261.93, d.f. = 5, p < 0.05; Table S3).

The CANAPE analyses revealed 208 sites as containing more PE than expected (Fig. 4a), which are mainly concentrated within and at the periphery of the four Cretan mountain massifs. Areas of mixed-endemism are the most common, followed by paleo- and neo-endemism areas (156, 26 and 21, respectively). Centres of paleo- and mixed-endemism occur in significantly higher altitude than not-significant sites (KWA: H = 15.73, d.f. = 4, p < 0.01; Table S4). Most paleo-endemism areas occur in western Crete and on the mountain massifs of eastern Crete (Fig. 4a). Overall results are congruent, regardless the grid resolution (Fig. S30).

**Figure 4.**
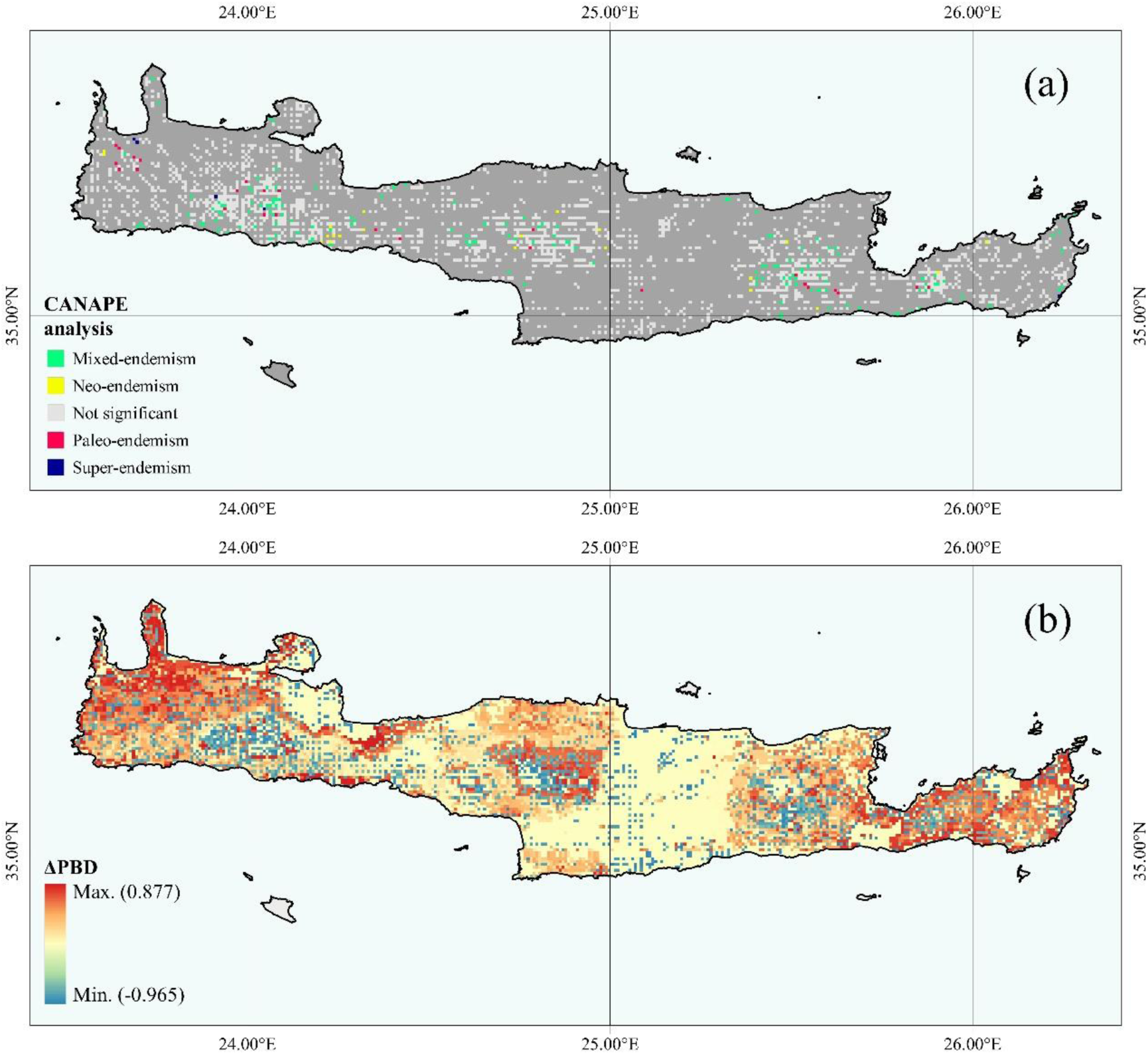
(a) Map of significant phylogenetic endemism (PE) identified by the Categorical Analysis of Neo- And Paleo-Endemism (CANAPE) analysis for 172 Cretan Single Island Endemics. Red squares: centres of mixed-endemism (i.e. mix of neo- and paleo-endemism). Blue squares: centres of neo-endemism; Purple squares: centres of paleo-endemism. Orange squares: centres of super-endemism. (b) Map of Crete showing predicted distribution of areas of biotic homogenization with respect to Cretan SIE species diversity, according to the CCSM4 26 GCM/RCP. Red areas indicate a decrease in beta-diversity (biotic homogenization). Blue areas indicate an increase in beta-diversity (biotic heterogenization). Analyses were performed under the assumption that a species cannot have unlimited dispersal capability, since this assumption could be overoptimistic.

The occurrence of the centres of endemism is projected to shift downwards (except for the super-endemism centres) in the future under any GCM/RCP (Figs. S31-36; Table S5) and these changes are significantly different for the mixed- and paleo-endemism centres mainly (KWA: H = 344.17, d.f. = 9, p < 0.01).

The SAR_err_ models indicate that elevation is the most important predictor of both PE and RPE (GR^2^ = 16.1% and 5.9%, respectively; Table S6), followed by MDR and CS, respectively.

### 3.9 Future diversity and biogeographical patterns

#### 3.9.1 Changes in ΔEDGE

Patterns of change regarding the EDGE index show that the high-altitude areas of Crete are currently identified as hosting assemblages of great evolutionary distinctiveness facing immediate extinction risk (Fig. S37). These areas are projected to become extinction hotspots under any GCM/RCP (Figs. S38-39). In the future, the mid-altitude areas are predicted to take up their place (Figs. S38-39).

### 3.9.2 Changes in ΔBD

Areas with high SIE_C_ β_sim_ are currently concentrated in high altitudes, mainly across the four Cretan mountain massifs (Fig. 5a). On the other hand, β_sim_ is relatively low over most of the lowlands and especially in the coastal area of northern Crete (Fig. 5a). β_sim_ is projected to generally increase in western Crete and in the mid-elevation areas (1000-1500 m a.s.l. – a trend towards biotic heterogeneity) and this pattern is projected to be greatest in a small coastal area of SW Crete (Fig. 4b). An entirely different pattern emerged mainly in the Cretan highlands, where β_sim_ is predicted to decrease (i.e., a trend towards biotic homogenization – Figs. 4b–5).

**Figure 5.**
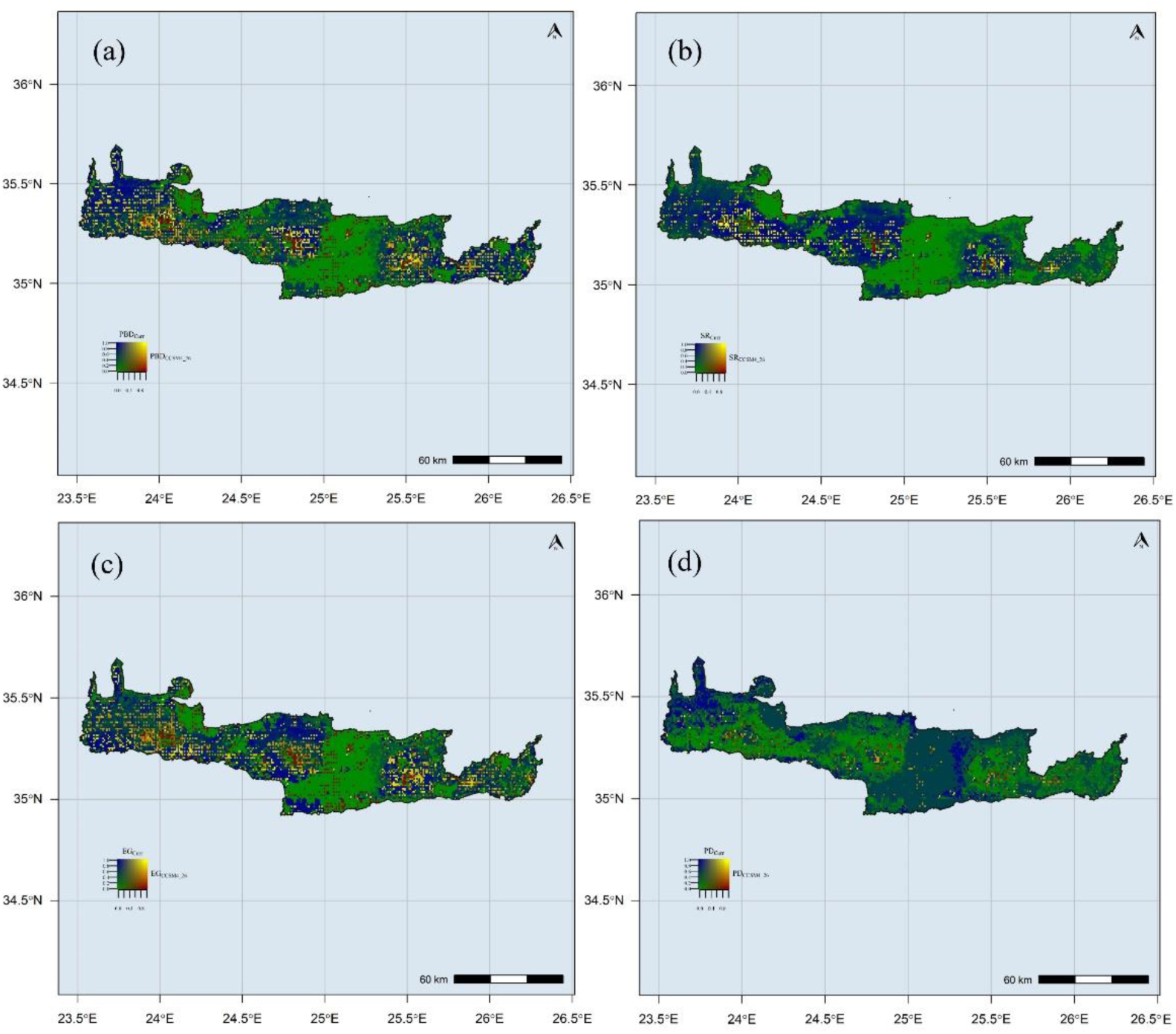
Bivariate maps depicting relative changes in biodiversity measures for Cretan SIE between the present and the CCSM4 26 GCM/RCP. Colours indicate the relative amount of change. The red and blue end of the spectrum indicate reductions and increases, respectively. Each transition in colour shading indicates a 10% quantile shift in the value of the variables. (a) beta-Diversity, (b) species richness (SR), (c) average level of ecological generalism (EG), (d) phylogenetic diversity (PD). Yellow areas indicate sites with high current beta-diversity that will continue to have proportionally high beta-diversity. Blue areas indicate sites that beta-diversity is predicted to increase in the future. Green areas indicate sites where beta-diversity will remain largely unchanged. We used a function generated by José Hidasi-Neto to generate the map (http://rfunctions.blogspot.ca/2015/03/bivariate-maps-bivariatemap-function.html).

The most parsimonious model for all GCMs/RCPs was the full GAM including ΔEG, ΔPD, ΔSR and elevation (Table S7, ΔAIC < 2), which explained 71.2-96.6% of the total variance in β_sim_ (Table S7). Both ΔPD and ΔSR were negatively correlated with β_sim_ change, while ΔEG was positively correlated with β_sim_ change (Fig. 7; Figs. S40-44). Here we focus on the results obtained for the CCSM4 2.6 GCM/RCP, as it shows the highest similarity with the current conditions (see *Changes in biogeographical patterns* below) and the patterns and trends do not deviate between the different GCMs and RCPs. The increasing heterogeneity of western Crete and mid-elevation areas (Figs. 4b–5) is mainly driven by range expansion and local extinction of generalist and specialist species, respectively (Fig. 6; see also red areas in Fig. 5b-c). Range expansions and local extinctions tend to drive species richness decreases, resulting in phylogenetic clustering (Fig. 6d). Hence, future SIE_C_ assemblages will tend to be species-poor and comprised of taxa that are more closely related than current assemblages (Fig. 6d; see also blue areas in Fig. 5d). The same processes are largely responsible for the predicted biotic homogenization of the higher elevation areas in Crete, even though these areas are predicted to host more specialist species.

**Figure 6.**
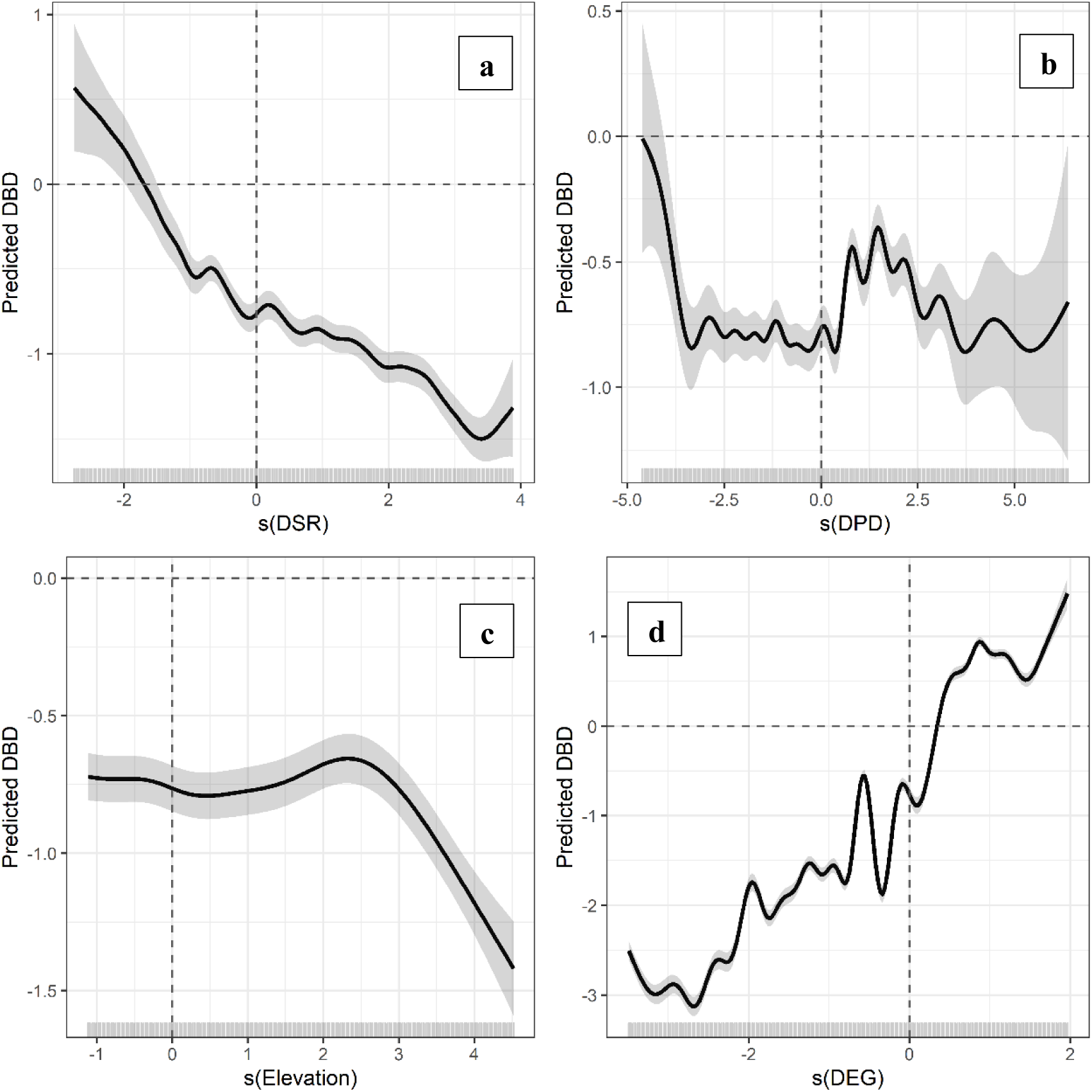
Biplots showing the predicted relationships between β_sim_ change and the four environmental predictors included in the best generalized additive model (GAM) for the CCSM4 26 GCM/RCP. (a) Change in species richness (ΔSR). (b) Change in phylogenetic diversity (ΔPD). (c) Change in elevation. (d) Change in the average level of ecological generalism (ΔEG). Fitted lines show the univariate GAMs with 95% confidence interval (dark grey). Rugs on the x-axes show the predictor values and how they are distributed. Labels on the y-axes indicate the smooth functions for the term of interest (ΔSR, ΔPD, ΔEG and elevation) and the estimated degrees of freedom (following the term). Values above and below the horizontal dashed line indicate heterogenization and homogenization, respectively. Values left and right of the vertical dashed line indicate in (a) species loss and gain, (b) PD decrease and increase and (d) assemblages composed by specialists and generalists, respectively.

### 3.10 Climate refugia and protected areas network overlap

Six areas were identified as climate refugia and are located along a W-E axis in Crete (Fig. S60). The largest and most species-rich climate refugium is located in the wider area of Lefka Ori in W Crete, while the smallest and species-poor is near the Asterousia mountain range in SC Crete (Fig. S60; Table S9). One and two of the climate refugia are significantly phylogenetically overdispersed and clustered, respectively (Table S9). In the future, all these areas are predicted to continue to serve as climate refugia, with different trends: the species-poor and species-rich refugia show a trend towards phylogenetic dispersion (PD increases) and clustering (PD decreases), respectively (Table S9).

The overlap between the PA network in Crete and endemism centres detected by CANAPE is rather high and ranges between 52-65% (Table S10; Fig. S61). The overlap between the areas recognised as climate refugia and endemism centres detected by CANAPE is lower than that reported for the PA network and ranges between 0-22% (Table S10; Fig. S62). Under any GCM/RCP, the predicted overlap for both the PA network and the climate refugia is lower than the one currently observed for most GCMs/RCPs and CANAPE categories (Table S10). The overlap between the PA network and the climate refugia ranges between 13.3-97.0% (Table S11). The percent overlap based on the recognised climate refugia of at least the neo-endemism centres is predicted to increase in the future, while the opposite trend is predicted for all the other CANAPE types (Tables S11-12). The mean irreplaceability index differs significantly between PAs and climate refugia (KWA: H = 56.6, d.f. = 1, p < 0.01; Fig. S63), as well as between PAs and any GCM/RCP (KWA: H = 103.3, d.f. = 6, p < 0.01). Regarding climate refugia, the mean irreplaceability index is significantly different only between the current and the HadGEM2 RCP 8.5 climate projection (KWA: H = 14.6, d.f. = 6, p < 0.05; Fig. S63). When taking out the effect of area, the mean irreplaceability index does not differ significantly between PAs and climate refugia, except for the current and the HadGEM2 RCP 8.5 climate conditions (KWA: H = 2.79, d.f. = 1, p = 0.09; Fig. S64).

## 4. Discussion

Climate change is projected to alter biodiversity and biogeographical patterns all over the globe (e.g., Enquist et al., 2019). The Mediterranean Basin is expected to face the largest changes in climate worldwide (e.g., Giorgi and Lionello, 2008), with these impacts being more prominent on islands and mountain summits (Médail, 2017). Here, we used the hottest endemic Mediterranean hotspot, Crete, as a case-study for management scenario building and decision making, based on an integrated assessment of climate-change impacts on biodiversity, biogeographical and conservation patterns. Our results highlight an augmented extinction risk for the majority of the SIE_C_, while pinpointing areas of high conservation and evolutionary value; they should alert conservation practices and management to take measures for biodiversity loss restrain and halt of further deterioration.

### 4.1 Diversity hotspots

The high-altitude areas of Crete are identified as species richness hotspots, with Lefka Ori, the western mountain massif, hosting the most SIE_C_ (Fig. S1). These areas are predicted to experience a sharp decline in species richness under any GCM/RCP and will no longer constitute biodiversity hotspots, as a result of the ‘escalator to extinction’ phenomenon (e.g., Urban, 2018). Due to upward species’ range shift and the extinction of high-altitude SIE_C_, mid-altitude areas will host an increasing number of species, thus intensifying the observed mid-domain effect of the SIE_C_ (Trigas et al., 2013). Most of the upward migrating species will probably be neo-endemic species, since neo-endemism centres occur at lower altitudes than paleo-endemism centres in Crete (Table S4).

### 4.2 Conservation assessment

We evaluated the potential conservation status of SIE_C_ using the IUCN Criteria A and B. Most of the SIE_C_ are potentially threatened by extinction (Table S1; Fig. 2). These potential extinctions were not randomly distributed among evolutionary groups, when accounting for mean future area loss (C_mean_ = 0.12; p < 0.01), a phenomenon observed in mammals as well (Davies and Yessoufou, 2013). The SIE_C_ will be highly vulnerable in the future, since up to 154 taxa are projected to become extinct (Table S1; Fig. 2). The high-altitude areas that now serve as diversity and conservation hotspots (due to high EDGE scores - Figs. S2 and S37) are projected to become extinction hotspots (due to very low ΔEDGE scores - Figs. S1 and S38-39) and should thus be prioritized in terms of conservation efforts. Eleven taxa (Table S1) should also be given high conservation priority, since they have a very high EDGE score. However, none of them is included in the IUCN Top-50 Mediterranean Island Plants initiative (de Montmollin and Strahm, 2005). Identifying conservation priorities and implementing effective actions are urgently needed and will become increasingly important, since human activities and land use are exerting unprecedented pressure on natural environments, leading to severe biodiversity changes (IPCC, 2018; Veron et al., 2016). In this context, locating areas with high EDGE and low ΔEDGE scores could help prioritize areas of high conservation and evolutionary importance and high extinction risk.

### 4.3 CANAPE

A fundamental block for understanding biodiversity patterns and consequently for conservation prioritization is to determine where species diversify (centres of neo-endemism – cradles) and persist (centres of paleo-endemism – museums) over evolutionary time. By this, we can decide whether or not PAs are indeed sheltering different aspects of biodiversity (Tucker and Cadotte, 2013) and improve conservation management in the unprecedented biodiversity decline setting which is currently underway (Humphreys et al., 2019). By incorporating evolutionary history and randomization processes, we were able to identify new areas of high evolutionary and conservation value. In total, 107 sites had PD higher or lower than expected by chance (Fig. S29), scattered across a W-E axis. Phylogenetically overdispersed sites of high conservation importance (Swenson et al., 2007) occur at higher altitudes in Crete and this may be related to a variety of reasons, such as the competitive exclusion of closely related taxa with high niche overlap, the colonization of phylogenetically distinct lineages (e.g., *Campanula*/*Roucela* species) or the high topographical heterogeneity of the Cretan mountain massifs that harbour distinctly adapted plant lineages (e.g., Edh et al., 2007).

In Crete, mixed-endemism centres occurring in higher altitude than the other types of endemism centres, prevail (Fig. 4). It seems that in Crete, montane regions do not act just as diversity cradles, but also as diversity museums – a pattern observed in other parts of the world as well (e.g., Dagallier et al., 2019). Most paleo-endemism centres occur in or near ravines/gorges in western Crete and on the mountain massifs of eastern Crete (Fig. 4a); these areas may have functioned as biogeographical museums (Chown and Gaston, 2000). Neo-endemism centres tend to occur in sub-montane altitude in Crete, rather near mixed-endemism centres (Fig. 4a); their recent diversification may have hindered their expansion. In the future, the occurrence of almost all types of endemism centres are projected to shift downwards under any GCM/RCP (Figs. S31-36; Table S5), suggesting that montane areas that have served as both diversity cradles and museums for a very long time, will probably become diversity ‘death-zones’ in the near future.

Climatic stability and high topographical/environmental heterogeneity may act in conjunction to provide the conditions needed for the simultaneous persistence of paleo-endemics and the diversification of neo-endemics (e.g., Azevedo et al., 2019). In this context, altitude – a proxy of environmental heterogeneity (e.g., Cabral et al., 2014) emerged as the most important predictor of PE and RPE in Crete, followed by MDR, pH and CS. Niche-related processes thus seem to play a minor role in generating endemism patterns in Crete, which is in line with previous studies at a coarser scale (Lazarina et al., 2019). Distinct endemism patterns may be found in sites with phylogenetically distinct assemblages, even if environmental similarity is high, due to phylogenetically conserved niche breadth and dispersal ability (Wiens and Graham, 2005). This could be the case in Crete, since SIE_C_’s niche breadth has a significant, though weak phylogenetic signal (C_mean_ = 0.11, p < 0.05). Our results lend weight to the suggestion that the interplay between topographical heterogeneity and climate may be linked with the configuration of centres of paleo- and neo-endemism on mountain massifs (Rangel et al., 2018), a phenomenon also recorded in the Neotropics (Azevedo et al., 2019). However, the low GR^2^ for both RE and RPE limits the scope of conclusions we can draw from these results, and so they should be regarded as informative rather than conclusive.

### 4.4 Future diversity and biogeographical patterns

We predict a trend towards biotic homogenization in the Cretan highlands, whereas an opposite trend towards biotic heterogenization will probably occur in low- and mid-elevation areas in Crete (Figs. 4b–5). Biotic homogenization is driven by the range expansion of generalists and the consequent local extinction of specialists, resulting in overall lower local species richness and phylogenetic diversity (Figs. 6 and S40-44). The same processes drive biotic heterogenization in Crete, however these assemblages will probably have higher species richness, due to the higher migration rate from the lowlands. The prominent homogenization of the Cretan highlands suggests that climate-driven extinctions of specialists may have greater impact on beta-diversity patterns than the generalists’ range expansion. After all, the most important gradient for determining SIE_C_ turnover is PET, followed by geographical distance (Figs. S21-22) and niche-based processes were found to exert some influence in the distribution of species occurring in Crete (Lazarina et al., 2019). These regional variation shifts in beta diversity could disrupt ecosystem functioning and provisioning of ecosystem services (Cardinale et al., 2012). To properly assess and predict future projections of landscape fragmentation, changes in demand, use and supply of ecosystem services should be taken into account, using a climate-change, management scenario-based approach (e.g. Kokkoris et al. 2019) and is particularly important for policy and decision-making related to land and resource use (Geijzendorffer et al., 2017). Our results are in line with the emergence of the ‘Homogenocene’ era, observed in other regions and organisms as well (e.g., Menéndez-Guerrero et al., 2019).

Currently Crete, a speciation centre (Panitsa et al., 2018) and a distinct Aegean phytogeographical region (Kougioumoutzis et al., 2017) is biogeographically subdivided into 14 different regions (Figs. S45-46), a result probably of paleogeographical history and local climate differentiation. This unique biogeographical compartmentalization seems to be at peril, since under any GCM/RCP it will drastically change (Figs. S47-58; Table S8), showing a trend towards biotic homogenization in a W-E axis, along Crete’s altitudinal range. This phenomenon seems to have a greater impact on the SIE_C_ of the Cretan highlands, since most of the habitat specialists occurring there, will probably be among the first to become extinct in the next decades.

### 4.5 Conclusions – Conservation and management implications

Incorporating climate-change projections is critically important for management strategies. Identifying climate refugia is an integral part of the ‘climate-smart’ conservation planning framework, since they constitute taxonomic and genetic diversity centres (e.g., Sandel et al., 2011), with macro-refugia – areas that sustain climatic suitability along broad spatiotemporal gradients (Ashcroft, 2010) – being generally more easily captured by GCMs. In the Mediterranean basin, where land-use change and the occurrence of fire events are expected to intensify (e.g., Turco et al., 2018), having a resilient PA system should be a priority. However, the current conservation strategy and management agenda for future conservation projects in Crete is based on the obligations of Dir. 92/43/EEC and especially for habitats and species of Annex I and II, respectively. These obligations include reporting for the conservation status and future trends for only a few species identified as under extinction risk by the present study. It is also evident that hesitancy toward anything but conventional conservation actions persists (Hagerman and Satterfield, 2013). Considering the worst-case scenario (i.e. extinction of the highest number of species by climate-change impact), in-situ conservation focusing at micro-reserves and ex-situ conservation practices should be considered, as an insurance policy against such losses of plant biodiversity (Hawkes et al., 2012), which constitute cost-effective conservation measures (Kadis et al., 2013). By this, our study urges scientists and authorities to aim the conservation effort at areas with overlaps among PAs and climate refugia, characterized simultaneously by high diversity and EDGE scores (including the 11 taxa with the highest EDGE score), as well as serving as mixed-endemism centres. These areas are qualified as future climate refugia and may actually constitute Anthropocene refugia (Monsarrat et al., 2019). By doing so, this ‘climate-smart’, cost-effective conservation prioritization planning will allow the preservation of evolutionary heritage, trait diversity and future services for human well-being (Veron et al., 2019).

Landscape connectivity as a prerequisite for successful migration is an important management issue. The current scale of habitat fragmentation is likely to hamper the potential success of migration that relies heavily upon the connectedness of populations across suitable environments (Hoffmann and Sgro 2011). Thus, measures to mitigate landscape fragmentation, as an active conservation strategy, should be included in ongoing and future Action Plans for habitats and species, based on climate-change projections and management scenarios. To properly assess and predict future projections of landscape fragmentation, changes in demand, use and supply of ecosystem services should be taken into account, using a climate-change, management scenario-based approach (e.g. Kokkoris et al. 2019). This procedure is important for policy and decision-making related to land and resource use (Geijzendorffer et al., 2017).

Future climatic projections can trigger the interest of the scientific community and the conservation society. However, to reach decision and policy making, raw scientific information is not always the appropriate mean of communication. All information produced by the present study should be transformed into tangible material using the ecosystem services approach and by this communicate the loss (via plant species extinction) of various ecosystem services and the related impacts into the socio-economic environment. In EU the MAES concept encapsulates this idea and acts as a management- and dissemination-tool among scientists and policy makers.

## Acknowledgments

This project has received funding from the Hellenic Foundation for Research and Innovation (HFRI) and the General Secretariat for Research and Technology (GSRT), under grant agreement No [2418].

## Appendix A. Extended Materials and Methods

## Appendix B. Supplementary Tables and Figures

