## Appendix A for "Management implications based on diversity patterns under climate change scenarios in a continental island biodiversity hotspot"

**Appendix A** Extended Materials and Methods

### *Environmental data*

Current and future climatic data were obtained from the WorldClim database (Hijmans et al., 2005) at a 30 sec resolution. We constructed 16 more climatic variables at the same resolution via the `envirem´ 1.1 (Title and Bemmels, 2017) package based on the 19 bioclimatic variables from WorldClim for current and future climate conditions. We selected three Global Circulation Models (GCMs) that are rendered more suitable and realistic for the study area's future climate based on McSweeney et al. (2015) and two different IPCC scenarios from the Representative Concentration Pathways (RCP) family: RCP2.6 (mild scenario) and RCP8.5 (severe scenario). Seven soil variables providing predicted values for the surface soil layer at varying depths, were obtained from the SoilGrids database (Hengl et al., 2017; https://www.soilgrids.org). Elevation data were derived from the CGIAR-CSI data-portal (http://srtm.csi.cgiar.org – Jarvis et al., 2008) and then aggregated and resampled using `raster´ 2.6.7 package (Hijmans, 2018) to match the resolution of the other environmental variables.

From this initial set of 43 predictors, only seven were not highly correlated (Spearman rank correlation < 0.7 and VIF < 5 – Dormann et al., 2013). Multicollinearity assessment was performed with the `usdm´ 1.1.18 package (Naimi et al., 2014).

### *Species Occurrence Data*

Crete hosts 2240 native plant species, 395 of which are Greek endemics and 183 are single island endemics ($SIE_C$ – Dimopoulos et al., 2013, 2016; Strid, 2016). Based on the most extensive and detailed database [Flora Hellenica Database, Strid (ongoing)] of plants occurring in Greece (~1M occurrences), we compiled a dataset of all the $SIE_C$ present in Crete (8773 occurrences). All $SIE_C$ were cross-checked for synonyms, following the nomenclature proposed by Dimopoulos et al. (2013, 2016) and Strid (2016). Our final dataset includes 172 $SIE_C$, since we modelled only those $SIE_C$ with at least three locations (following van Proosdij et al., 2016).

To avoid pseureplication and associated spatial sampling biases, we selected occurrences with a minimum distance from each other (1 km). We removed the fewest records necessary to substantially reduce the effects of sampling bias, while simultaneously retaining the greatest amount of useful information. This subsampling reduces the spatial aggregation of records to prevent that SDMs reflect a possible over-representation of environmental conditions associated with regions of higher sampling, which hinder interpretation and application of models (Elith et al., 2010; Aiello-Lammens et al., 2015). This data cleaning and organizing procedure followed the protocols as set out in Robertson et al. (2016) and we used the 'biogeo' 1.0 (Robertson et al., 2016) and 'spThin' 0.1.0 (Aiello-Lammens et al., 2015) packages. We also evaluated whether any geographical sampling bias existed in our species occurrence data by comparing the statistical distance distribution observed in our dataset to a simulated distribution expected under random sampling via the 'sampbias' 0.1.1 (Zizka and Silvestro, 2020) package.

***Phylogenetic tree***

Since molecular data are not available for many of the $SIE_C$, we used a "supertree" approach to generate our phylogenetic tree. A phylogenetic tree was generated for all the $SIE_C$ included in our analyses, based on the recently published phylogeny of seed plants by Smith and Brown (2018) and Jin and Qian (2019). We used the GBOTB extended tree (i.e., GenBank taxa with a backbone provided by Open Tree of Life version 9.1) that contains 74,533 taxa and is the largest dated mega-tree for vascular plants (Jin and Qian, 2019). We appended any $SIE_C$ that were present in Crete, but missing from the phylogeny, by adding them next to a randomly selected congener, following Bruelheide et al. (2019) and Maitner et al. (2018). We did not add the missing taxa as polytomies to their respective genera, as this approach adds substantial bias to any ensuing analyses (Davies et al., 2012). We then pruned the phylogeny to keep only the 172 $SIE_C$.

Evolutionary distinctiveness (ED) was then calculated in the time-calibrated tree using the 'picante' 1.6.2 (Kembel et al., 2010) package. EDGE scores were calculated using the formula given in Isaac et al. (2007):

$$\text{EDGE} = \ln(1 + ED) + GE \times \ln(2) \quad (1)$$

where ED is the ED value of a species as calculated in 'picante' and GE is its weighted IUCN threat category [LC = 0; NT = 1; VU = 2; EN = 3; CR = 4] on a log scale.

We also estimated for each grid cell the phylogenetic alpha diversity (PD - sensu Faith 1992) of the species inhabiting each of the grid cells with the 'picante' 1.6-2 package (Kembel et al. 2010) and the standardized effect size scores with 'PhyloMeasures' 2.1 (Tsirogiannis and Sandel, 2017). We tested for non-random patterns in PD by estimating their standardized effect size (SES) scores as

$$\text{SES} = \frac{X_{obs} - mean(X_{null})}{s.d.(X_{null})} \quad (2)$$

where $X_{obs}$ is the observed score within each grid cell and mean ($X_{NULL}$) and standard deviation ($X_{NULL}$) are the mean and standard deviation of a null distribution of scores generated by shuffling taxa labels of the grid cell-by-species matrix 999 times. We assessed the statistical significance of the SES scores by calculating two-tailed p-values (quantiles) as

$$\text{p-values} = \frac{rank_{obs}}{runs+1} \quad (3)$$

where $rank_{OBS}$ is the rank of the observed scores compared with those of their null distributions, and runs is the number of randomizations (Kembel et al. 2010). SES scores with $p < 0.05$ and $p > 0.95$ were considered as significantly lower and higher than expected for a given PD value, respectively. Positive SES values indicate phylogenetic overdispersion, whereas negative SES values indicate phylogenetic clustering. The greater sensitivity of $SES_{PD}$ to more terminal structure makes it better suited to explore assembly processes working at finer temporal and spatial scales (Mazel et al. 2015).

### ***Species distribution models***

*Model parameterization and evaluation*

We modelled the realized climatic niche of each species by combining the available occurrence data with current environmental predictors with the 'biomod2' 3.3.7 package (Thuiller et al., 2009). We used three different modelling algorithms for species with more than ten occurrences: Random Forest (RF), Classification Tree Analysis (CTA) and Multiple Adaptive Regression Splines (MARS) in an ensemble modelling scheme, as ensemble forecasting integrates the results of multiple SDM algorithms into a single geographical projection for each time period, reducing the uncertainties associated with the use of a single model algorithm (Araújo and New, 2007; Araújo et al., 2019). Since these algorithms require presence/absence (PA) data, we generated PAs following the recommendations of Barbet-Messin et al. (2012) and pseudo-absences were generated at a minimum distance of 19.8 km from presence locations to reduce the probability of false absences. We chose that minimum distance due to the median autocorrelation of 19.7 km among the non-collinear environmental variables, which we computed with 'blockCV' 1.0.0 (Valavi et al., 2018) package. PA generation and model calibration was repeated 100 times per species to ensure that the selected pseudo-absences did not bias the final predictions. Regarding species with less than ten occurrences, we followed the ensemble of small models (ESM) framework (Breiner et al., 2015), which is suitable for modelling rare species (Breiner et al., 2015, 2017, 2018), using the RF algorithm (single-technique ESMs perform equally good as double ensembles – Breiner et al., 2018), which is robust to overfitting (Lawler et al., 2006). We calibrated ESMs by fitting numerous bivariate models, which were then averaged into an ensemble model using weights based on model performances. For all models, the weighted sum of presences was equal to that of the PAs. The models' predictive performance was evaluated via the True Skill Statistic (TSS; Allouche et al., 2006) based on a repeated (10 times) split-sampling approach in which models were calibrated with 80% of the data and evaluated over the remaining 20%. We used null model significance testing (Raes and ter Steege, 2007) to evaluate the performance of all models and estimated the probability that each model performed better than 100 null models. All models were found to outperform the null expectation at $P < 0.001$.

*Model projections*

Calibrated models were used to project the suitable area for each species in Crete under current and future conditions through an ensemble forecast approach (Araújo and New, 2007). The contribution of each model to the ensemble forecast was weighted according to its TSS score. Models with a TSS score < 0.8 were excluded from building projections, so as to avoid working with poorly calibrated models.

*Area range change*

To assess whether the 172 $SIE_C$ will experience range contraction or expansion under future conditions, we used the 'biomod' 3.3.7 package (Thuiller et al., 2009). Species were not assumed to have unlimited dispersal capability, since this assumption could be overoptimistic.

### ***IUCN measures***

We followed the Preliminary Automated Conservation Assessment (PACA) framework (Stévart et al., 2019) and calculated the standard IUCN measures EOO (Extent of Occurrence) and AOO (Area of Occupancy) and we assigned each $SIE_C$ to a preliminary IUCN threat category according to Criterion A and B under current and future conditions using the 'ConR' 1.1.1 package (Dauby, 2017). Under the same framework, we implemented assessments aligned with the parameters of Criterion A, using the R code provided by Stévart et al. (2019; https://github.com/gdauby/stevart_el_al_PACA). To calculate Criterion A under this approach, we used the CORINE land cover (CLC) data v.20 (https://land.copernicus.eu/pan-european/corine-land-cover/clc2018?tab=download). CLC layers 1-2 [apart from 223 (Olive groves) and 243-244 (Land principally occupied by agriculture, with significant areas of natural vegetation and Agro-forestry areas, respectively)] are directly linked to the main threats on $SIE_C$. After species were assigned to an IUCN and PACA category, we estimated the total number of taxa recorded and the proportion of taxa assessed under each IUCN and PACA category, under Criteria A and B separately, and by combining both criteria, i.e., a taxon would, for example, be categorized as CR if it is assessed as CR by at least one of the two criteria.

***Niche breadth***

Levins' inverse concentration measure of niche breadth (Levins, 1968) was computed for each taxon using `ENMTools´ 0.2 package (Warren et al., 2017). Niche breadth values range from 0 (specialists) to 1 (generalists) being thus comparable among taxa (Levins, 1968). Difference of niche breadth between the different IUCN threat categories, was investigated via a Kruskal-Wallis non-parametric test (KWA), as well as the difference of area range change between $SIE_C$ with broad or narrow environmental niches (above and below the first quartile of the niche breadth values of the dataset, respectively).

***GDM analysis***

We used Generalised Dissimilarity Modelling (GDM – Ferrier et al., 2007) to model pairwise plant community compositional dissimilarity (Sorensen's dissimilarity) within map grid cells across Crete as a response to environmental and spatial variables. GDM models species turnover as a set of non-linear monotonic functions along environmental gradients and recognises the curvilinear relationship between ecological dissimilarity and environmental distance, providing a sophisticated and realistic approach to correlative modelling of ecological communities. We used the same environmental variables as in the SDM analyses, with the significance of all variables assessed through a Monte Carlo permutation test (1000 repetitions). The results included a set of significant predictor variables, a unique fitted I-spline for each significant predictor variable describing the relationship between beta diversity and that gradient, and percent deviance explained by the model, the metric used by GDM to assess model fit. We plotted the I-splines to assess how magnitudes and rates of species turnover varied along and between gradients. To quantify the magnitude of turnover along each gradient and the relative importance of that gradient in driving species turnover, we summed the coefficients of the I-splines (each spline has three coefficients), which describes the proportion of compositional turnover explained by that variable and is determined by the maximum height of its I-spline (Ferrier et al., 2007; Fitzpatrick et al., 2013). To evaluate the unique contributions of environment and space in explaining species turnover, we partitioned the deviance resulting from sets of three GDMs that used either environmental variables, geographical distance or both as predictor variables (Borcard et al., 1992). We projected models trained on current environmental conditions onto the future climate models and scenarios described above. The resulting predicted species turnover is a function of the magnitude of climate change at a given location and at a given time, starting position along climatic gradients and the modelled rate of turnover at that position. All GDM analyses were performed with the 'gdm' 1.3.7 package (Manion et al., 2018).

***Significant Differences***

To identify sites where predictions of hotspot location differed significantly between current and future environmental conditions, we applied the methodology proposed by Januchowski et al. (2010), using the functions from the 'SDMTools' 1.1.221 package (VanDerWal et al., 2014). Significance of the cell-specific differences between predictions was calculated as the probability for any single difference relative to the mean and variance of all location-specific differences. The probability value represents the area under the curve of a Gaussian distribution defined by the mean and variance across all cells. The spatial surfaces representing the individual significance values were reclassified to indicate areas where the first distribution predicted significantly more suitable habitat (SD $\geq$ 0.975), where the second distribution did (SD $\leq$ 0.025) and where there was no significant difference between models.

***Biodiversity analyses***

We followed the CANAPE protocol for spatial phylogenetics analyses as set out in Laffan et al. (2010) and Mishler et al. (2014). We carried out all the relevant analyses in Biodiverse version 3.0 (Laffan et al., 2010). We first calculated phylogenetic endemism (PE – Rosauer et al. 2009) and relative phylogenetic endemism (RPE – Mishler et al. 2014). PE is the total branch length from the dated phylogenetic tree of the lineages present at a grid cell divided by the range sizes of the respective lineages. RPE is the ratio between PE measured from the original phylogeny in relation to the PE estimated from a phylogeny with equally distributed branch lengths (see Mishler et al. 2014 for more details). Relative phylogenetic diversity (RPD) is also a ratio that compares the phylogenetic diversity (PD) observed on the actual tree in the numerator to that observed on a comparison tree in the denominator. To make them easily comparable between analyses, the trees in both the numerator and the denominator are scaled such that branch lengths are calculated as a fraction of the total tree length. The comparison tree retains the actual tree topology but makes all branches of equal length. Thus, RPD is PD measured on the actual tree divided by PD measured on the comparison tree, while RPE is PE measured on the actual tree divided by PE measured on the comparison tree (Mishler et al., 2014). RPE is the basis for the Categorical Analyses of Neo- and Paleo-Endemism (CANAPE).

*Randomization tests*

We assessed the statistical significance of PD, PE, RPD and RPE by following Mishler et al. (2014) approach. We compared the actual PE and RPE values of each grid cells to the 999 values of a null distribution, using the 'rand_structured' option in Biodiverse. In this model, species occurrences in grid cells are randomly reassigned to grid cells without replacement, thus keeping constant both the total number of grid cells for each species and the SR of each grid cell. We ran 999 randomisations, calculating PD, PE, RPD and RPE for each run. These values formed a null distribution for each grid cell for use in non-parametric tests of the significance of observed values. We estimated p-values from a two-tailed distribution of values to identify areas with higher (> 0.975) or lower (< 0.025) PE or RPE than the null distribution (Mishler et al., 2014).

*CANAPE*

CANAPE is a two-step procedure discriminating grid cells with significantly high PE in neo- or paleo-endemism based on species occurrences and the dated phylogenetic tree (Mishler et al., 2014).

First, to determine whether a site is a centre of significantly high endemism, a grid cell needs to be significantly high (one-tailed test, $\alpha = 0.05$) in the numerator of RPE, the denominator or both.

If (and only if) grid cells pass one of those tests, then they are divided into four meaningful, non-overlapping categories of centres of endemism (Mishler et al., 2014). If a point is significantly high in the RPE ratio (two-tailed test, $\alpha = 0.05$), then it is a centre of paleo-endemism (contains significantly more endemic species on long branches). If a point is significantly low in the RPE ratio (two-tailed test, $\alpha = 0.05$), then it is a centre of neo-endemism (contains significantly more endemic species on short branches). If it is significantly high in both the numerator and the denominator (taken alone), but not significant for RPE, then it is a centre of mixed endemism. Mixed endemism can be interpreted as a centre of endemism having a mix of rare long and rare short branches, so not significantly dominated by either paleo-endemism or neo-endemism. The mixed endemism areas are further subdivided: those grid cells that are significantly high in both the numerator and the denominator at the $\alpha = 0.01$ level are termed super-endemic sites (i.e., highly significant concentration of endemic long and short branches – Mishler et al., 2014).

As CANAPE results might be sensitive to the grid cell size, CANAPE was also carried out with four different grid cell sizes across the study area: 2.5, 5 and 10 $km^2$. Overall results for the study area are congruent, regardless of the grid resolution.

All analyses were performed using Perl wrapper functions to run Biodiverse in R modified from https://github.com/NunzioKnerr/biodiverse_pipeline.

*Spatial autoregressive models*

We employed spatial autoregressive (SAR) models with spatially autocorrelated errors ($SAR_{err}$) to test the relationships among PE and RPE with elevation, pH, mean diurnal range (MDR) and climate stability (these predictors were not correlated; VIF < 2). All variables were standardized [i.e., (value-mean/standard deviation)] to enhance comparability of parameter estimates. We used correlograms of the residuals of both $SAR_{err}$ and generalized linear models (GLMs) to infer the degree of spatial autocorrelation, using functions from the ‘spdep’ 1.1.3 R package (Bivand and Wong, 2018). We selected the number of neighbours for the $SAR_{err}$ models so as to minimize the corrected Akaike Information Criterion (AICc). We then tested models for all combinations of variables and selected the best model (lower AICc).

***Future diversity and biogeographical patterns***

We derived species composition in each grid cell under current and future climatic conditions by stacking the presences from the individual species models. We used a grid cell resolution of ~1 km to match the resolution of the predictor variables. A grid cell was considered occupied if it overlapped any part of a species projected distribution (McKnight et al., 2007).

*Changes in species richness (ΔSR)*

We estimated projected ΔSR by subtracting future projected species richness (SR) from current SR. Negative and positive values represent projected species losses and gains, respectively.

*Changes in PD (ΔPD)*

We estimated current and future the standardized effect size PD scores as described above for each grid cell. Negative and positive ΔPD indicate that assemblages are projected to become increasingly clustered or overdispersed, respectively.

*Changes in ecological generalism (ΔEG)*

Niche breadth is a reasonable surrogate for ecological generalism-specialism of species (Brown, 1984). We derived the mean of the current niche breadth for the species present in each grid cell under current and future climate scenarios and calculated their difference. Negative and positive ΔEG indicate assemblages that are predicted to shift their composition towards a greater proportion of specialists and generalists, respectively (Menéndez-Guerrero et al., 2019).

*Changes in EDGE (ΔEDGE)*

We derived the mean of the current EDGE for the species present in each grid cell under current and future climate scenarios and calculated their difference. Negative and positive values indicate areas that are predicted to become extinction hotspots and coldspots, respectively.

*Changes in phylogenetic beta diversity (ΔBD)*

Phylogenetic beta diversity (PBD) is built upon two major components: turnover ($\beta_{sim}$ – species replacement) and nestedness ($\beta_{nes}$ – loss or gain of species), which may occur between nested or non-nested assemblages (Baselga, 2010, 2012; Legendre, 2014). PBD and its components were computed using the 'betapart' 1.3 R package (Baselga and Orme, 2012). We focused on $\beta_{sim}$ as it contributes more than $\beta_{nes}$ to overall beta diversity among sites (Baselga et al., 2012; Xu et al., 2015). We estimated projected ΔBD by subtracting future projected $\beta_{sim}$ from current $\beta_{sim}$. Negative and positive values indicate a trend towards biotic homogenization and heterogeneity, respectively (Menéndez-Guerrero et al., 2019).

We assessed whether changes in $\beta_{sim}$ were associated with ΔSR, ΔPD, ΔEG and elevation by fitting generalized additive models (GAMs) with the 'mgcv' 1.8.31 R package (Wood, 2017) in a model selection framework (Burnham and Anderson, 2003) to determine the best models for describing $\beta_{sim}$.

From the initial set of 43 predictors, only four were not highly correlated (Spearman rank correlation < 0.7 and VIF < 5 – Dormann et al., 2013). Multicollinearity assessment was performed with the 'usdm' 1.1.18 package (Naimi et al., 2014). All variables were standardized [i.e., (value-mean/standard deviation)] to help ensure model convergence and enhance comparability of parameter estimates. Model selection was based on Akaike's Information Criterion corrected for small sample sizes (AICc). We used the *dredge* function in the 'MuMIn' 1.15.6 package (Barton, 2017) to run a complete set of models with all possible combinations of the predictor variables and to identify the set of 'best models' according to the widely accepted criterion for different AICc values: ΔAICc < 2 (all models with ΔAICc < 2 are considered as equally parsimonious and as having relatively similar levels of support – Burnham and Anderson, 2003). If more than one model had ΔAICc < 2, we calculated the relative importance of each variable as the sum of AICc weights (Akaike weights – *w*AICc) for the models in which the variable was included (Burnham and Anderson, 2003; see also Cameron et al., 2013). Akaike weights are directly interpreted in terms of each model's probability of being the best supported for explaining the data (Burnham and Anderson,

2003; Cameron *et al*., 2013). Finally, we calculated the normalized root mean square error (RMSE) for each set of the 'best' models with the 'sjstats' 0.11.2 R package (Lüdecke, 2017).

*Changes in biogeographical patterns*
Based on the GDM results and following the framework of Fitzpatrick et al. (2011), we estimated the current and future bioregionalization of Crete via an unsupervised classification procedure, using two clustering algorithms: k-means and CLARA. We assessed the optimal number of clusters via the Silhouette index (Rousseeuw, 1987) for each time-period. Finally, we quantitively assessed the similarity of the different bioregionalizations via the V-measure index of spatial association (Rosenberg and Hirschberg, 2007; Nowosad and Stepinski, 2018). All analyses were performed using functions from the 'raster' 3.0.7 (Hijmans, 2018), 'cluster' 2.0.7-1 (Maechler et al., 2017), 'clusterCrit' 1.2.8 (Desgraupes, 2018) and 'sabre' 0.3.1 (Nowosad and Stepinski, 2018) R packages.

***Protected areas network and climate refugia overlap***
We overlapped current and future CANAPE and hotspot results with the areas recognised as climate refugia, as well as with the protected areas (PA) network retrieved from the World Database on Protected Areas (WPDA) using functions from the \`wpdar´ 1.0.0 (Hanson, 2019) and the \`sf´ 0.8.0 R package (Pebesma, 2018). Protected areas exclusively related to marine protection were excluded. Based on each species' current and future EOO, we calculated the irreplaceability of each PA and climate refugium in Crete for the current and future climate conditions. The irreplaceability index represents biotic uniqueness and quantifies the degree of overlap between each PA/climate refugium and the range of $SIE_C$ (Le Saout et al., 2013). We investigated whether the degree of overlap differed between the current and future conditions via a Kruskal-Wallis non-parametric test.

**References – Appendix A**

Aiello-Lammens, M.E., Boria, R.A., Radosavljevic, A., Vilela, B., Anderson, R.P., 2015. spThin: An R package for spatial thinning of species occurrence records for use in ecological niche models. Ecography (Cop.). 38, 541–545. https://doi.org/10.1111/ecog.01132

Allouche, O., Tsoar, A., Kadmon, R., 2006. Assessing the accuracy of species distribution models: Prevalence, kappa and the true skill statistic (TSS). J. Appl. Ecol. 43, 1223–1232. https://doi.org/10.1111/j.1365-2664.2006.01214.x

Araújo, M.B., New, M., 2007. Ensemble forecasting of species distributions. Trends Ecol. Evol. 22, 42–47. https://doi.org/10.1016/j.tree.2006.09.010

Araújo, M.B., Anderson, R.P., Barbosa, A.M., Beale, C.M., Dormann, C.F., Early, R., Garcia, R.A., Guisan, A., Maiorano, L., Naimi, B., others, 2019. Standards for distribution models in biodiversity assessments. Sci. Adv. 5, eaat4858.

Barbet-Massin, M., Jiguet, F., Albert, C.H., Thuiller, W., 2012. Selecting pseudo-absences for species distribution models: how, where and how many? Methods Ecol. Evol. 3, 327–338.

Barton, K., 2017. MuMIn: Multi-Model Inference.

Baselga, A., 2012. The relationship between species replacement, dissimilarity derived from nestedness, and nestedness. Glob. Ecol. Biogeogr. 21, 1223–1232.

Baselga, A., 2010. Partitioning the turnover and nestedness components of beta diversity. Glob. Ecol. Biogeogr. 19, 134–143. https://doi.org/10.1111/j.1466-

8238.2009.00490.x
Baselga, A., Gómez-Rodríguez, C., Lobo, J.M., 2012. Historical Legacies in World Amphibian Diversity Revealed by the Turnover and Nestedness Components of Beta Diversity. PLoS One 7, e32341. https://doi.org/10.1371/journal.pone.0032341
Baselga, A., Orme, C.D.L., 2012. betapart: an R package for the study of beta diversity. Methods Ecol. Evol. 3, 808–812.
Bivand, R.S., Wong, D.W., 2018. Comparing implementations of global and local indicators of spatial association. Test, 27(3), 716-748.
Borcard, D., Legendre, P., Drapeau, P., 1992. Partialling out the Spatial Component of Ecological Variation. Ecology, 73, 1045–1055.
Breiner, F.T., Guisan, A., Bergamini, A., Nobis, M.P., 2015. Overcoming limitations of modelling rare species by using ensembles of small models. Methods Ecol. Evol. 6, 1210–1218. https://doi.org/10.1111/2041-210X.12403
Breiner, F.T., Guisan, A., Nobis, M.P., Bergamini, A., 2017. Including environmental niche information to improve IUCN Red List assessments. Divers. Distrib. 23, 484–495.
Breiner, F.T., Nobis, M.P., Bergamini, A., Guisan, A., 2018. Optimizing ensembles of small models for predicting the distribution of species with few occurrences. Methods Ecol. Evol. 9, 802–808.
Brown, J.H., 1984. On the relationship beween abundance and distribution of species. Am. Nat. 124, 255–279. https://doi.org/10.1086/284267
Bruelheide, H., Dengler, J., Jiménez-Alfaro, B., Purschke, O., Hennekens, S.M., Chytrý, M., Pillar, V.D., Jansen, F., Kattge, J., Sandel, B., Aubin, I., Biurrun, I., Field, R., Haider, S., Jandt, U., Lenoir, J., Peet, R.K., Peyre, G., Sabatini, F.M., Schmidt, M., Schrodt, F., Winter, M., Aćić, S., Agrillo, E., Alvarez, M., Ambarlı, D., Angelini, P., Apostolova, I., Arfin Khan, M.A.S., Arnst, E., Attorre, F., Baraloto, C., Beckmann, M., Berg, C., Bergeron, Y., Bergmeier, E., Bjorkman, A.D., Bondareva, V., Borchardt, P., Botta-Dukát, Z., Boyle, B., Breen, A., Brisse, H., Byun, C., Cabido, M.R., Casella, L., Cayuela, L., Černý, T., Chepinoga, V., Csiky, J., Curran, M., Ćušterevska, R., Dajić Stevanović, Z., De Bie, E., de Ruffray, P., De Sanctis, M., Dimopoulos, P., Dressler, S., Ejrnæs, R., El-Sheikh, M.A.E.R.M., Enquist, B., Ewald, J., Fagúndez, J., Finckh, M., Font, X., Forey, E., Fotiadis, G., García-Mijangos, I., de Gasper, A.L., Golub, V., Gutierrez, A.G., Hatim, M.Z., He, T., Higuchi, P., Holubová, D., Hölzel, N., Homeier, J., Indreica, A., Işık Gürsoy, D., Jansen, S., Janssen, J., Jedrzejek, B., Jiroušek, M., Jürgens, N., Kącki, Z., Kavgacı, A., Kearsley, E., Kessler, M., Knollová, I., Kolomiychuk, V., Korolyuk, A., Kozhevnikova, M., Kozub, Ł., Krstonošić, D., Kühl, H., Kühn, I., Kuzemko, A., Küzmič, F., Landucci, F., Lee, M.T., Levesley, A., Li, C.F., Liu, H., Lopez-Gonzalez, G., Lysenko, T., Macanović, A., Mahdavi, P., Manning, P., Marcenò, C., Martynenko, V., Mencuccini, M., Minden, V., Moeslund, J.E., Moretti, M., Müller, J. V., Munzinger, J., Niinemets, Ü., Nobis, M., Noroozi, J., Nowak, A., Onyshchenko, V., Overbeck, G.E., Ozinga, W.A., Pauchard, A., Pedashenko, H., Peñuelas, J., Pérez-Haase, A., Peterka, T., Petřík, P., Phillips, O.L., Prokhorov, V., Rašomavičius, V., Revermann, R., Rodwell, J., Ruprecht, E., Rūsiņa, S., Samimi, C., Schaminée, J.H.J., Schmiedel, U., Šibík, J., Šilc, U., Škvorc, Ž., Smyth, A., Sop, T., Sopotlieva, D., Sparrow, B., Stančić, Z., Svenning, J.C., Swacha, G., Tang, Z., Tsiripidis, I., Turtureanu, P.D., Uğurlu, E., Uogintas, D., Valachovič, M., Vanselow, K.A., Vashenyak, Y., Vassilev, K., Vélez-Martin, E., Venanzoni, R., Vibrans, A.C., Violle, C., Virtanen, R., von Wehrden, H.,

Wagner, V., Walker, D.A., Wana, D., Weiher, E., Wesche, K., Whitfeld, T., Willner, W., Wiser, S., Wohlgemuth, T., Yamalov, S., Zizka, G., Zverev, A., 2019. sPlot – A new tool for global vegetation analyses. J. Veg. Sci. 30, 161–186. https://doi.org/10.1111/jvs.12710

Burnham, K.P., Anderson, D.R., 2003. Model selection and multimodel inference: a practical information-theoretic approach. Springer Science and Business Media.

Cameron, R. a D., Triantis, K. a., Parent, C.E., Guilhaumon, F., Alonso, M.R., Ibáñez, M., de Frias Martins, A.M., Ladle, R.J., Whittaker, R.J., 2013. Snails on oceanic islands: Testing the general dynamic model of oceanic island biogeography using linear mixed effect models. J. Biogeogr. 40, 117–130. https://doi.org/10.1111/j.1365-2699.2012.02781.x

Davies, T.J., Kraft, N.J., Salamin, N., Wolkovich, E.M., 2012. Incompletely resolved phylogenetic trees inflate estimates of phylogenetic conservatism. Ecology, 93(2), 242-247.

Dauby, G., 2017. ConR: Computation of Parameters Used in Preliminary Assessment of Conservation Status.

Degnan, J.H., Rosenberg, N. a., 2009. Gene tree discordance, phylogenetic inference and the multispecies coalescent. Trends Ecol. Evol. 24, 332–340. https://doi.org/10.1016/j.tree.2009.01.009

Desgaupes, B., 2018. clusterCrit: Clustering Indices. R package version 1.2.8. https://CRAN.R-project.org/package=clusterCrit

Dimopoulos, P.., Raus, T.., Bergmeier, E.., Constantinidis, T.., Iatrou, G., Kokkini, S.., Strid, A.., Tzanoudakis, D., 2013. Vascular Plants of Greece. Englera.

Dimopoulos, P., Raus, T., Bergmeier, E., Constantinidis, T., Iatrou, G., Kokkini, S., Strid, A., Tzanoudakis, D., 2016. Willdenowia. Willdenowia - Ann. Bot. Gard. Bot. Museum Berlin-Dahlem 46, 301–347.

Dormann, C.F., Elith, J., Bacher, S., Buchmann, C., Carl, G., Carré, G., Marquéz, J.R.G., Gruber, B., Lafourcade, B., Leitão, P.J., Münkemüller, T., Mcclean, C., Osborne, P.E., Reineking, B., Schröder, B., Skidmore, A.K., Zurell, D., Lautenbach, S., 2013. Collinearity: A review of methods to deal with it and a simulation study evaluating their performance. Ecography (Cop.). 36, 027–046. https://doi.org/10.1111/j.1600-0587.2012.07348.x

Elith, J., Kearney, M., Phillips, S., 2010. The art of modelling range-shifting species. Methods Ecol. Evol. 1, 330–342. https://doi.org/10.1111/j.2041-210X.2010.00036.x

Faith, D.P., 1992. Conservation evaluation and phylogenetic diversity. Biological conservation, 61, 1–10.

Ferrier, S., Manion, G., Elith, J., Richardson, K., 2007. Using generalized dissimilarity modelling to analyse and predict patterns of beta diversity in regional biodiversity assessment. Diversity and Distributions, 13, 252–264.

Fitzpatrick, M.C., Sanders, N.J., Normand, S., Svenning, J.-C., Ferrier, S., Gove, A.D., Dunn, R.R., 2013. Environmental and historical imprints on beta diversity: insights from variation in rates of species turnover along gradients. Proceedings. Biological sciences, 280, 20131201.

Fitzpatrick, M.C., Sanders, N.J., Ferrier, S., Longino, J.T., Weiser, M.D., Dunn, R., 2011. Forecasting the future of biodiversity: a test of single- and multi-species models for ants in North America. Ecography (Cop.). 34, 836–847. https://doi.org/10.1111/j.1600-0587.2011.06653.x

Hanson, J.O., 2019. wdpar: Interface to the World Database on Protected Areas. R package version 1.0.0. https://CRAN.R-project.org/package=wdpar

Hengl, T., de Jesus, J.M., Heuvelink, G.B.M., Gonzalez, M.R., Kilibarda, M., Blagotić, A., Shangguan, W., Wright, M.N., Geng, X., Bauer-Marschallinger, B., others, 2017. SoilGrids250m: Global gridded soil information based on machine learning. PLoS One 12, e0169748.

Hijmans, R.J., 2018. raster: Geographic Data Analysis and Modeling.

Hijmans, R.J., Cameron, S.E., Parra, J.L., Jones, P.G., Jarvis, A., 2005. Very high resolution interpolated climate surfaces for global land areas. Int. J. Climatol. 25, 1965–1978.

Januchowski, S. R., Pressey, R. L., VanDerWal, J., Edwards, A., 2010. Characterizing errors in digital elevation models and estimating the financial costs of accuracy. International Journal of Geographical Information Science, 24(9), 1327-1347.

Jarvis, A., Reuter, H.I., Nelson, A., Guevara, E., 2008. Hole-filled SRTM for the globe Version 4.

Jin, Y., Qian, H., 2019. V.PhyloMaker: an R package that can generate very large phylogenies for vascular plants. Ecography, 42, 1353–1359.

Kembel, S.W., Cowan, P.D., Helmus, M.R., Cornwell, W.K., Morlon, H., Ackerly, D.D., Blomberg, S.P., Webb, C.O., 2010. Picante: R tools for integrating phylogenies and ecology. Bioinformatics, 26, 1463–1464.

Laffan, S.W., Lubarsky, E., Rosauer, D.F., 2010. Biodiverse, a tool for the spatial analysis of biological and related diversity. Ecography (Cop.). 33, 643–647. https://doi.org/10.1111/j.1600-0587.2010.06237.x

Lawler, J.J., White, D., Neilson, R.P., Blaustein, A.R., 2006. Predicting climate-induced range shifts: model differences and model reliability. Global Change Biology, 12(8), 1568-1584.

Legendre, P., 2014. Interpreting the replacement and richness difference components of beta diversity. Glob. Ecol. Biogeogr. 23, 1324–1334. https://doi.org/10.1111/geb.12207

Le Saout, S., Hoffmann, M., Shi, Y., Hughes, A., Bernard, C., Brooks, T.M., Bertzky, B., Butchart, S.H.M., Stuart, S.N., Badman, T., Rodrigues, A.S.L., 2013. Protected areas and effective biodiversity conservation. Science (80-. ). https://doi.org/10.1126/science.1239268

Levins, R., 1968. Evolution in changing environments: some theoretical explorations (No. 2). Princeton University Press.

Lüdecke, D., 2017. sjstats: Statistical Functions for Regression Models.

Maechler, M., Rousseeuw, P., Struyf, A., Hubert, M. and Hornik, K., 2018. cluster: Cluster Analysis Basics and Extensions. R package version 2.0.7-1.

Maitner, B.S., Boyle, B., Casler, N., Condit, R., Donoghue, J., Durán, S.M., Guaderrama, D., Hinchliff, C.E., Jørgensen, P.M., Kraft, N.J.B., McGill, B., Merow, C., Morueta-Holme, N., Peet, R.K., Sandel, B., Schildhauer, M., Smith, S.A., Svenning, J.-C., Thiers, B., Violle, C., Wiser, S., Enquist, B.J., 2018. The <scp>bien r</scp> package: A tool to access the Botanical Information and Ecology Network (BIEN) database. Methods Ecol. Evol. 9, 373–379. https://doi.org/10.1111/2041-210X.12861

Manion, G., Lisk, M., Ferrier, S., Nieto-Lugilde, D., Mokany, K., Fitzpatrick, M.C., 2018. gdm: Generalized Dissimilarity Modeling. R package version 1.3.7.

Mazel, F., Renaud, J., Guilhaumon, F., Mouillot, D., Gravel, D., Thuiller, W., 2015. Mammalian phylogenetic diversity–area relationships at a continental scale. Ecology, 96(10), 2814-2822.

McKnight, M.W., White, P.S., McDonald, R.I., Lamoreux, J.F., Sechrest, W., Ridgely, R.S., Stuart, S.N., 2007. Putting beta-diversity on the map: broad-scale

congruence and coincidence in the extremes. PLoS biology, 5(10), e272.
McSweeney, C.F., Jones, R.G., Lee, R.W., Rowell, D.P., 2015. Selecting CMIP5 GCMs for downscaling over multiple regions. Clim. Dyn. 44, 3237–3260. https://doi.org/10.1007/s00382-014-2418-8
Menéndez-Guerrero, P.A., Green, D.M., Davies, T.J., 2019. Climate change and the future restructuring of Neotropical anuran biodiversity. Ecography (Cop.). ecog.04510. https://doi.org/10.1111/ecog.04510
Mishler, B.D., Knerr, N., González-Orozco, C.E., Thornhill, A.H., Laffan, S.W., Miller, J.T., 2014. Phylogenetic measures of biodiversity and neo-and paleo-endemism in Australian acacia. Nat. Commun. 5. https://doi.org/10.1038/ncomms5473
Naimi, B., Hamm, N.A.S., Groen, T.A., Skidmore, A.K., Toxopeus, A.G., 2014. Where is positional uncertainty a problem for species distribution modelling? Ecography (Cop.). 37, 191–203. https://doi.org/10.1111/j.1600-0587.2013.00205.x
Nowosad, J., Stepinski, T.F., 2018. Spatial association between regionalizations using the information-theoretical *V* -measure. Int. J. Geogr. Inf. Sci. 32, 2386–2401. https://doi.org/10.1080/13658816.2018.1511794
Pebesma, E., 2018. Simple Features for R: Standardized Support for Spatial Vector Data. The R Journal 10.
(1), 439-446, https://doi.org/10.32614/RJ-2018-009Raes, N., ter Steege, H., 2007. A null-model for significance testing of presence-only species distribution models. Ecography (Cop.). 30, 727–736.
Robertson, M.P., Visser, V., Hui, C., 2016. Biogeo: an R package for assessing and improving data quality of occurrence record datasets. Ecography (Cop.). 39, 394–401.
Rosauer, D., Laffan, S.W., Crisp, M.D., Donnellan, S.C., Cook, L.G., 2009. Phylogenetic endemism: a new approach for identifying geographical concentrations of evolutionary history. Molecular Ecology, 18(19), 4061-4072.
Rousseeuw, P.J., 1987. Silhouettes: a graphical aid to the interpretation and validation of cluster analysis, Journal of Computational and Applied Mathematics.
Smith, S.A., Brown, J.W., 2018. Constructing a broadly inclusive seed plant phylogeny. American Journal of Botany, 105(3), 302-314.
Stévart, T., Dauby, G., Lowry, P.P., Blach-Overgaard, A., Droissart, V., Harris, D.J., Mackinder, B.A., Schatz, G.E., Sonké, B., Sosef, M.S.M., Svenning, J.-C., Wieringa, J.J., Couvreur, T.L.P., 2019. A third of the tropical African flora is potentially threatened with extinction. Sci. Adv. 5, eaax9444. https://doi.org/10.1126/sciadv.aax9444
Strid, A., 2016. Atlas of the Aegean flora. Botanic Garden and Botanical Museum Berlin-Freie Universität Berlin, Berlin.
Thuiller, W., Lafourcade, B., Engler, R., Araújo, M.B., 2009. BIOMOD - A platform for ensemble forecasting of species distributions. Ecography (Cop.). 32, 369–373. https://doi.org/10.1111/j.1600-0587.2008.05742.x
Title, P.O., Bemmels, J.B., 2017. ENVIREM: An expanded set of bioclimatic and topographic variables increases flexibility and improves performance of ecological niche modeling. Ecography (Cop.).
Tsirogiannis, K., Sandel, B., 2017. PhyloMeasures: Fast and exact algorithms for computing phylogenetic biodiversity measures. *R package version*, *2*.
Valavi, R., Elith, J., Lahoz-Monfort, J., Guillera-Arroita, G., 2018. blockCV: Spatial and environmental blocking for k-fold cross-validation.
VanDerWal, J., Falconi, L., Januchowski, S., Shoo, L., Storlie, C., 2014. SDMTools:

Species Distribution Modelling Tools: Tools for processing data associated with species distribution modelling exercises. *R package version*, *1*, 1-221.

van Proosdij, A.S.J., Sosef, M.S.M., Wieringa, J.J., Raes, N., 2016. Minimum required number of specimen records to develop accurate species distribution models. Ecography (Cop.). 39, 542–552. https://doi.org/10.1111/ecog.01509

Wood, S.N., 2017. Generalized additive models: an introduction with R. Chapman and Hall/CRC.

Xu, J., Su, G., Xiong, Y., Akasaka, M., García Molinos, J., Matsuzaki, S. ichiro S., Zhang, M., 2015. Complimentary analysis of metacommunity nestedness and diversity partitioning highlights the need for a holistic conservation strategy for highland lake fish assemblages. Glob. Ecol. Conserv. 3, 288–296. https://doi.org/10.1016/j.gecco.2014.12.004

Zizka, A., Antonelli, A., Silvestro, D., 2020. sampbias, a method for quantifying geographic sampling biases in species distribution data. bioRxiv, DOI: https://doi.org/10.1101/2020.01.13.903757
