## Appendix B for "Management implications based on diversity patterns under climate change scenarios in a continental island biodiversity hotspot"

**Appendix B** Supplementary Tables and Figures

#### Supplementary tables

**Table S2.** Median range contraction values for every Global Circulation Models (GCMs) and Representative Concentration Pathways (RCPs) included in the analyses.

| GCM/RCP | Median |
|---|---|
| BCC 2.6 | 96.7 |
| BCC 8.5 | 100.0 |
| CCSM4 2.6 | 68.1 |
| CCSM4 8.5 | 100.0 |
| HadGEM2 2.6 | 94.4 |
| HadGEM2 8.5 | 100.0 |

**Table S3.** Median altitude and pH for sites having higher or lower than expected phylogenetic alpha diversity (PD,) as well as for the not-significant sites. NS: not-significant. Over: sites with higher than expected PD. Under: sites with lower than expected PD.

| Type | Altitude (m a.s.l.) | pH |
|---|---|---|
| Over | 985 | 7.06 |
| Under | 865 | 7.05 |
| NS | 446 | 7.12 |

**Table S4.** Median altitude for the different types of endemism centres as well as for the not-significant sites. NS: not-significant.

| Type | Altitude (m a.s.l.) |
|---|---|
| Mixed | 811 |
| Neo | 652 |
| Paleo | 930 |
| Super | 82 |
| NS | 562 |

**Table S5.** Median altitude for the different types of endemism centres for the present, as well as for all Global Circulation Models (GCMs) and Representative Concentration Pathways (RCPs) included in the analyses. * denotes a p-value < 0.001 in the Kruskal-Wallis ANOVA.

| GCM/RCP | Mixed | Neo | Paleo | Super |
|---|---|---|---|---|
| Current | 811 | 652 | 930 | 82 |
| BCC 2.6 | 191* | 327 | 108* | 156 |
| BCC 8.5 | 332* | 562 | 299* | 239 |
| CCSM4 2.6 | 307* | 154* | 547 | 1458 |
| CCSM4 8.5 | 493* | 448 | 276* | 344 |
| HadGEM2 2.6 | 402* | 160* | 161* | 197 |
| HadGEM2 8.5 | 395 | 78* | 549 | 284 |

**Table S6.** Best spatial autoregressive error models ($SAR_{err}$) for the relationships among phylogenetic endemism (PE), relative phylogenetic endemism (RPE) and the predictor variables. $GR^2$: Gelkerke pseudo R-squared. AICc: Akaike Information criterion corrected for small samples. Asterisks denote: * p < 0.05, ** p < 0.01, *** p < 0.001. Alt: Altitude. CS: Climate stability. MDR: Mean diurnal range.

| Response | Predictor | Coefficients | $GR^2$ | AICc |
|---|---|---|---|---|
| PE | Alt | 1.44*** | 16.1 | 11612.6 |
| | MDR | 0.44** | | |
| | pH | 0.40** | | |
| | CS | 0.33** | | |
| RPE | Alt | 1.01* | 5.9 | 4455.3 |
| | MDR | -0.02 | | |
| | pH | -0.03 | | |
| | CS | 0.03 | | |

**Table S7.** Results for BCC 2.6 GCM/RCP combination of the best (i.e., full) generalized additive model relating change in $SIE_C$ beta-diversity to change in species richness (ΔSR), average level of ecological generalism (ΔEG) and phylogenetic diversity (ΔPD), as well as elevation. Variables were standardized [i.e., (value-mean/standard deviation)] prior to analysis. AIC: Akaike Information Criterion. AICc: Akaike Information criterion corrected for small samples. df: Degrees of freedom. F: F-values. logLik: log-likehood. $R^2_{adj}$: adjusted $R^2$. The full model was the only model with ΔAICc < 2. Asterisks denote: * p < 0.05, ** p < 0.01, *** p < 0.001.

| Model | Predictors | df | AIC | AICc | logLik | $R^2_{adj.}$ | Deviance (%) | F | p |
|---|---|---|---|---|---|---|---|---|---|
| BCC 2.6 | - | 88.4 | 20067 | 20068 | -9945 | 0.710 | 71.2 | - | - |
| | ΔSR | | | | | | | 33.9 | *** |
| | ΔPD | | | | | | | 2.7 | *** |
| | ΔEG | | | | | | | 515.5 | *** |
| | Elevation | | | | | | | 21.2 | *** |
| BCC 8.5 | - | 102 | 1817 | 1819 | -806 | 0.933 | 93.3 | - | - |
| | ΔSR | | | | | | | 77.4 | *** |
| | ΔPD | | | | | | | 30.2 | *** |
| | ΔEG | | | | | | | 2450.1 | *** |
| | Elevation | | | | | | | 22.1 | *** |
| CCSM4 2.6 | - | 82.8 | 11866 | 11867 | -5850 | 0.850 | 85.1 | - | - |
| | ΔSR | | | | | | | 36.4 | *** |
| | ΔPD | | | | | | | 22.4 | *** |
| | ΔEG | | | | | | | 1634.8 | *** |
| | Elevation | | | | | | | 19.8 | *** |
| CCSM4 8.5 | - | 102 | 1817 | 1819 | -806 | 0.933 | 93.3 | - | - |
| | ΔSR | | | | | | | 77.4 | *** |
| | ΔPD | | | | | | | 30.2 | *** |
| | ΔEG | | | | | | | 2450.1 | *** |
| | Elevation | | | | | | | 22.1 | *** |
| HadGEM2 2.6 | - | 82.7 | 8966 | 8967 | | | | - | - |
| | ΔSR | | | | | | | 85.3 | *** |
| | ΔPD | | | | | | | 4.05 | *** |
| | ΔEG | | | | | | | 1695.0 | *** |
| | Elevation | | | | | | | 93.9 | *** |
| HadGEM2 8.5 | - | 91.7 | -6566 | -6564 | 3375 | 0.966 | 96.6 | - | - |
| | ΔSR | | | | | | | 195.9 | *** |
| | ΔPD | | | | | | | 23.1 | *** |
| | ΔEG | | | | | | | 5620.5 | *** |
| | Elevation | | | | | | | 2.5 | *** |

**Table S8.** Number of biogeographical regions (BR), Silhouette index (SI) for the k-means and CLARA unsupervised clustering algorithms and the V-measure index for the present and every Global Circulation Model (GCM) and Representative Concentration Pathway (RCP) combination.

| GCM/RCP | BR | SI k-means | SI CLARA | V-measure |
|---|---|---|---|---|
| Present | 14 | 0.430 | 0.410 | 1.000 |
| BCC 2.6 | 7 | 0.414 | 0.392 | 0.714 |
| BCC 8.5 | 4 | 0.475 | 0.432 | 0.533 |
| CCSM4 2.6 | 16 | 0.421 | 0.400 | 0.769 |
| CCSM4 8.5 | 12 | 0.499 | 0.412 | 0.503 |
| HADGEM2 2.6 | 9 | 0.429 | 0.377 | 0.683 |
| HADGEM2 8.5 | 2 | 0.431 | 0.430 | 0.338 |

**Table S9.** Extent ($km^2$), species richness (SR) and the standardized effect scores of phylogenetic alpha diversity ($SES_{PD}$) of the areas identified as climate refugia in Crete for the present and every Global Circulation Model (GCM) and Representative Concentration Pathway (RCP) combination. A: Asterousia. B: East. C: East-Central. D: Eastern. E: West. F: Western.

| GCM/RCP | Variable | A | B | C | D | E | F |
|---|---|---|---|---|---|---|---|
| Present | Area | 2.1 | 54.8 | 167.8 | 31.6 | 189.2 | 39.2 |
| | SR | 4 | 46 | 82 | 8 | 121 | 14 |
| | $SES_{PD}$ | -1.05 | 0.39 | 2.07 | -1.80 | 1.02 | -2.13 |
| BCC 2.6 | Area | 2.1 | 54.8 | 167.8 | 31.6 | 189.2 | 39.2 |
| | SR | 4 | 8 | 21 | 3 | 109 | 77 |
| | $SES_{PD}$ | 0.64 | 1.52 | -0.02 | 0.75 | -0.77 | 0.33 |
| BCC 8.5 | Area | 2.1 | 54.8 | 167.8 | 31.6 | 189.2 | 39.2 |
| | SR | 1 | 1 | 19 | 1 | 5 | 8 |
| | $SES_{PD}$ | 0 | 0 | -0.76 | 0 | 0.98 | -0.83 |
| CCSM4 2.6 | Area | 2.1 | 54.8 | 167.8 | 31.6 | 189.2 | 39.2 |
| | SR | 11 | 51 | 70 | 12 | 119 | 96 |
| | $SES_{PD}$ | -2.31 | -0.22 | -0.95 | -1.82 | -1.00 | -0.72 |
| CCSM4 8.5 | Area | 2.1 | 54.8 | 167.8 | 31.6 | 189.2 | 39.2 |
| | SR | 1 | 1 | 6 | 1 | 7 | 11 |
| | $SES_{PD}$ | 0 | 0 | -0.22 | 0 | -0.64 | -1.46 |
| HadGEM2 2.6 | Area | 2.1 | 54.8 | 167.8 | 31.6 | 189.2 | 39.2 |
| | SR | 14 | 38 | 55 | 23 | 82 | 70 |
| | $SES_{PD}$ | -1.61 | -0.78 | -1.56 | -0.71 | -1.07 | -1.36 |
| HadGEM2 8.5 | Area | 2.1 | 54.8 | 167.8 | 31.6 | 189.2 | 39.2 |
| | SR | 1 | 1 | 1 | 1 | 1 | 1 |
| | $SES_{PD}$ | 0 | 0 | 0 | 0 | 0 | 0 |

**Table S10.** Percent overlap (%) between the protected areas (PA) network in Crete, the climate refugia (CR) recognised in Crete and the endemism centres detected by the Categorical Analyses of Neo- and Paleo-Endemism (CANAPE). GCM: Global Circulation Model. RCP: Representative Concentration Pathway. The extent (in $km^2$) of each CANAPE category for every GCM/RCP combination is also presented.

| Type | GCM/RCP | Mixed | Neo | Paleo | Super |
|---|---|---|---|---|---|
| PA | Present | 63 | 52 | 65 | 60 |
| | BCC 2.6 | 23 | 45 | 44 | 22 |
| | BCC 8.5 | 18 | 52 | 30 | 43 |
| | CCSM4 2.6 | 47 | 28 | 65 | 61 |
| | CCSM4 8.5 | 23 | 20 | 13 | 0 |
| | HadGEM2 2.6 | 45 | 38 | 42 | 40 |
| | HadGEM2 8.5 | 22 | 0 | 0 | 0 |
| CR | Present | 22 | 0 | 4.9 | 0.7 |
| | BCC 2.6 | 6.7 | 13.6 | 3.1 | 0 |
| | BCC 8.5 | 0.01 | 14.3 | 1.6 | 0 |
| | CCSM4 2.6 | 9.4 | 4.5 | 23.2 | 29.5 |
| | CCSM4 8.5 | 0 | 0 | 0 | 0 |
| | HadGEM2 2.6 | 13.2 | 6.9 | 7.5 | 20.8 |
| | HadGEM2 8.5 | 0 | 0 | 0 | 0 |
| Extent | Present | 109.4 | 14.7 | 18.2 | 3.5 |
| | BCC 2.6 | 555 | 15 | 22 | 26 |
| | BCC 8.5 | 133 | 15 | 89 | 15 |
| | CCSM4 2.6 | 446 | 123 | 85 | 43 |
| | CCSM4 8.5 | 61.8 | 3.5 | 5.6 | 42 |
| | HadGEM2 2.6 | 219 | 20 | 47 | 34 |
| | HadGEM2 8.5 | 6.6 | 2.8 | 0.7 | 2.1 |

**Table S11.** Percent overlap (%) between the protected areas (PA) network and the climate refugia (CR) recognised in Crete. The extent (in $km^2$) of each climate refugium is also presented.

| Climate refugium | Extent | Percent overlap (%) |
|---|---|---|
| Asterousia | 2.1 | 33.33 |
| East | 54.8 | 48.70 |
| East-Central | 167.8 | 69.90 |
| Eastern | 31.6 | 13.33 |
| West | 189.2 | 97.00 |
| Western | 39.2 | 35.70 |

**Table S12.** Median percent overlap (%) between the protected areas (PA) network in Crete, the climate refugia (CR) recognised in Crete and the endemism centres detected by the Categorical Analyses of Neo- and Paleo-Endemism (CANAPE) for the present, as well as for the future climate conditions (averaged for all Global Circulation Models and Representative Concentration Pathways).

| Type | Median | | Current state | |
|---|---|---|---|---|
| | PA | CR | PA | CR |
| Mixed | 23.00 | 3.36 | 63.00 | 22.00 |
| Neo | 33.00 | 5.70 | 52.00 | 0.00 |
| Paleo | 36.00 | 2.35 | 65.00 | 4.90 |
| Super | 31.00 | 0.00 | 60.00 | 0.70 |

### Supplementary Figures

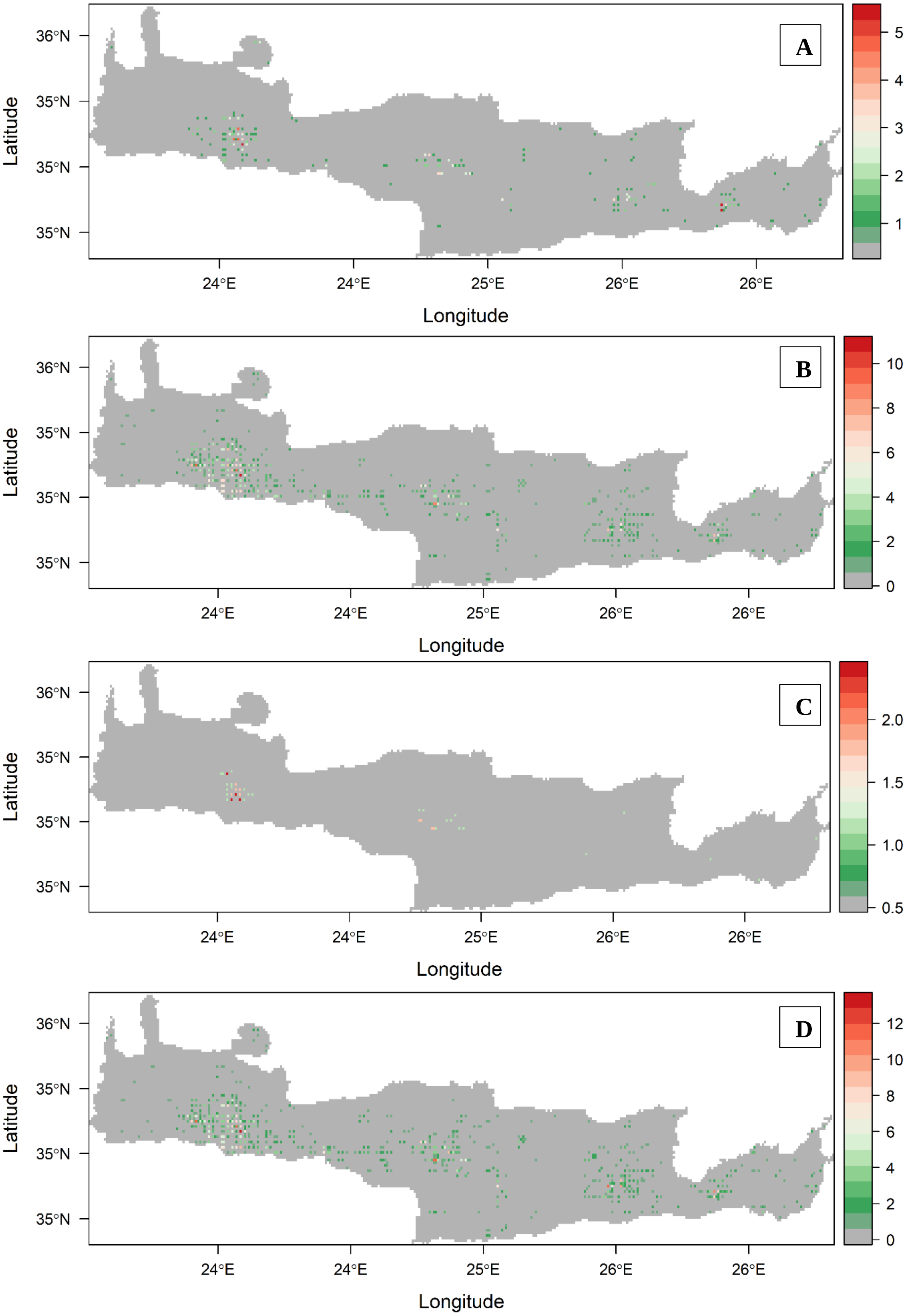

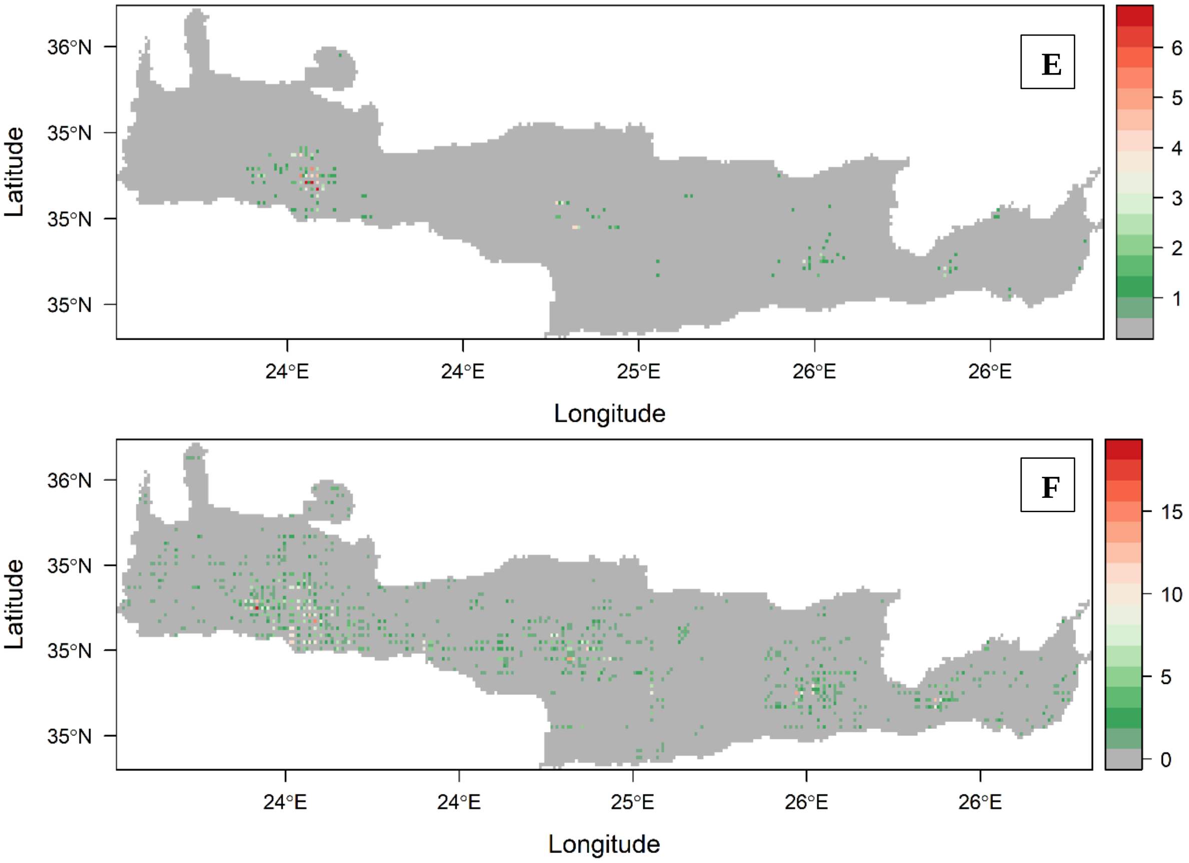


**Figure S1.** Number of species predicted to become extinct under (A) BCC 2.6, (B) BCC 8.5, (C) CCSM4 2.6, (D) CCSM4 8.5, (E) HadGEM2 2.6 and (F) HadGEM2 8.5 Global Circulation Model and Representative Concentration Pathway.

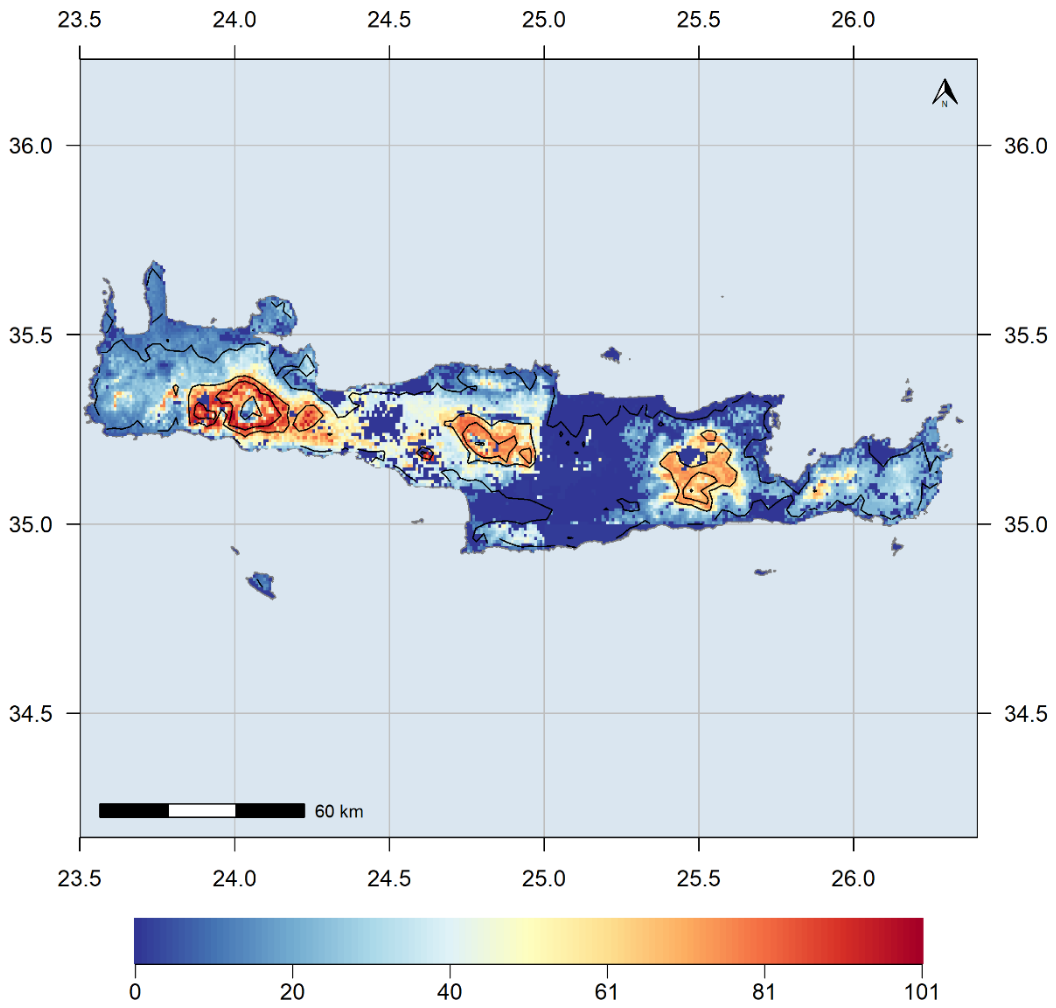


**Figure S2.** Species richness map for the $SIE_C$ for the present.

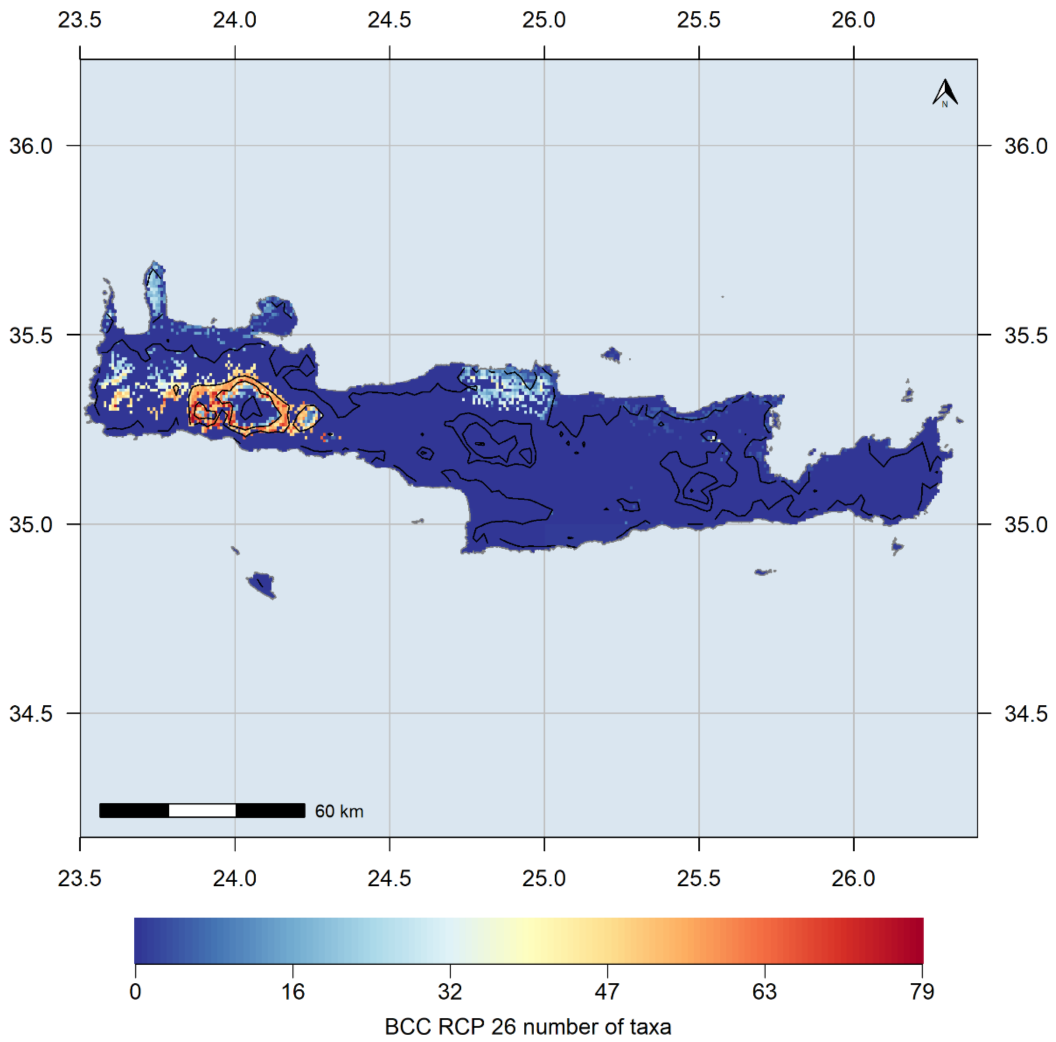


**Figure S3.** Species richness map for the $SIE_C$ for the BCC 2.6 GCM/RCP combination.

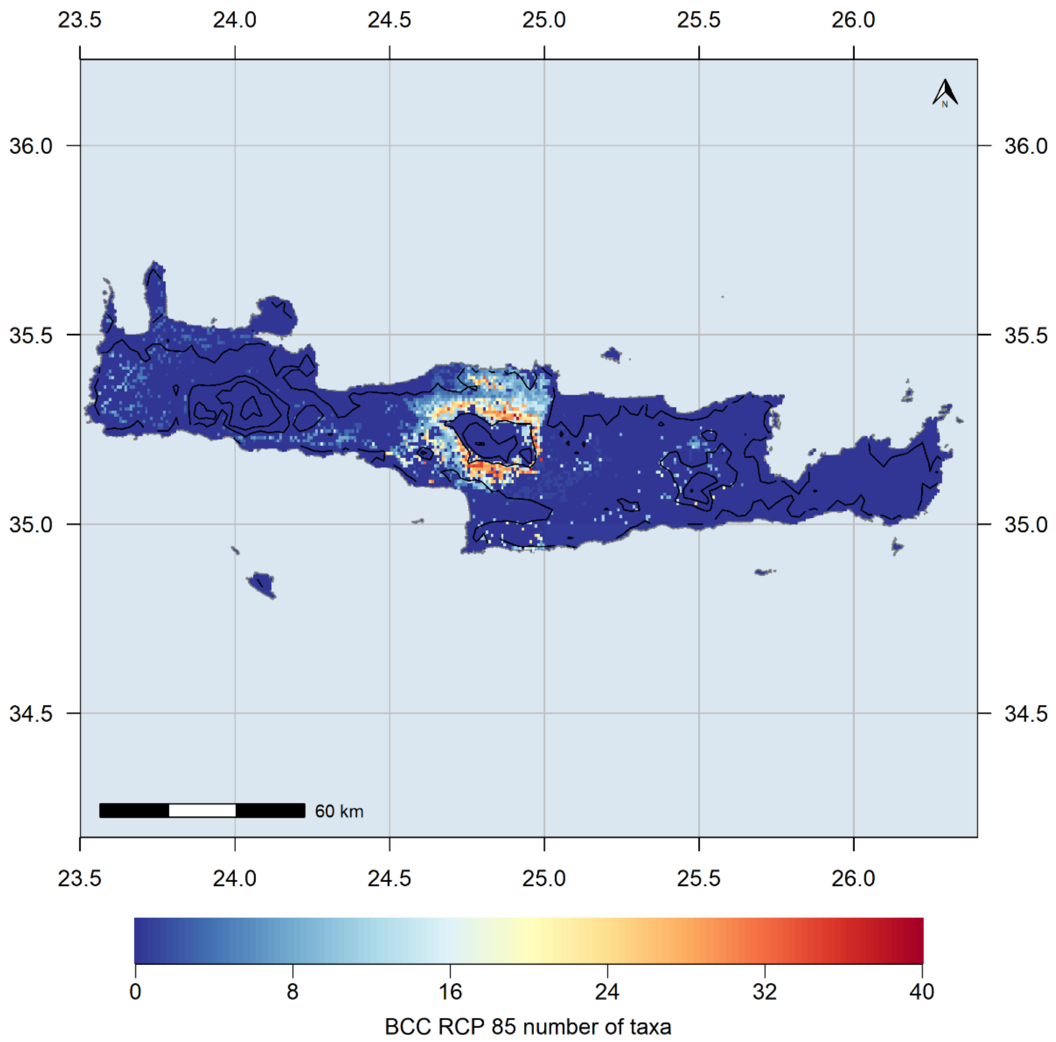


**Figure S4.** Species richness map for the $SIE_C$ for the BCC 8.5 GCM/RCP combination

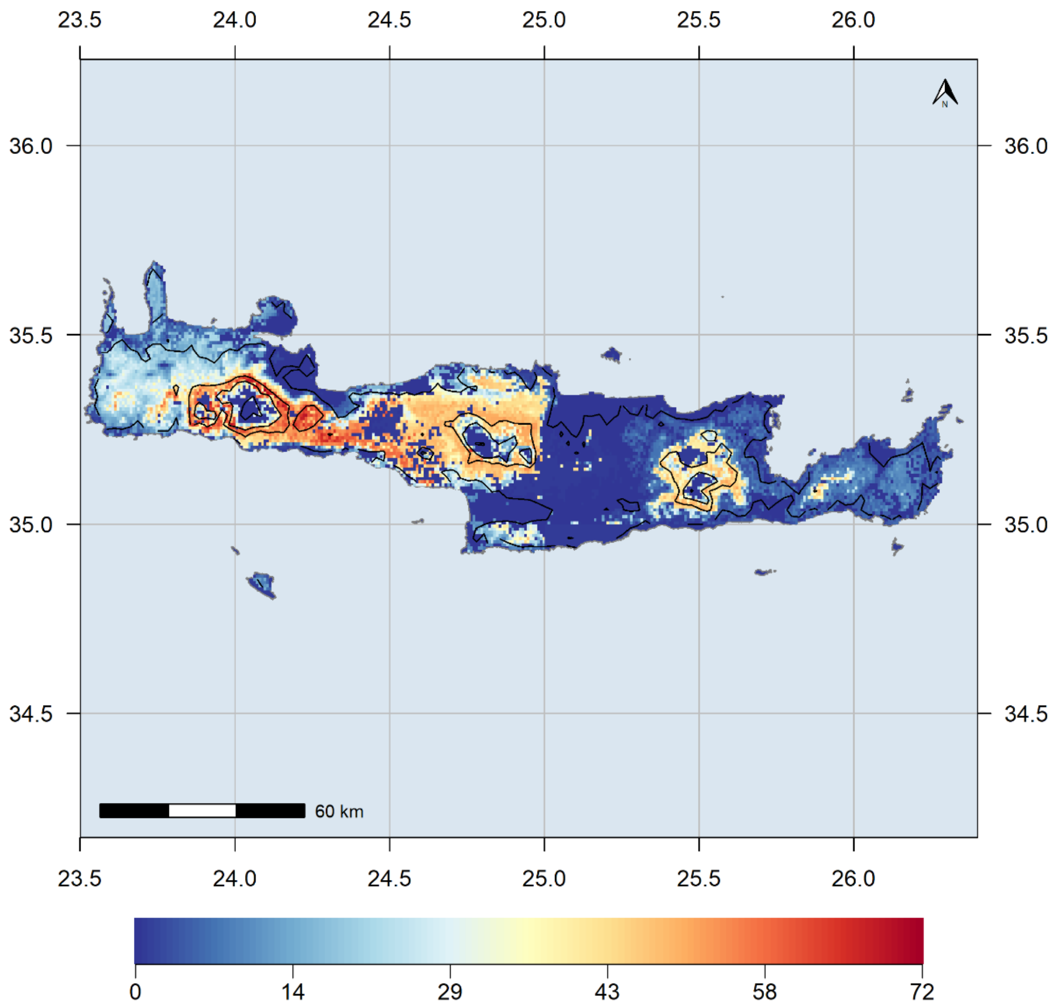


**Figure S5.** Species richness map for the $SIE_C$ for the CCSM4 2.6 GCM/RCP combination

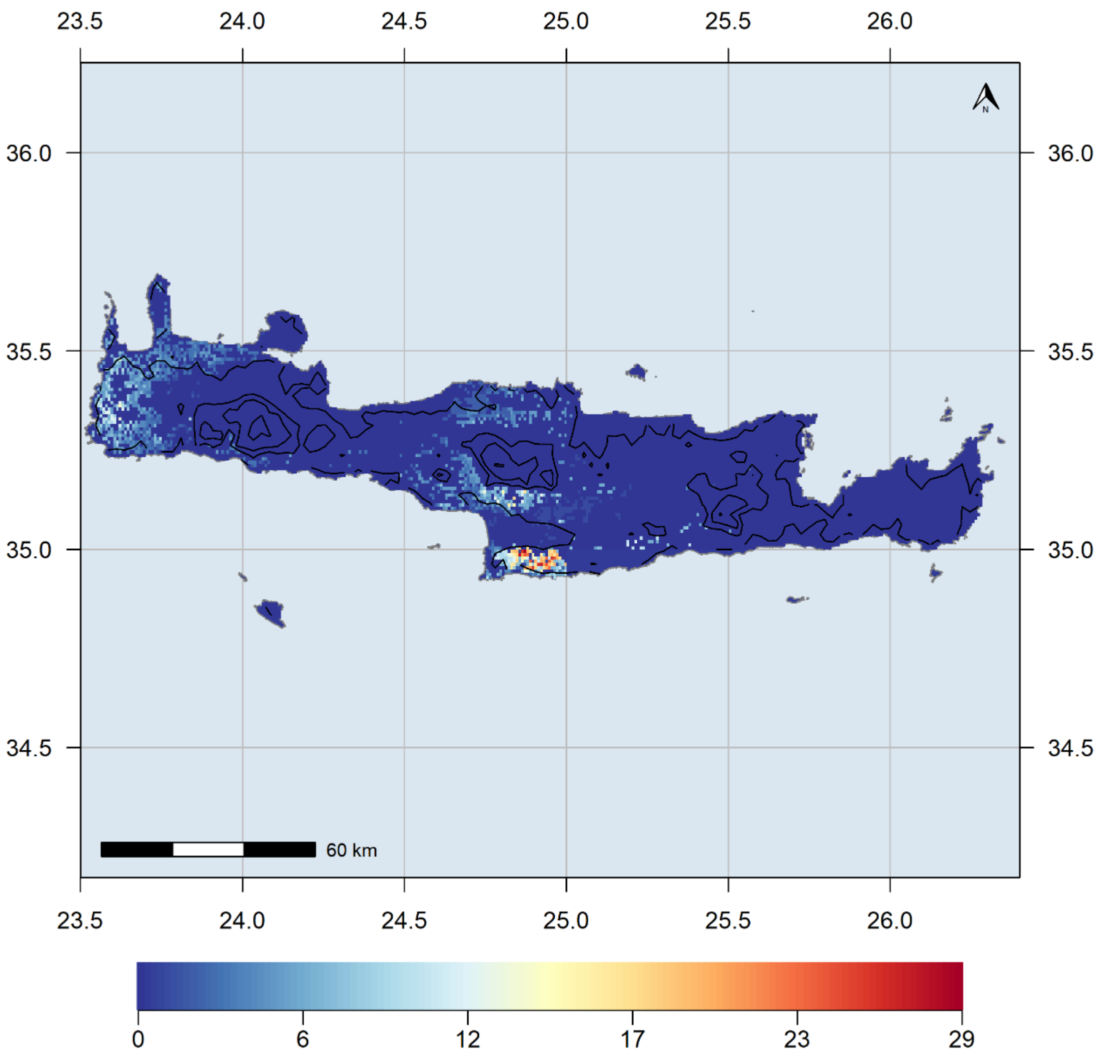


**Figure S6.** Species richness map for the SIE$_C$ for the CCSM4 8.5 GCM/RCP combination

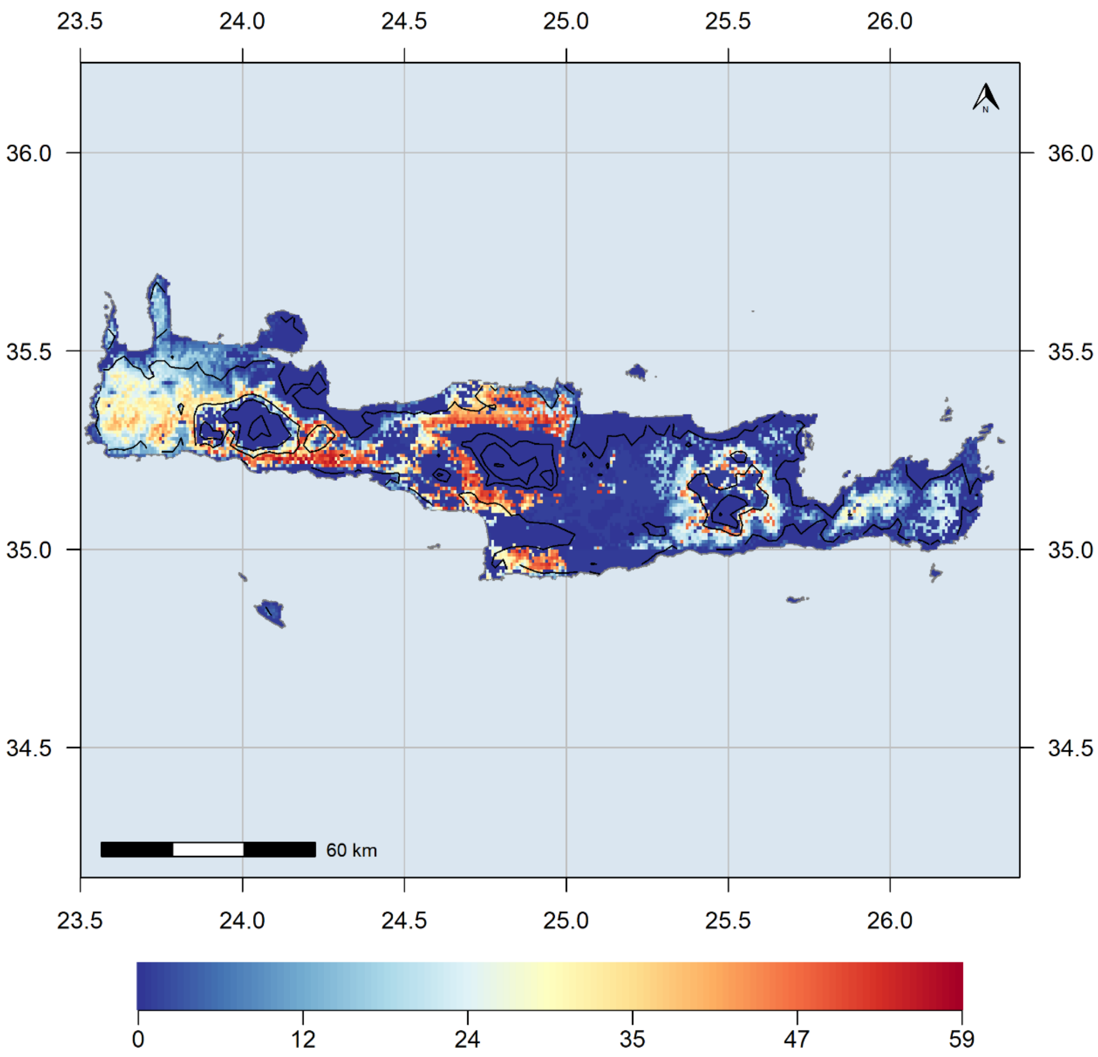


**Figure S7.** Species richness map for the $SIE_C$ for the HadGEM2 2.6 GCM/RCP combination

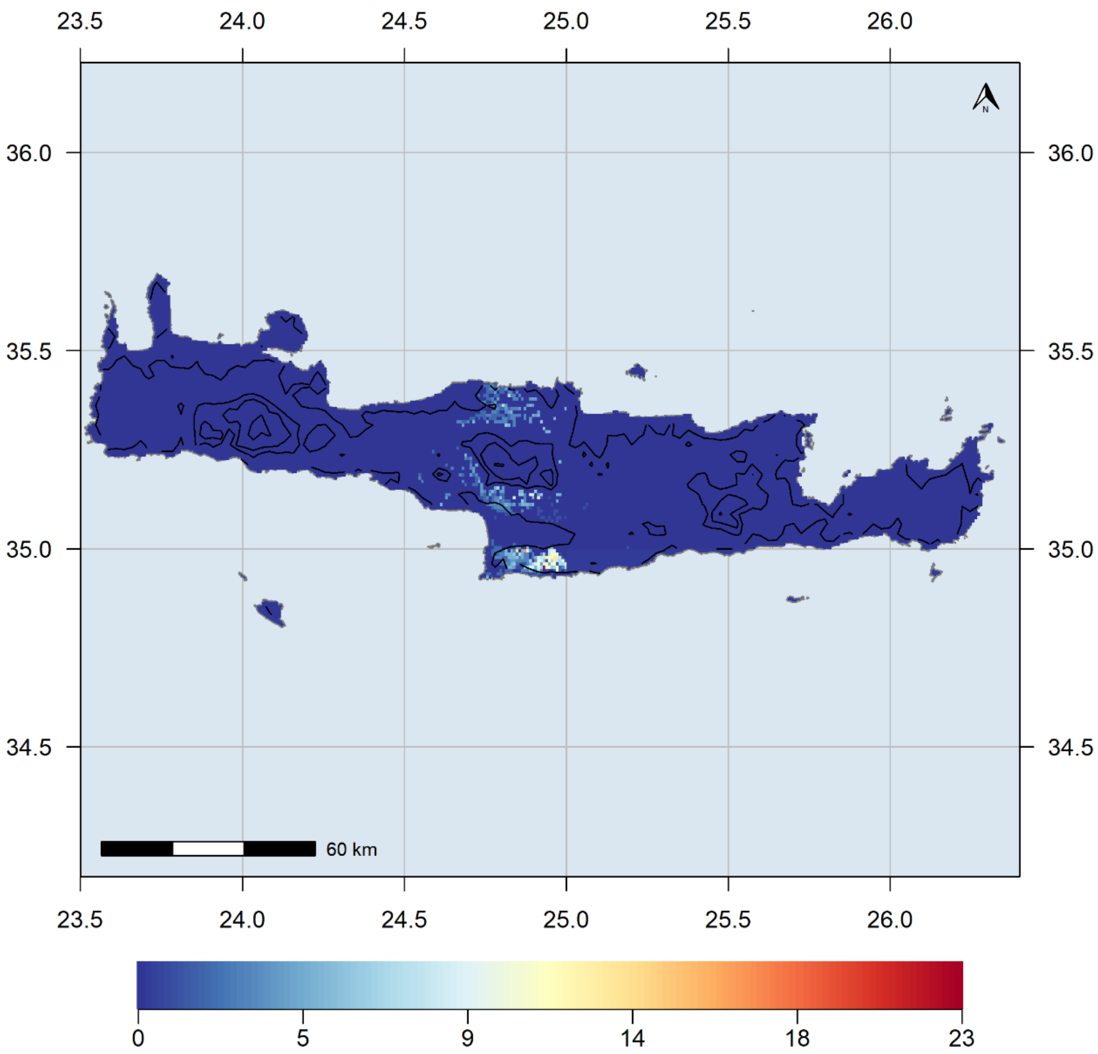


**Figure S8.** Species richness map for the $SIE_C$ for the HadGEM2 8.5 GCM/RCP combination

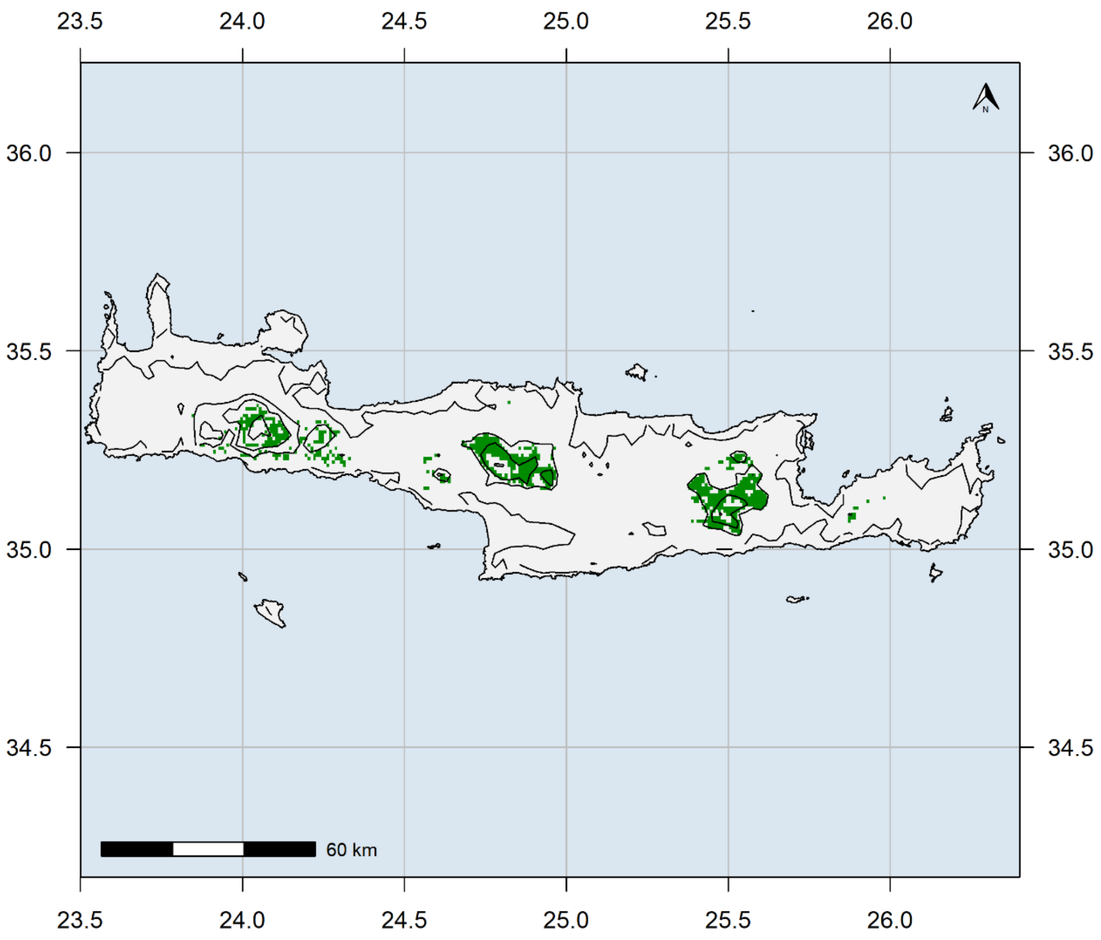


**Figure S9.** Absolute difference between current hotspots and the BCC 2.6 GCM/RCP combination.

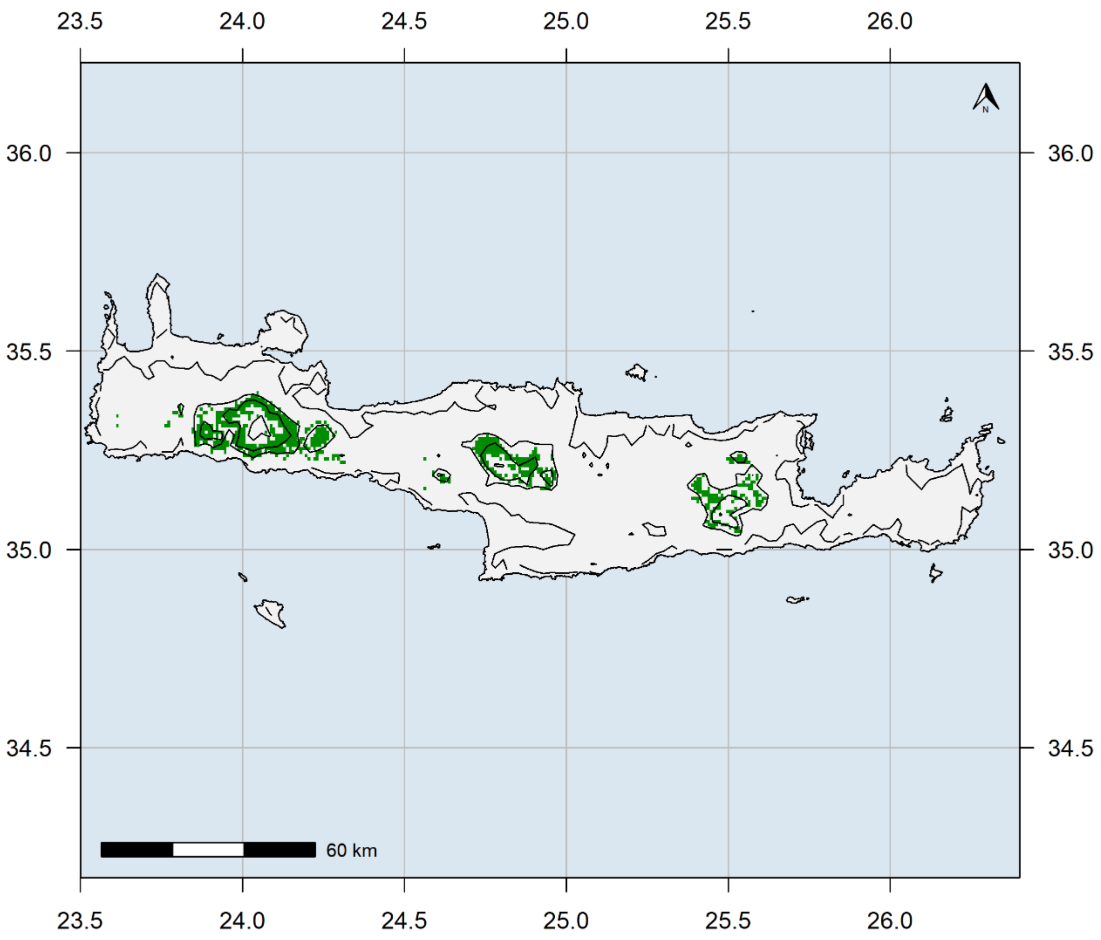


**Figure S10.** Absolute difference between current hotspots and the BCC 8.5 GCM/RCP combination.

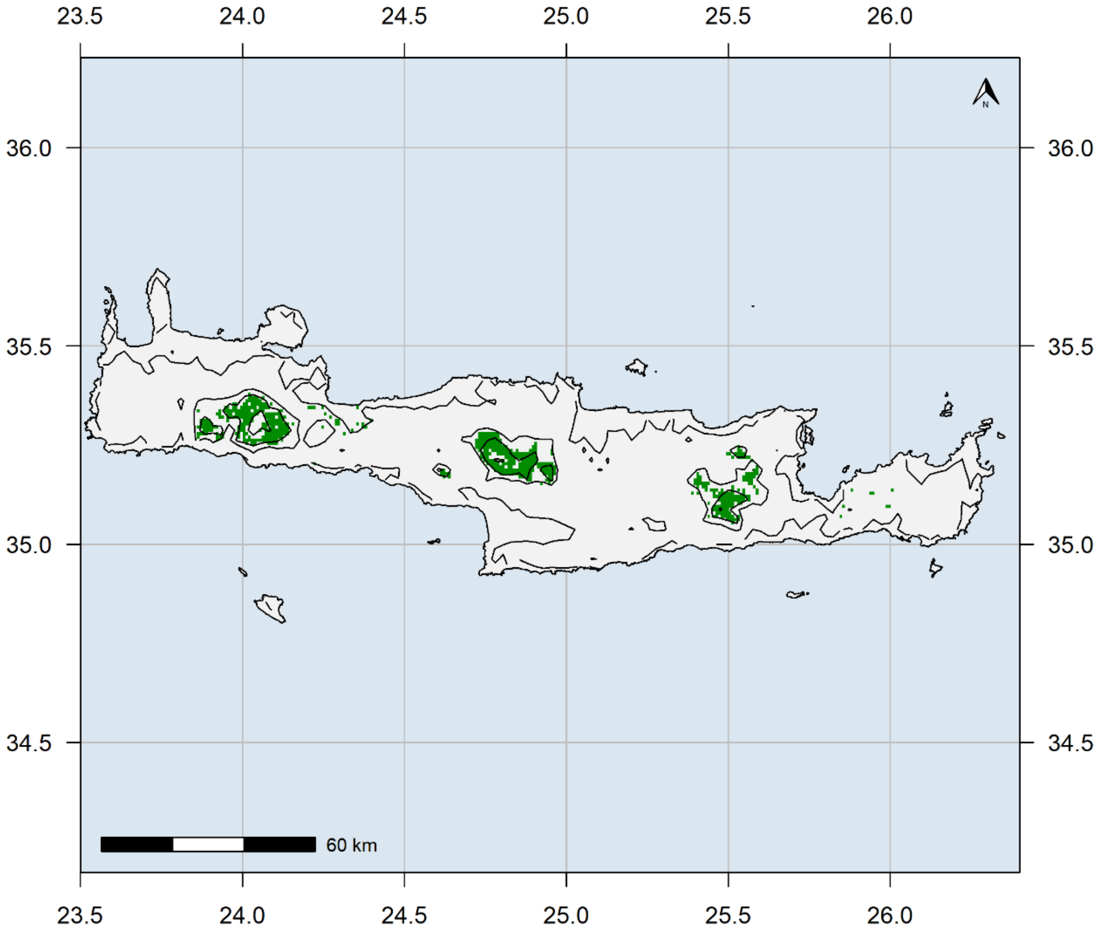


**Figure S11.** Absolute difference between current hotspots and the CCSM4 2.6 GCM/RCP combination.

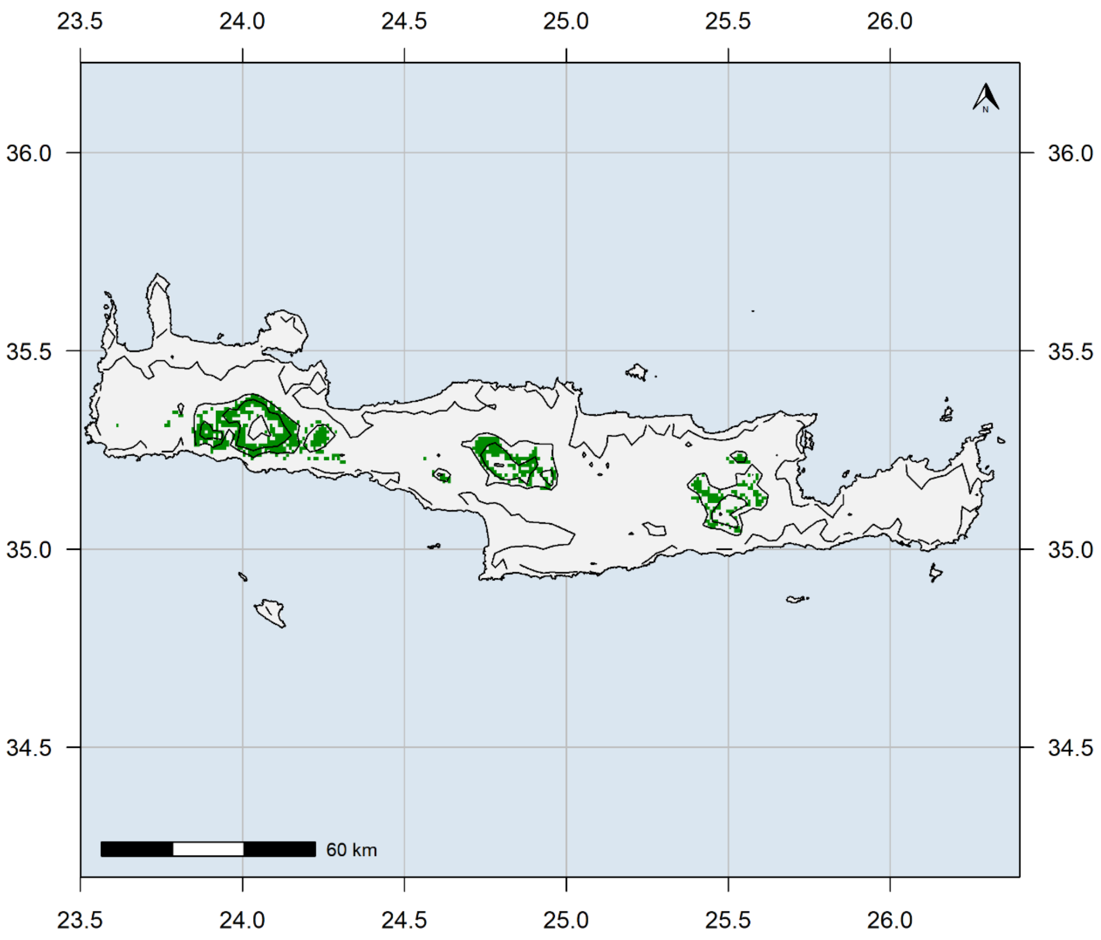


**Figure S12.** Absolute difference between current hotspots and the CCSM4 8.5 GCM/RCP combination.

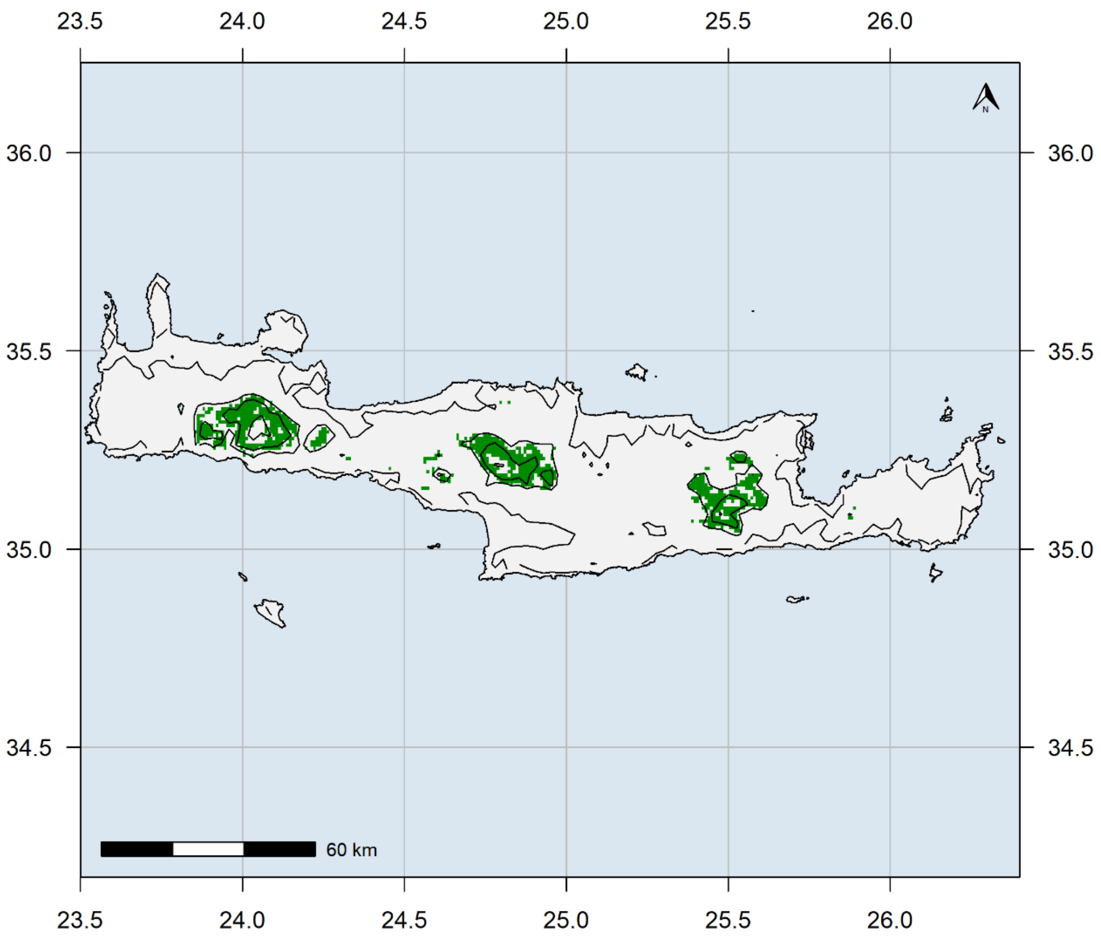


**Figure S13.** Absolute difference between current hotspots and the HadGEM2 2.6 GCM/RCP combination.

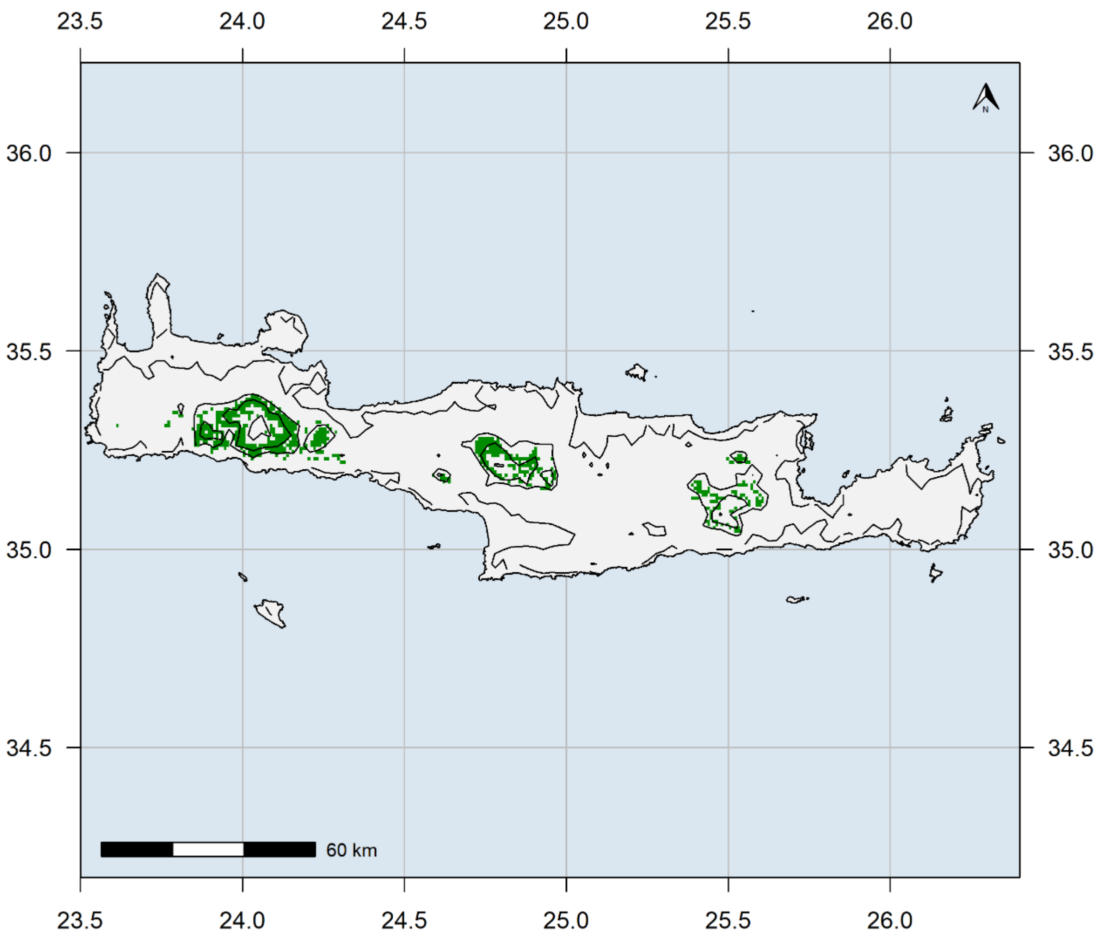


**Figure S14.** Absolute difference between current hotspots and the HadGEM2 8.5 GCM/RCP combination.

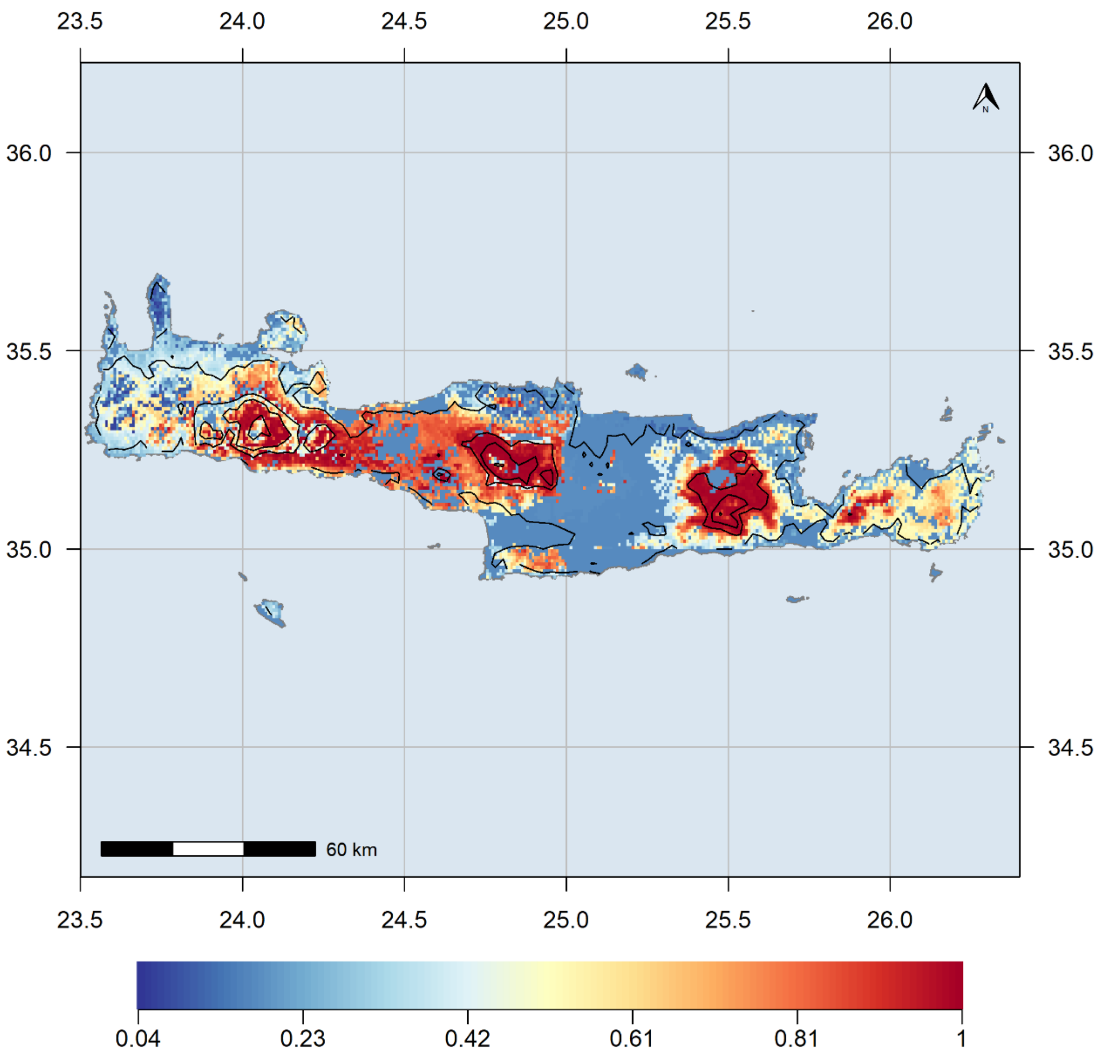


**Figure S15.** Relative difference between current hotspots and the BCC 2.6 GCM/RCP combination.

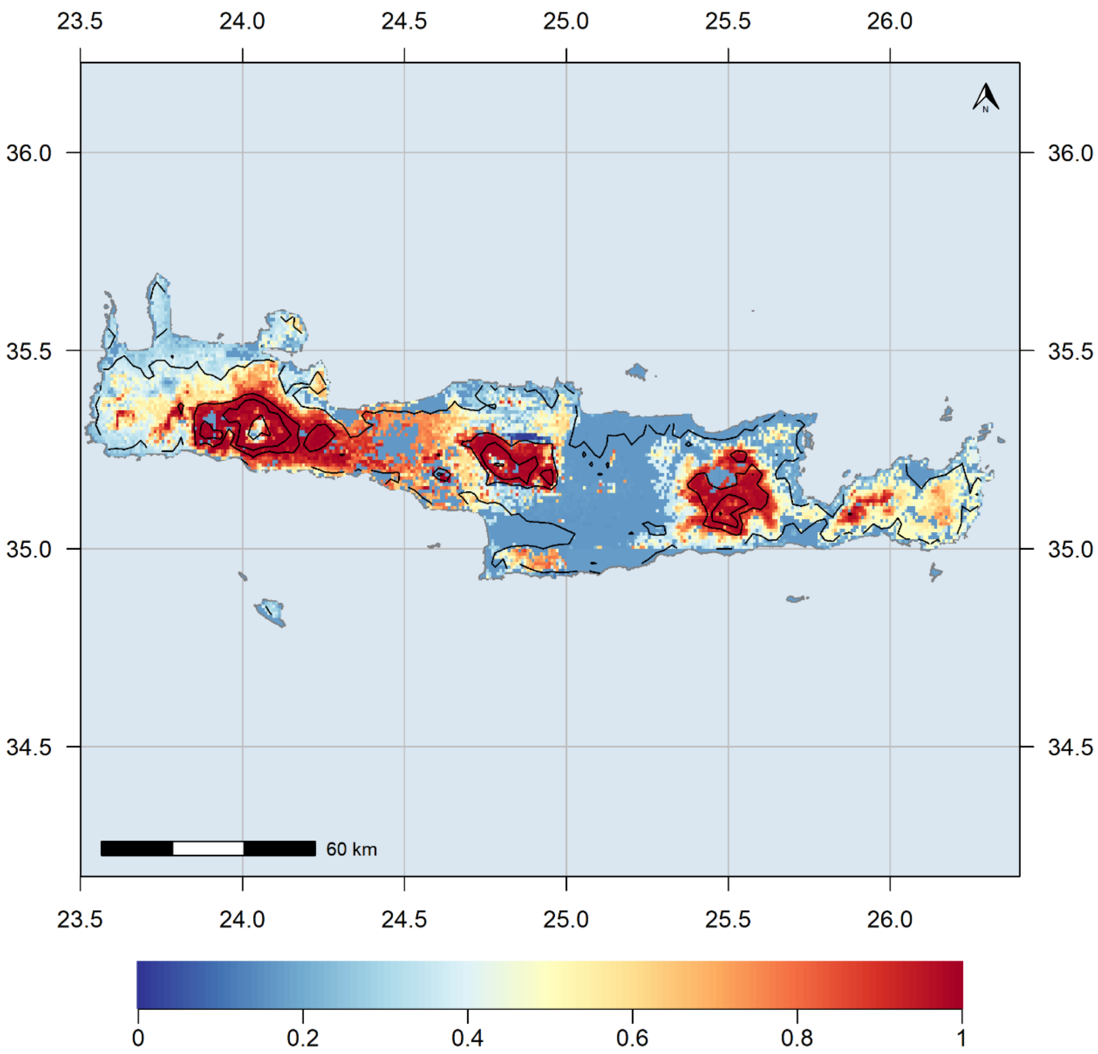


**Figure S16.** Relative difference between current hotspots and the BCC 8.5 GCM/RCP combination.

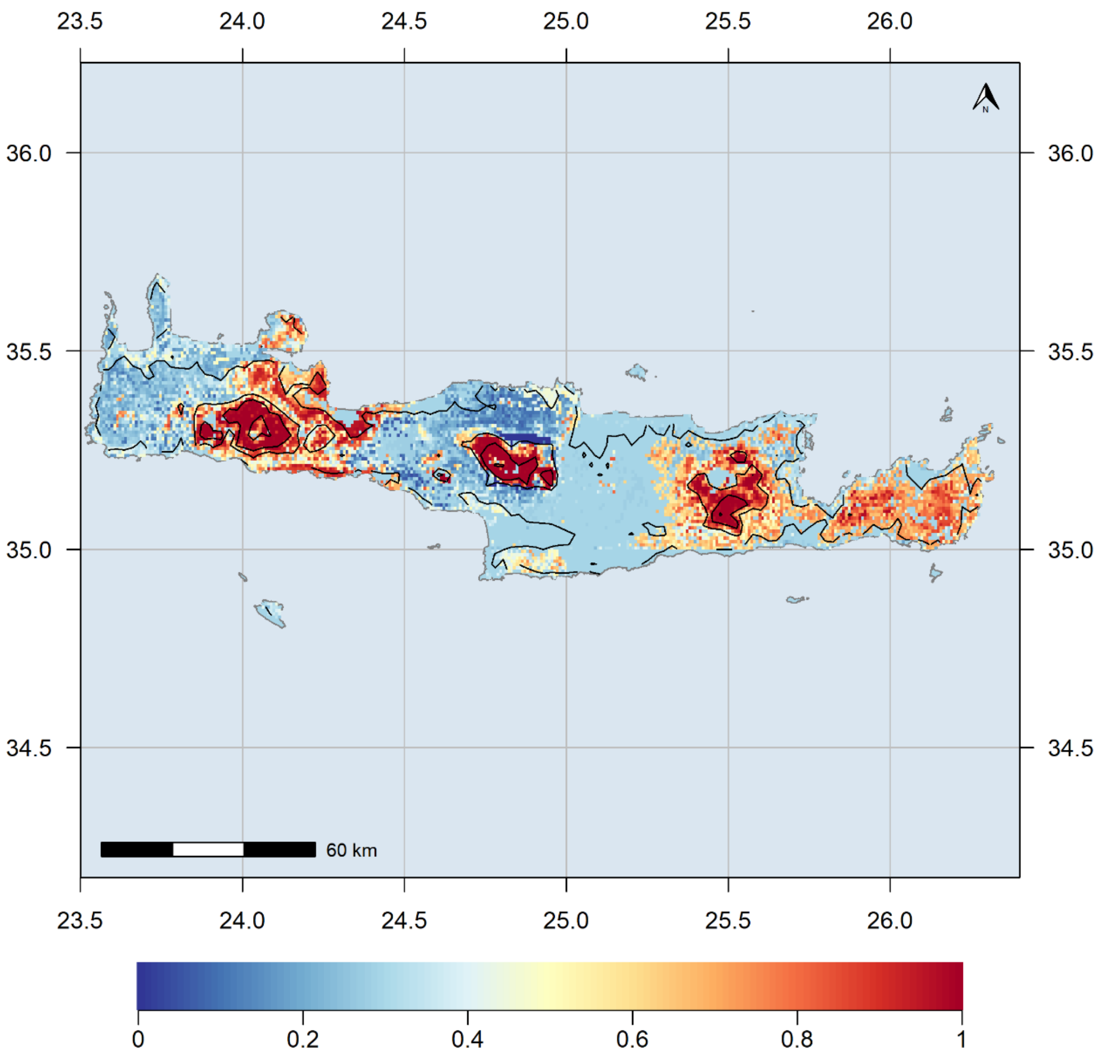


**Figure S17.** Relative difference between current hotspots and the CCSM4 2.6 GCM/RCP combination.

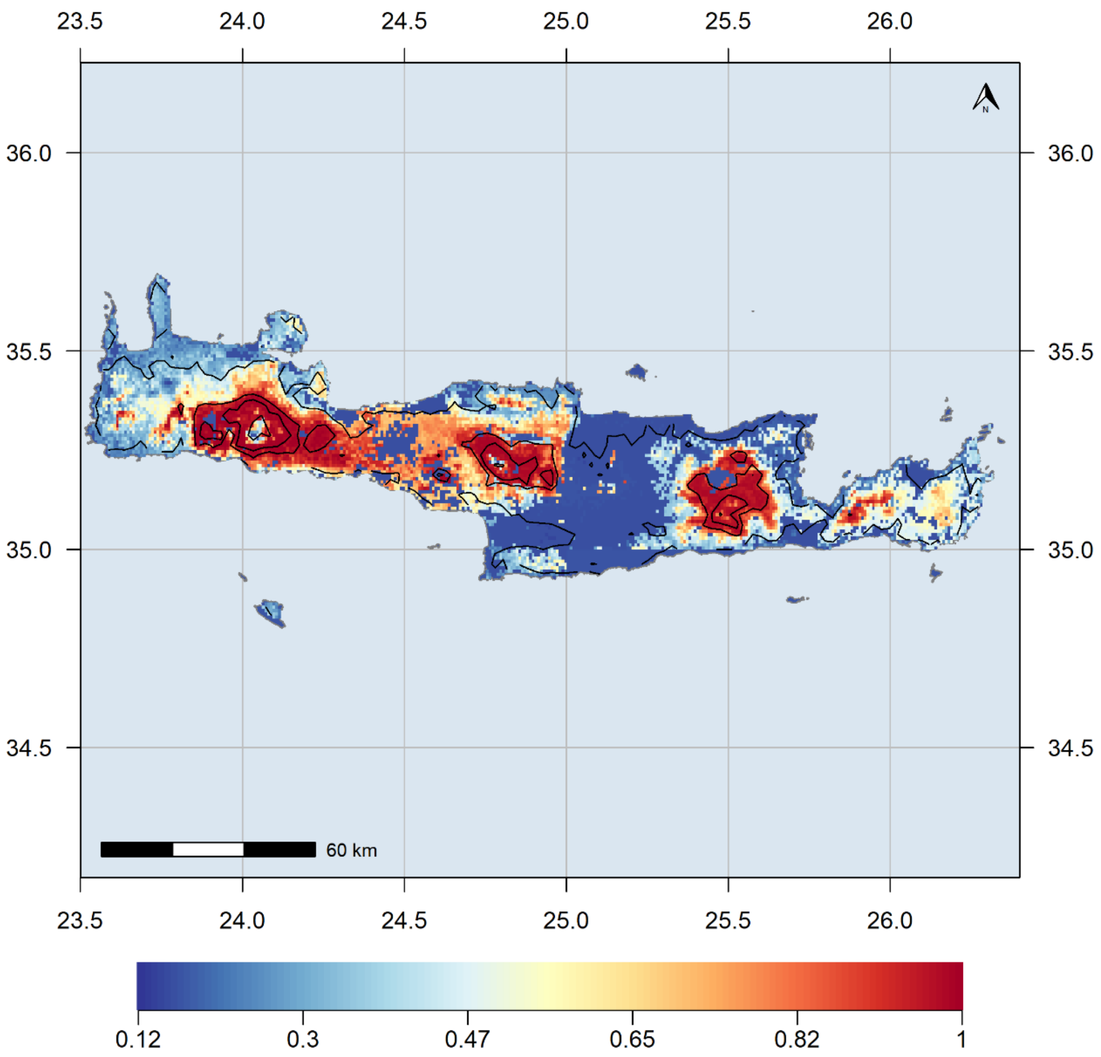


**Figure S18.** Relative difference between current hotspots and the CCSM4 8.5 GCM/RCP combination.

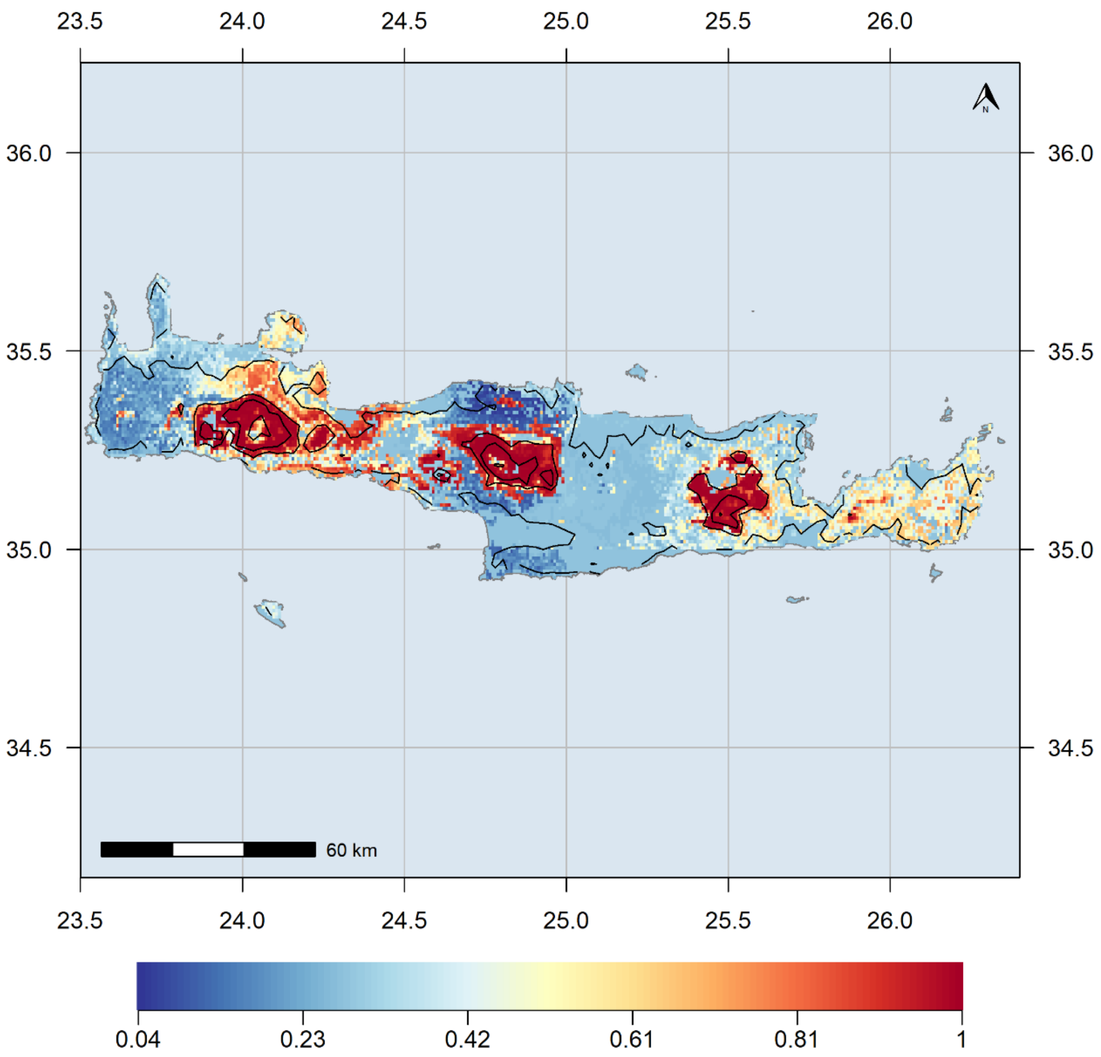


**Figure S19.** Relative difference between current hotspots and the HadGEM2 2.6 GCM/RCP combination.

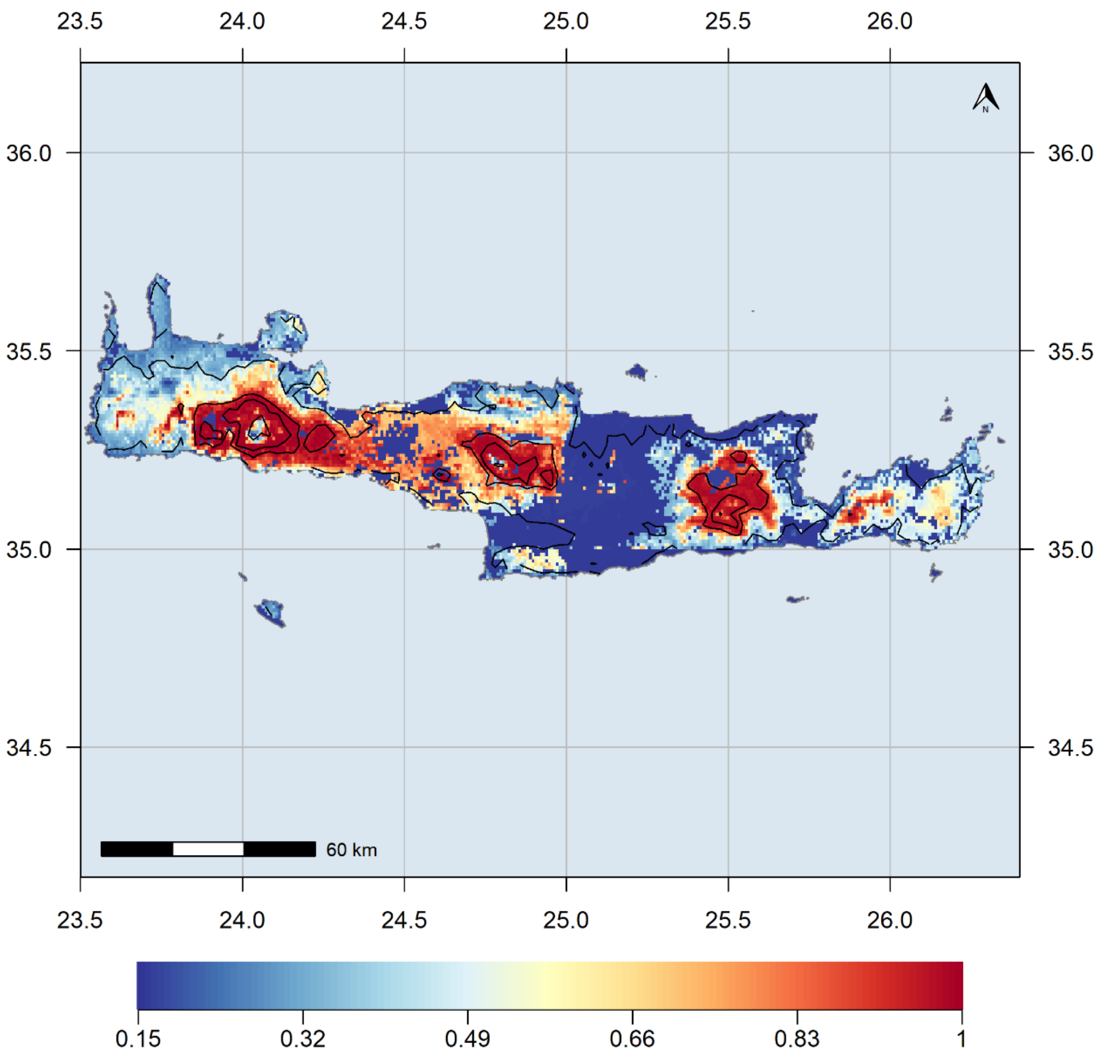


**Figure S20.** Relative difference between current hotspots and the HadGEM2 8.5 GCM/RCP combination.

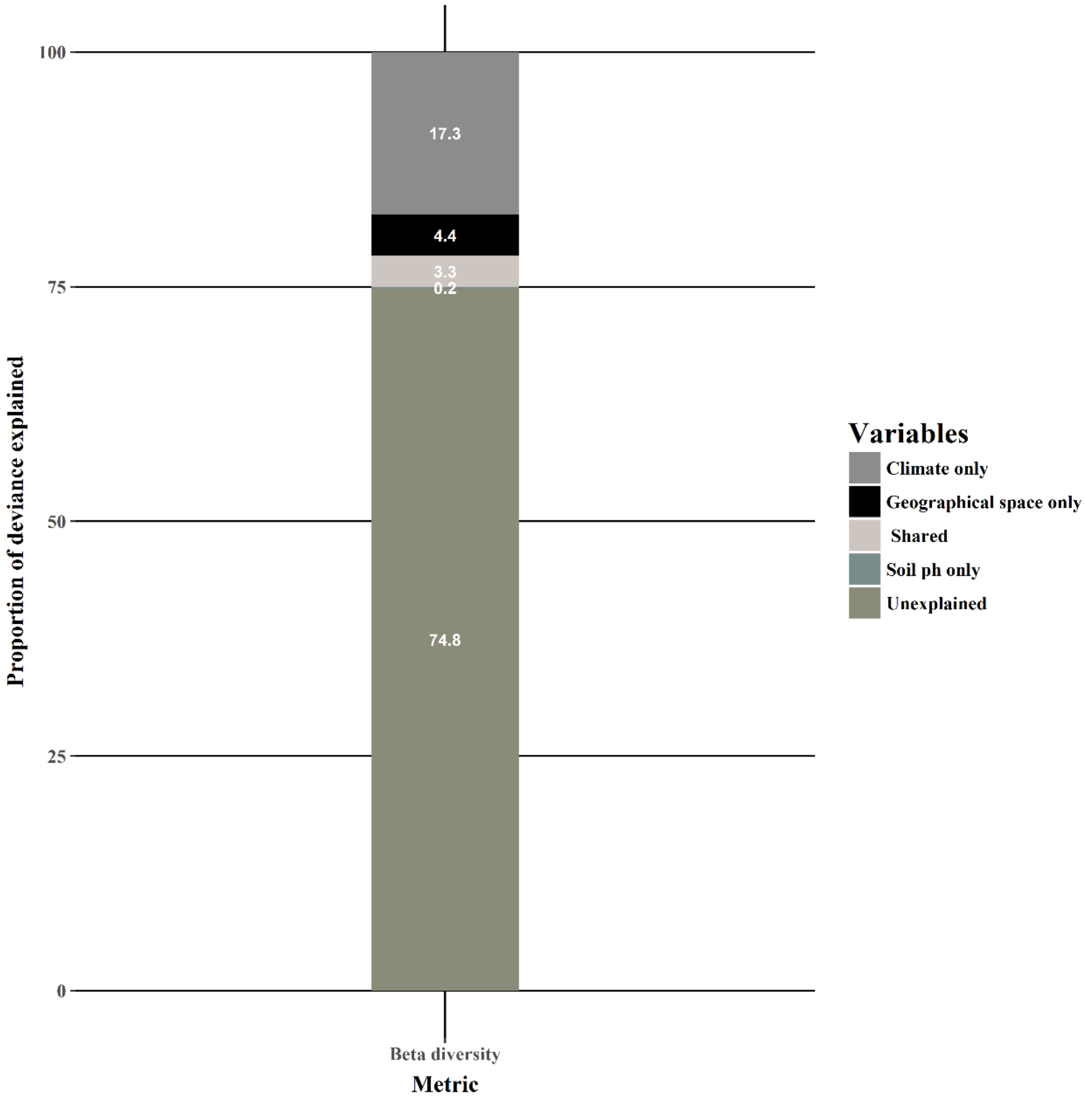


**Fig. S21.** Partition of deviance explained

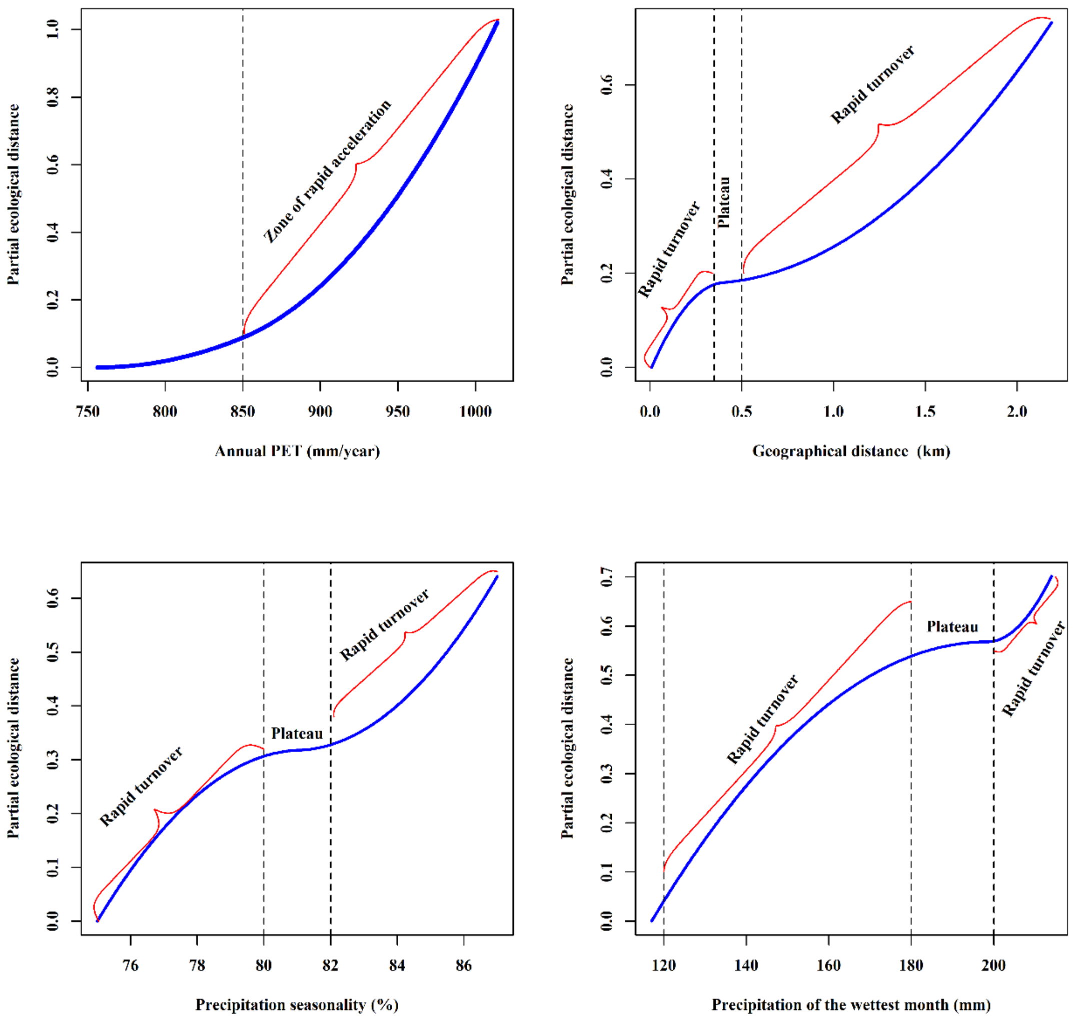


**Fig. S22**. Variable importance for the variables included in the GDM analysis.

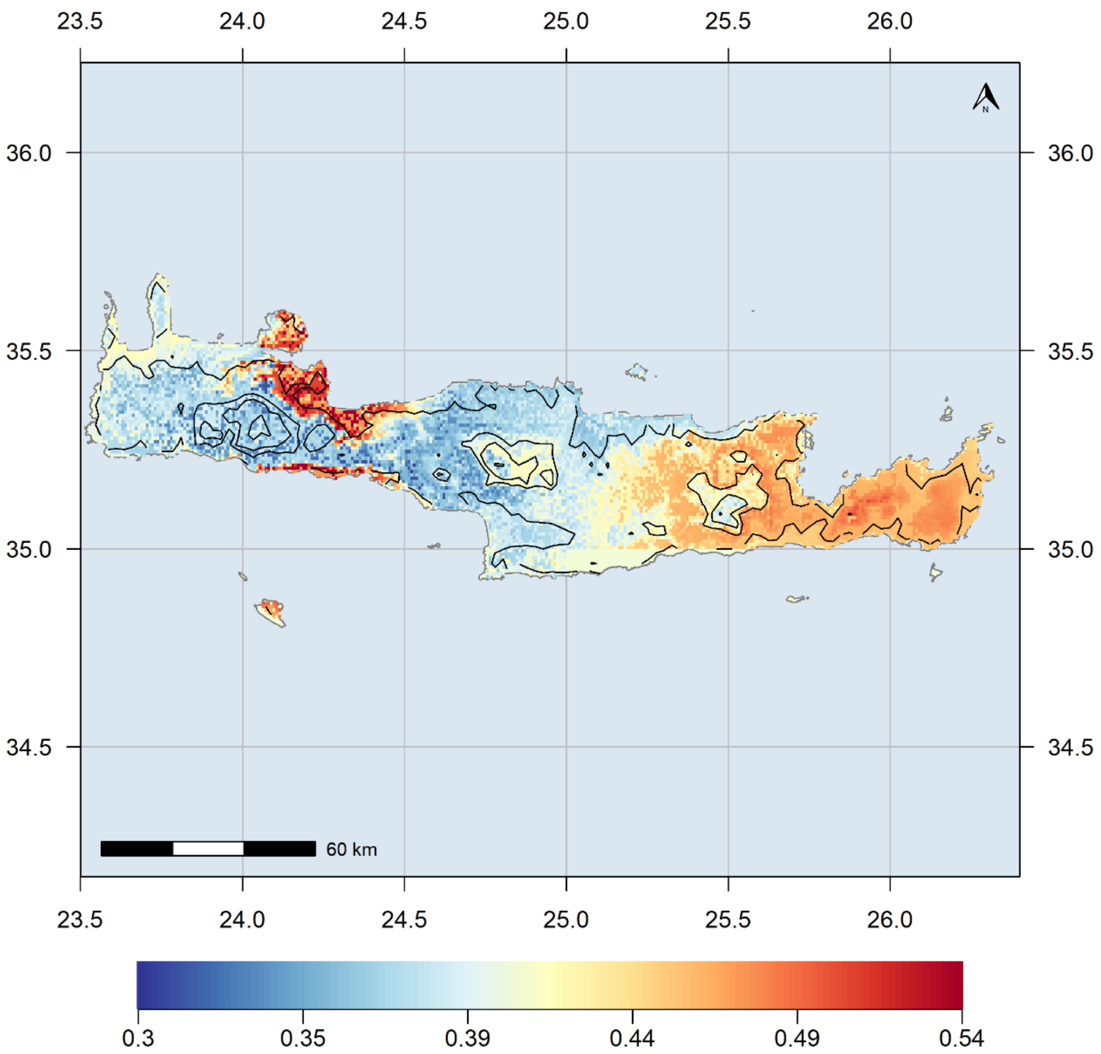


**Figure S23.** Compositional variation for the $SIE_C$ assemblages for the BCC 2.6 GCM/RCP combination.

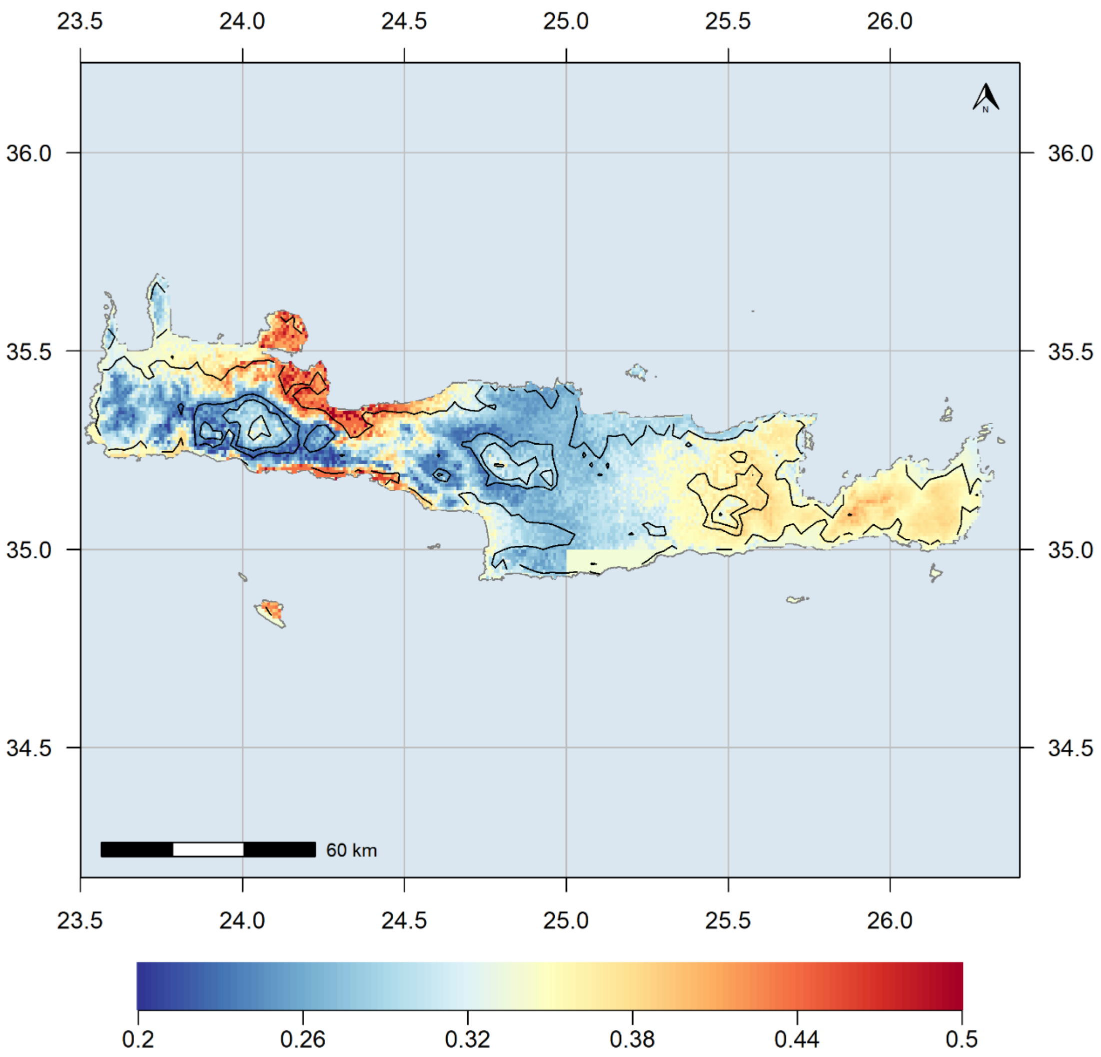

**Figure S24.** Compositional variation for the $SIE_C$ assemblages for the BCC 8.5 GCM/RCP combination.

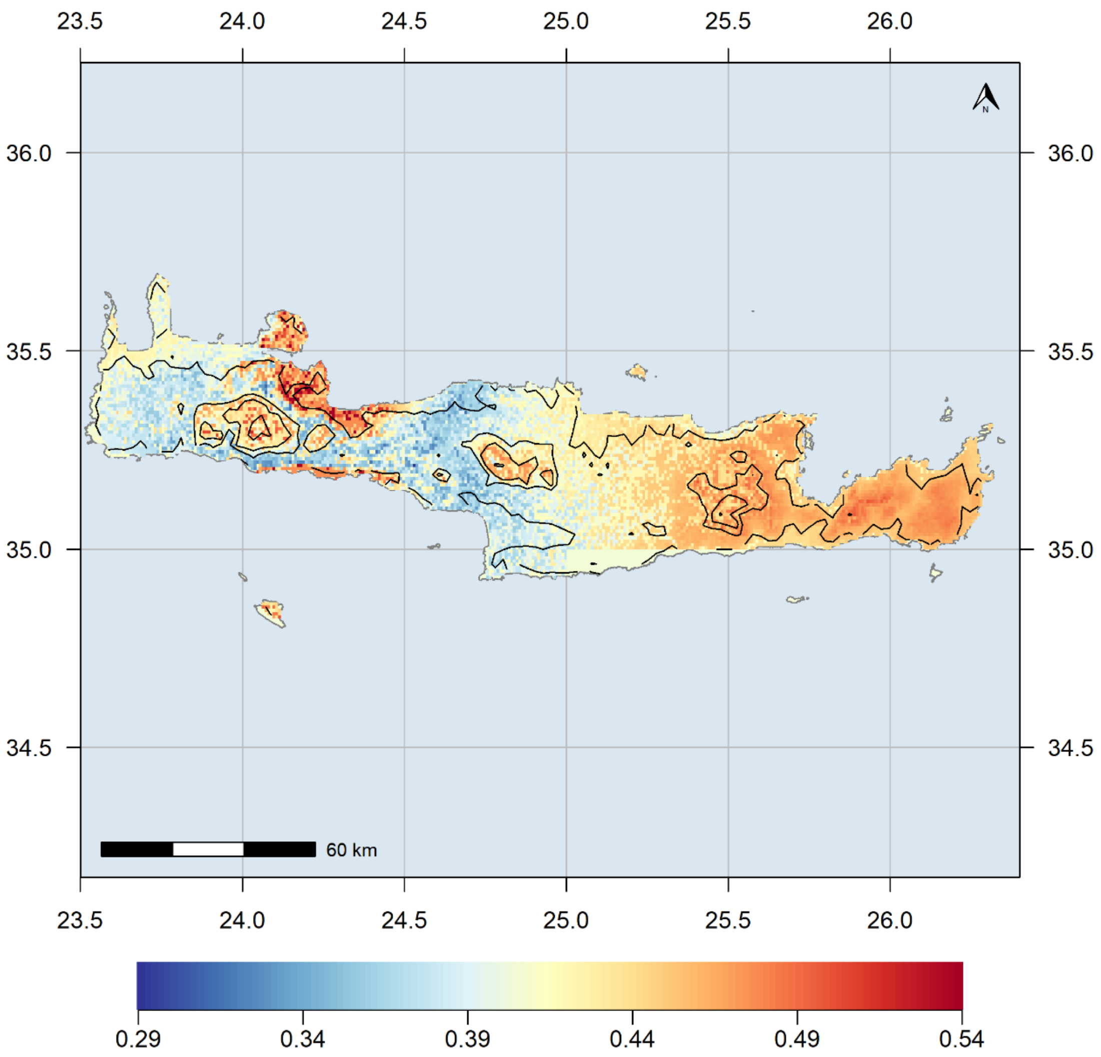


**Figure S25.** Compositional variation for the $SIE_C$ assemblages for the CCSM4 2.6 GCM/RCP combination.

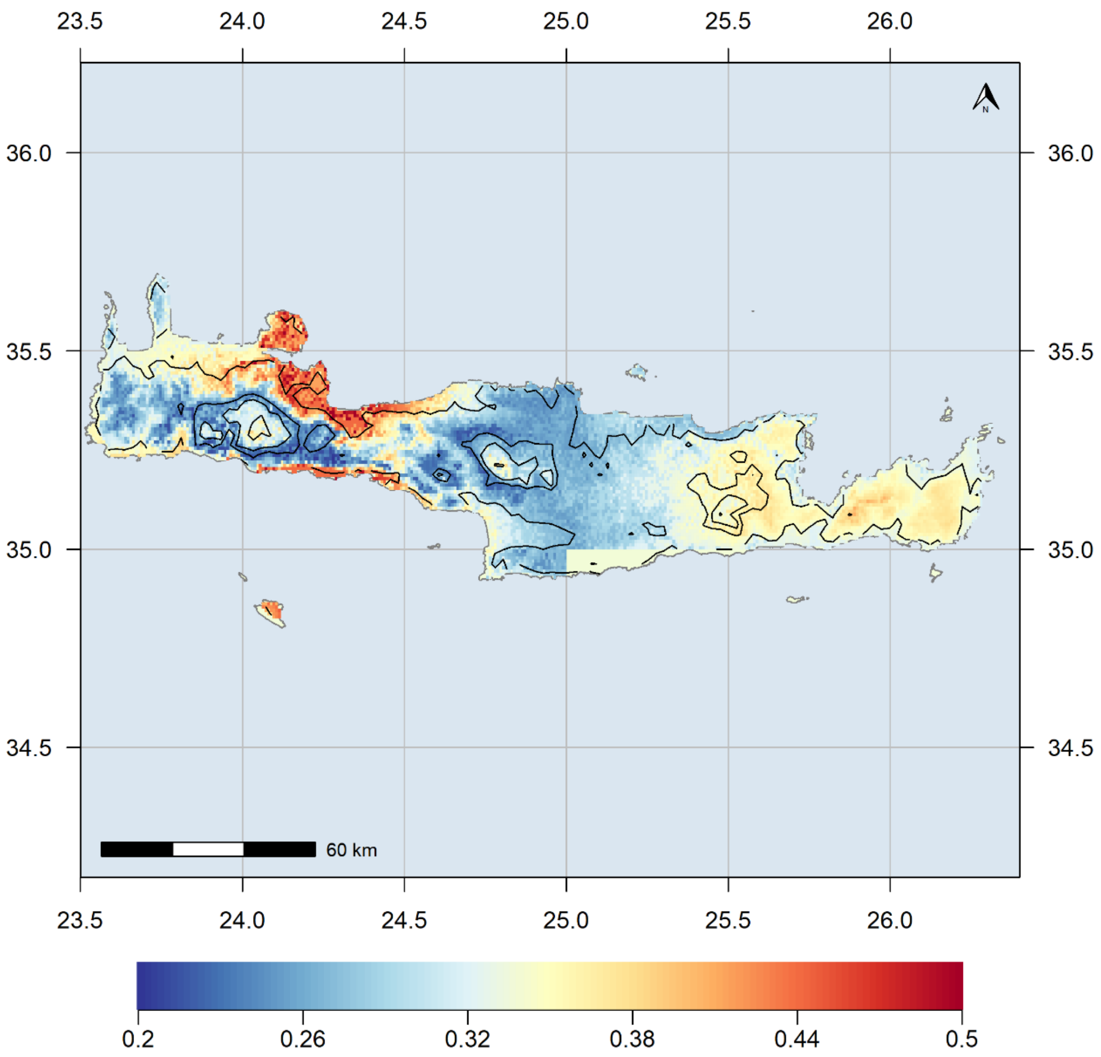


**Figure S26.** Compositional variation for the $SIE_C$ assemblages for the CCSM4 2.6 GCM/RCP combination.

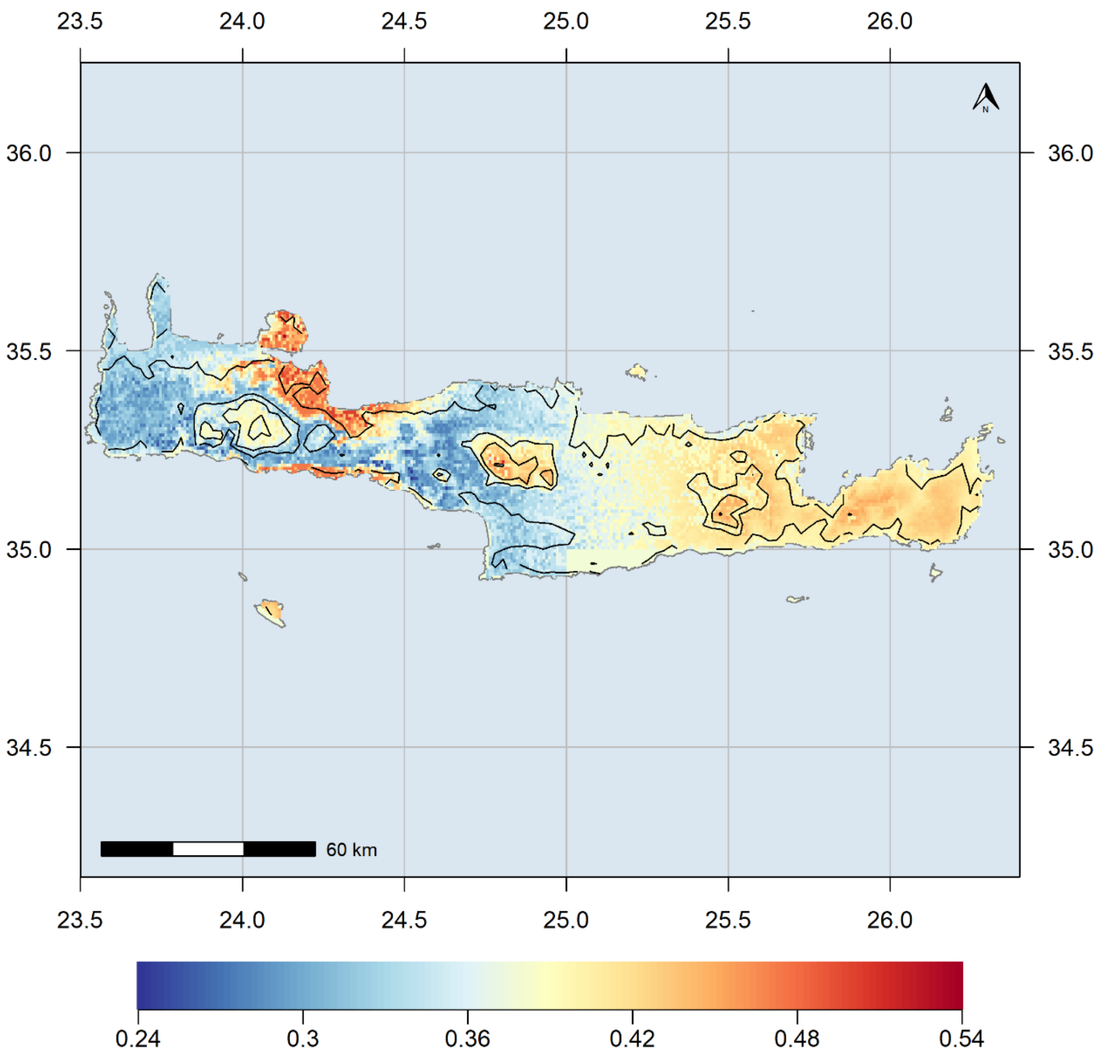


**Figure S27.** Compositional variation for the $SIE_C$ assemblages for the HadGEM2 2.6 GCM/RCP combination.

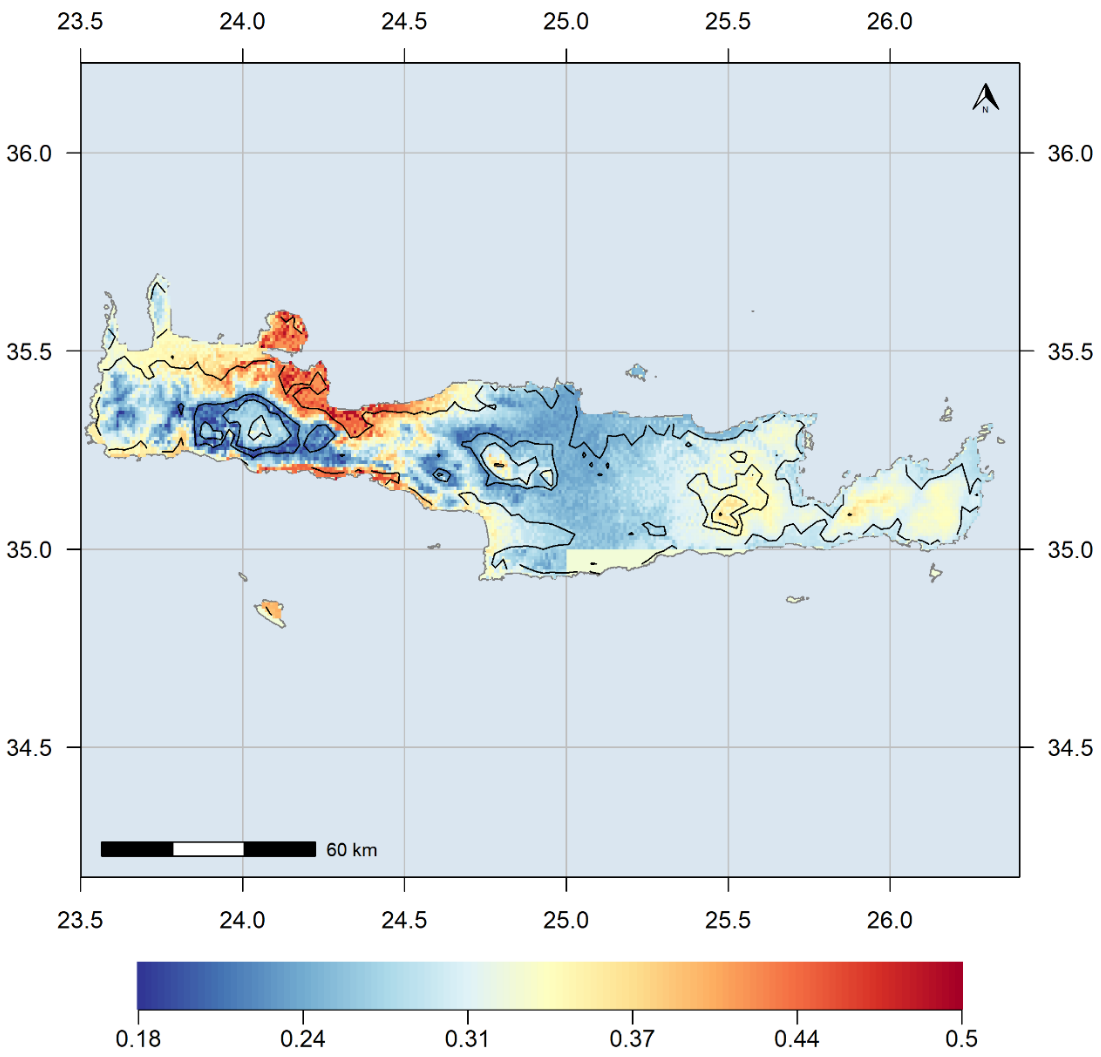


**Figure S28.** Compositional variation for the $SIE_C$ assemblages for the HadGEM2 8.5 GCM/RCP combination.

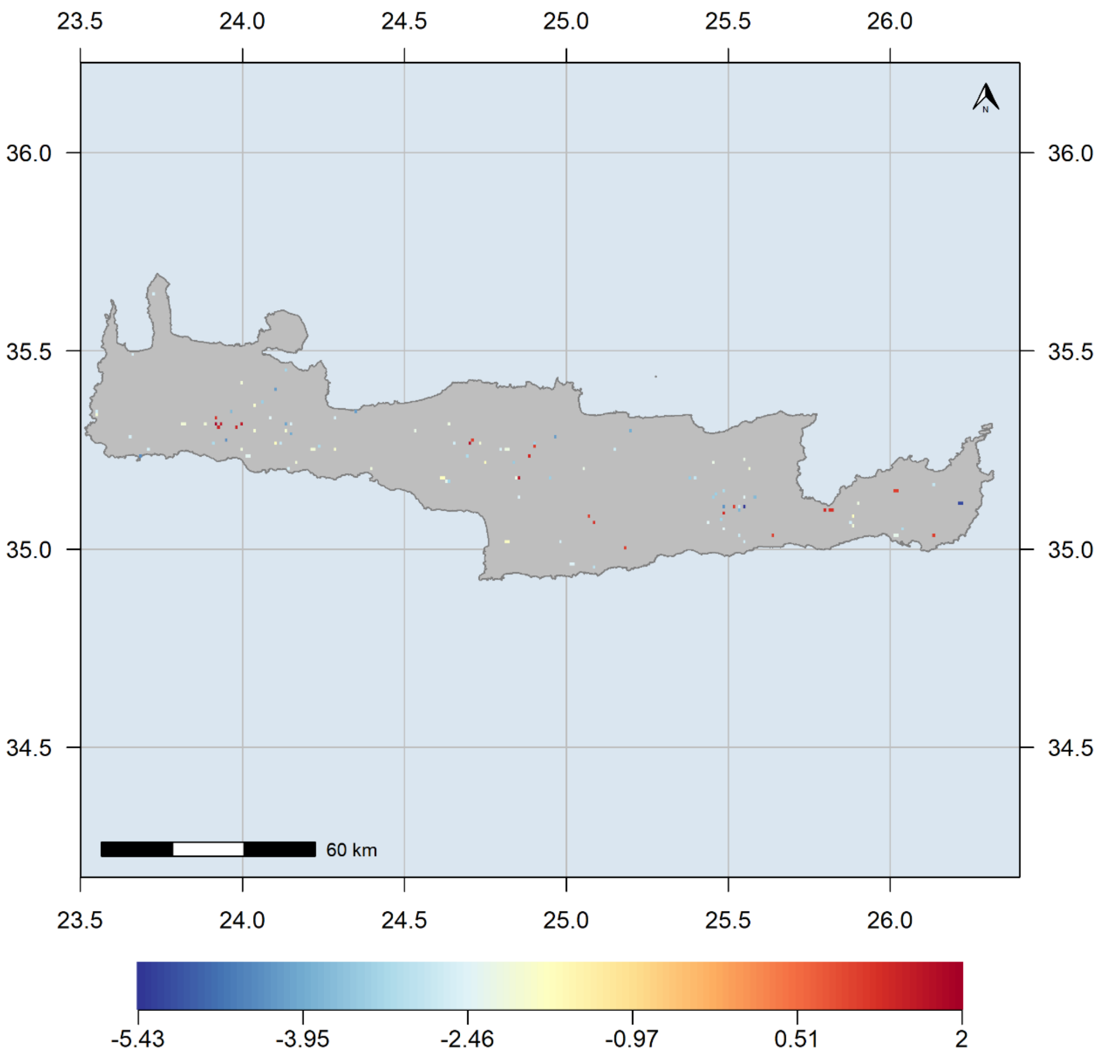


**Figure S29.** Sites with significantly higher or lower than expected PD.

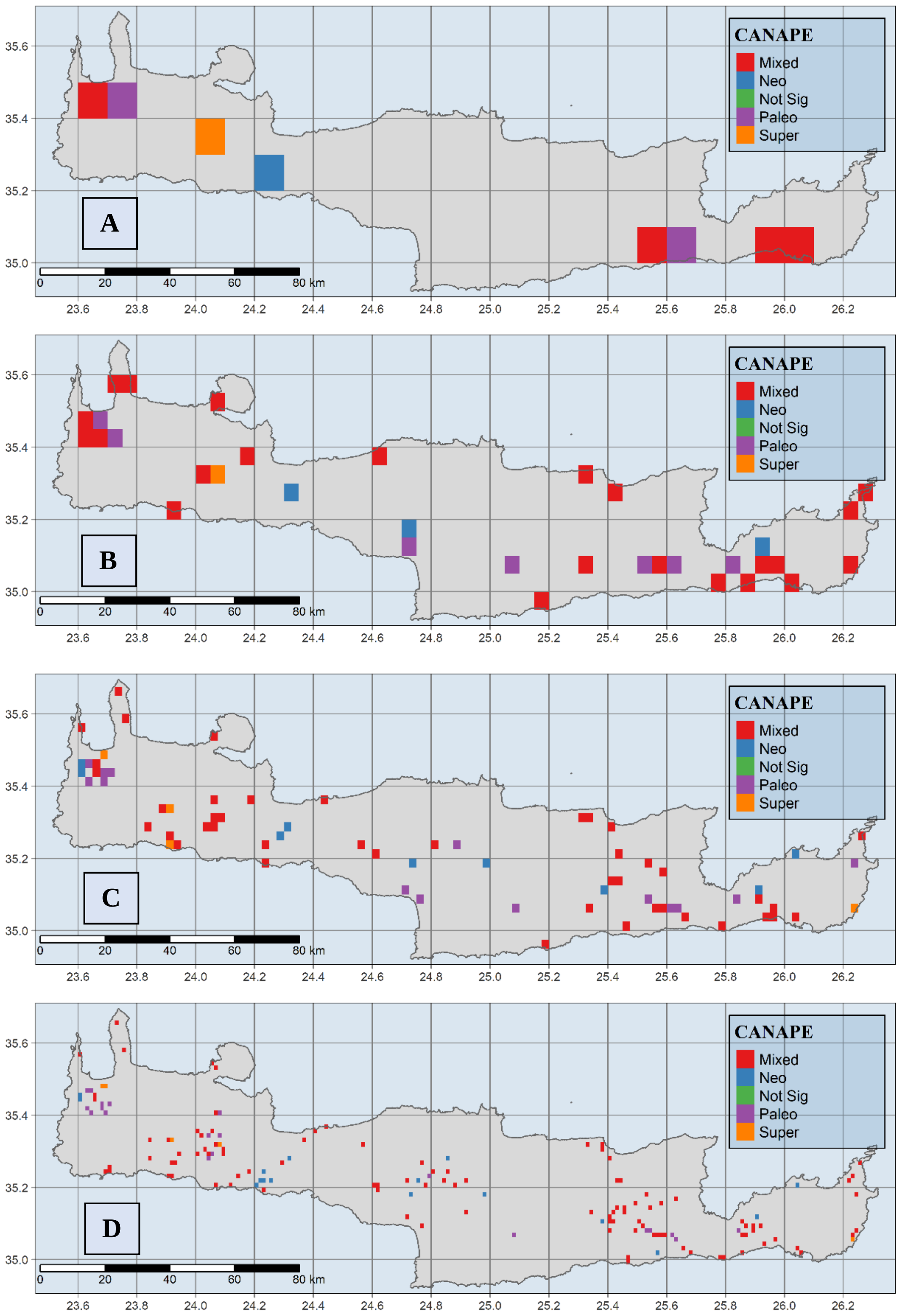


**Figure S30.** CANAPE results estimated for coarser geographical scales. A-D: 0.1, 0.05, 0.025 and 0.0125 degrees, respectively. Overall results are congruent, especially for the mixed-endemism patterns, regardless the grid resolution.

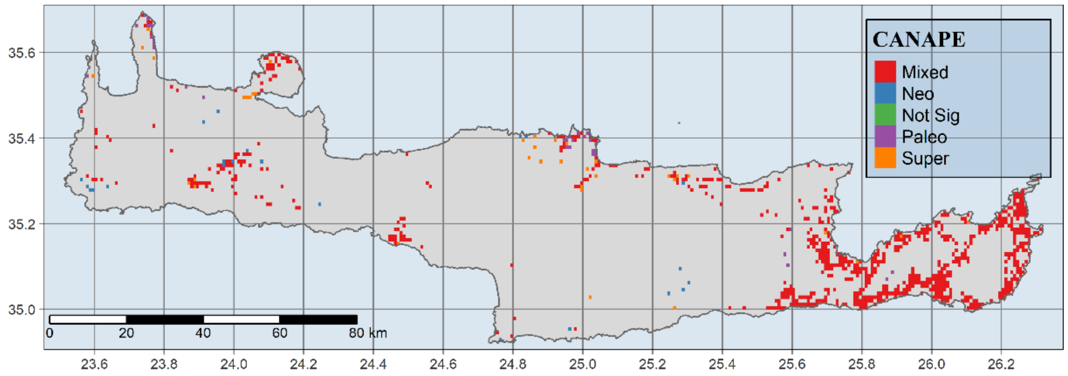


**Figure S31.** Map of significant phylogenetic endemism (PE) identified by the CANAPE analysis for 172 Cretan Single Island Endemics for the BCC 2.6 GCM/RCP combination.

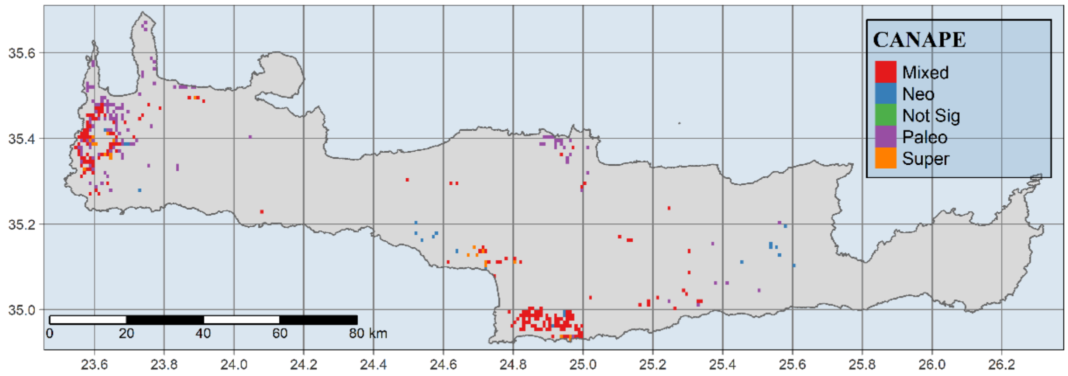


**Figure S32.** Map of significant phylogenetic endemism (PE) identified by the CANAPE analysis for 172 Cretan Single Island Endemics for the BCC 8.5 GCM/RCP combination.

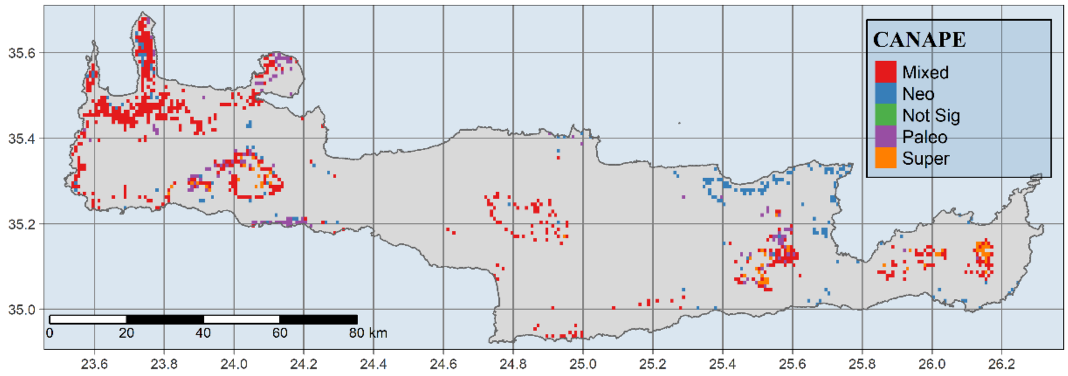


**Figure S33.** Map of significant phylogenetic endemism (PE) identified by the CANAPE analysis for 172 Cretan Single Island Endemics for the CCSM4 2.6 GCM/RCP combination.

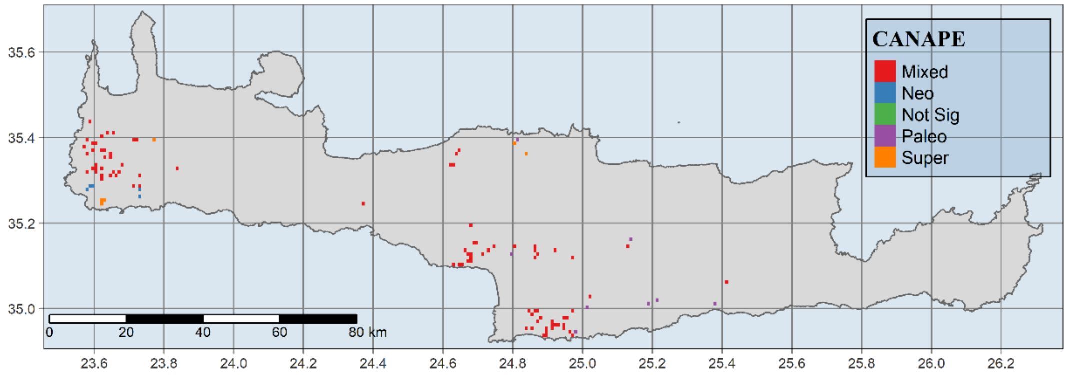


**Figure S34.** Map of significant phylogenetic endemism (PE) identified by the CANAPE analysis for 172 Cretan Single Island Endemics for the CCSM4 8.5 GCM/RCP combination.

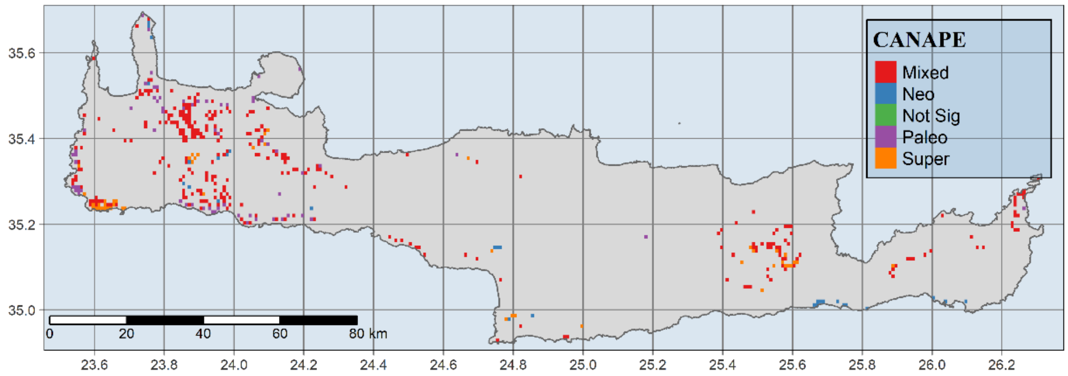


**Figure S35.** Map of significant phylogenetic endemism (PE) identified by the CANAPE analysis for 172 Cretan Single Island Endemics for the HadGEM2 2.6 GCM/RCP combination.

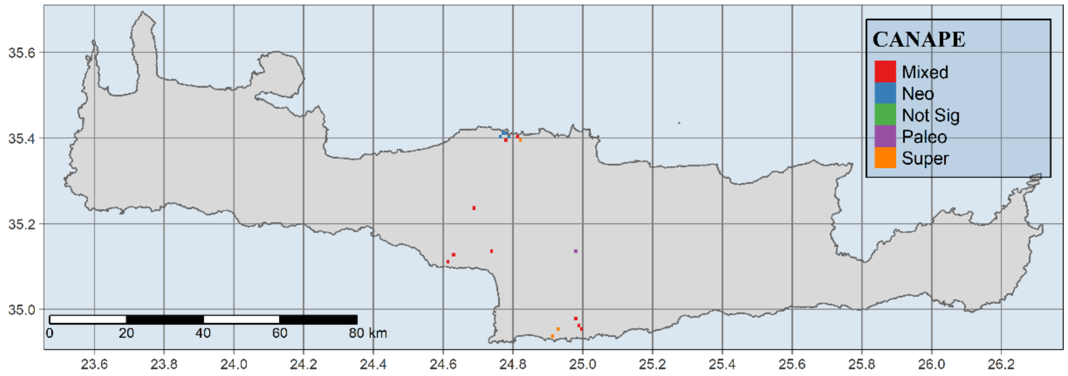


**Figure S36.** Map of significant phylogenetic endemism (PE) identified by the CANAPE analysis for 172 Cretan Single Island Endemics for the HadGEM2 8.5 GCM/RCP combination.

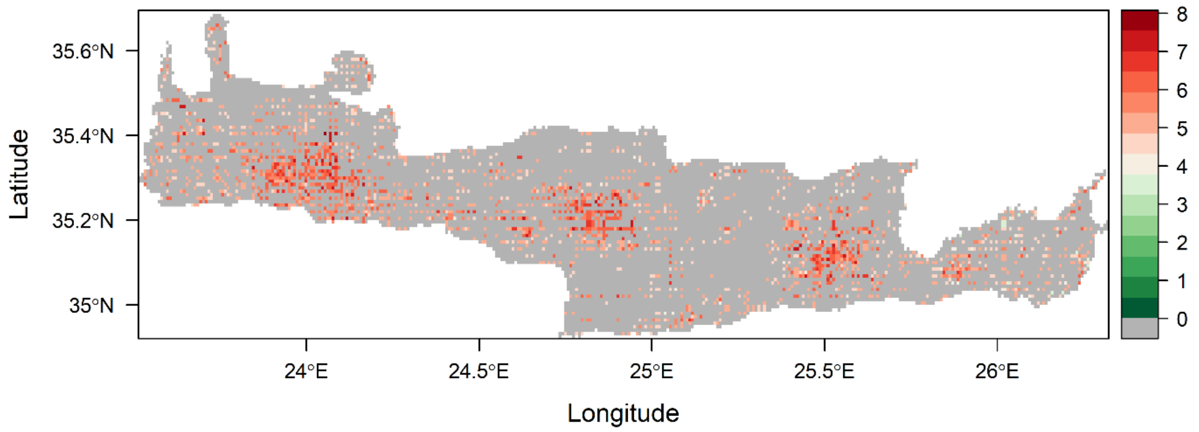


**Figure S37**. Map of Crete showing $SIE_C$ assemblages with their respective EDGE index.

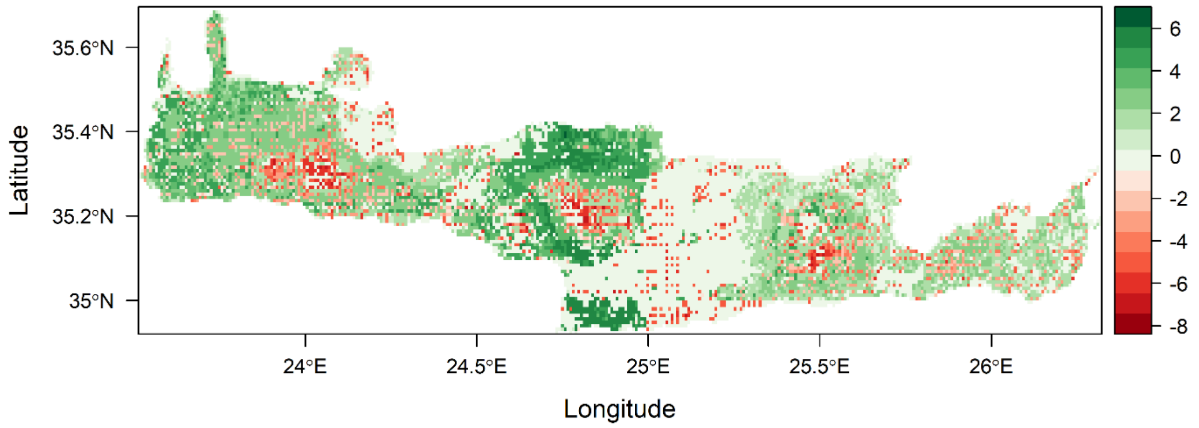


**Figure S38.** Map of Crete showing predicted $SIE_C$ assemblages with their respective ΔEDGE index, averaged between the six GCM/RCP combinations included in our study (BCC 2.6, BCC 8.5, CCSM4 2.6, CCSM4 8.5, HadGEM2 2.6 & HadGEM2 8.5). Red and green areas indicate extinction hotspots and coldspots, respectively.

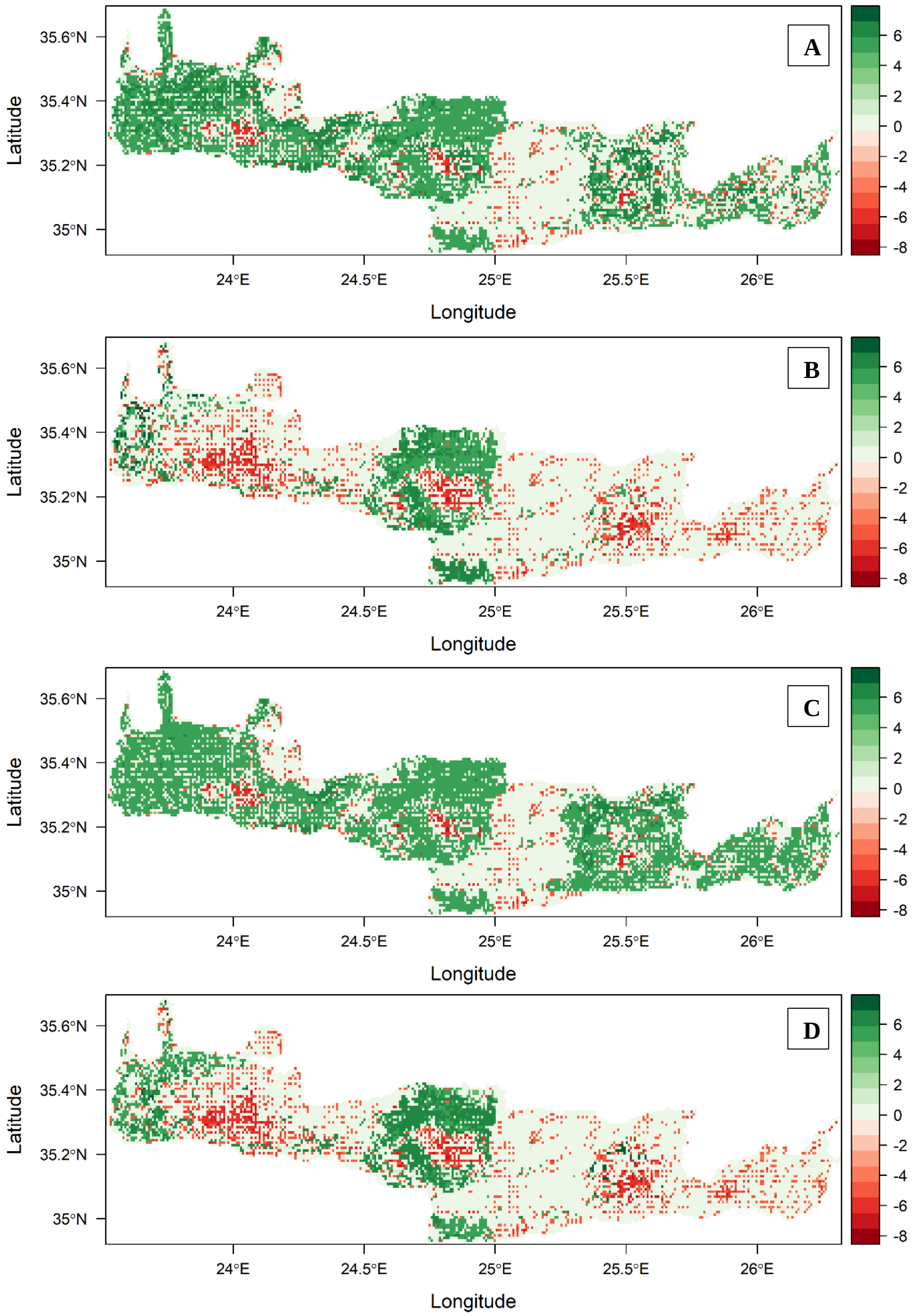

A
B
C
D
35.6°N
35.4°N
35.2°N
35°N
Latitude
24°E
24.5°E
25°E
25.5°E
26°E
Longitude
6
4
2
0
-2
-4
-6
-8

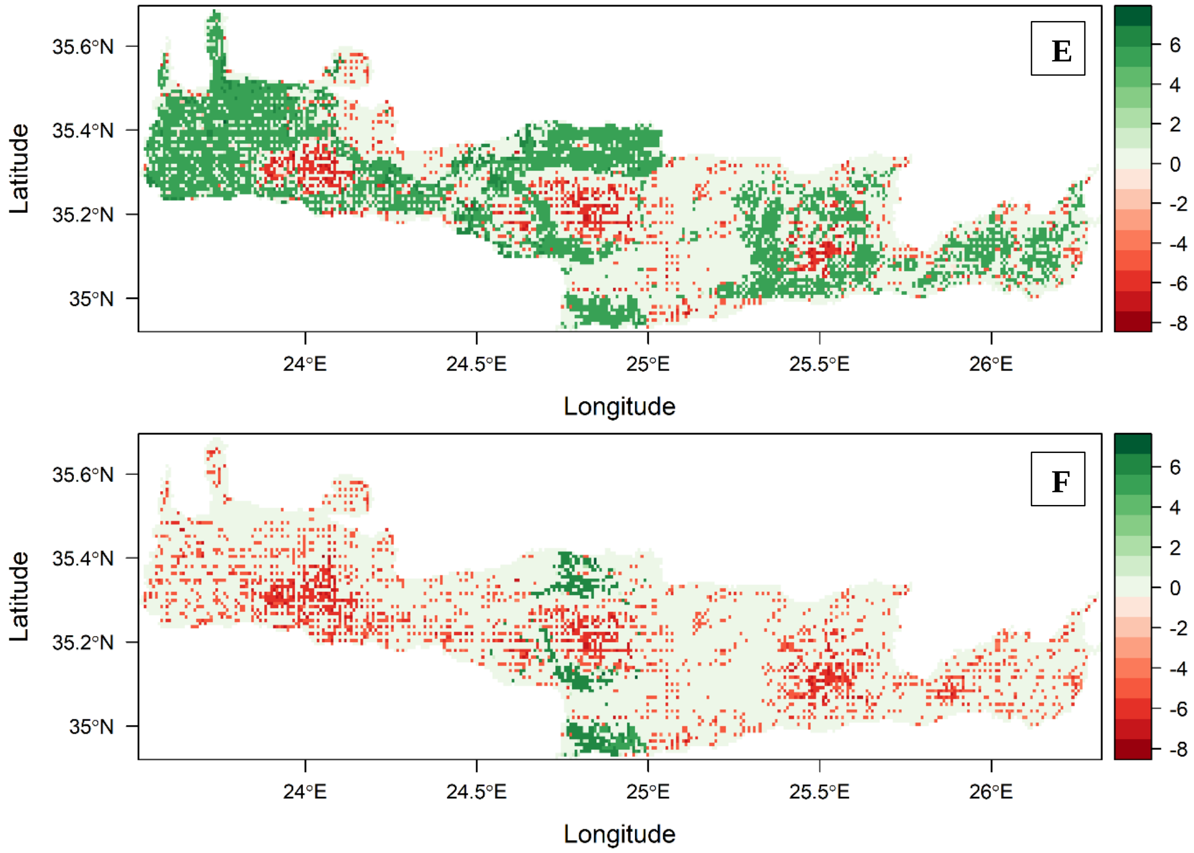


**Figure S39.** Map of Crete showing predicted $SIE_C$ assemblages with their respective ΔEDGE index, for the (A) BCC 2.6, (B) BCC 8.5, (C) CCSM4 2.6, (D) CCSM4 8.5, (E) HadGEM2 2.6 & (F) HadGEM2 8.5 GCM/RCP combination. Red and green areas indicate extinction hotspots and coldspots, respectively.

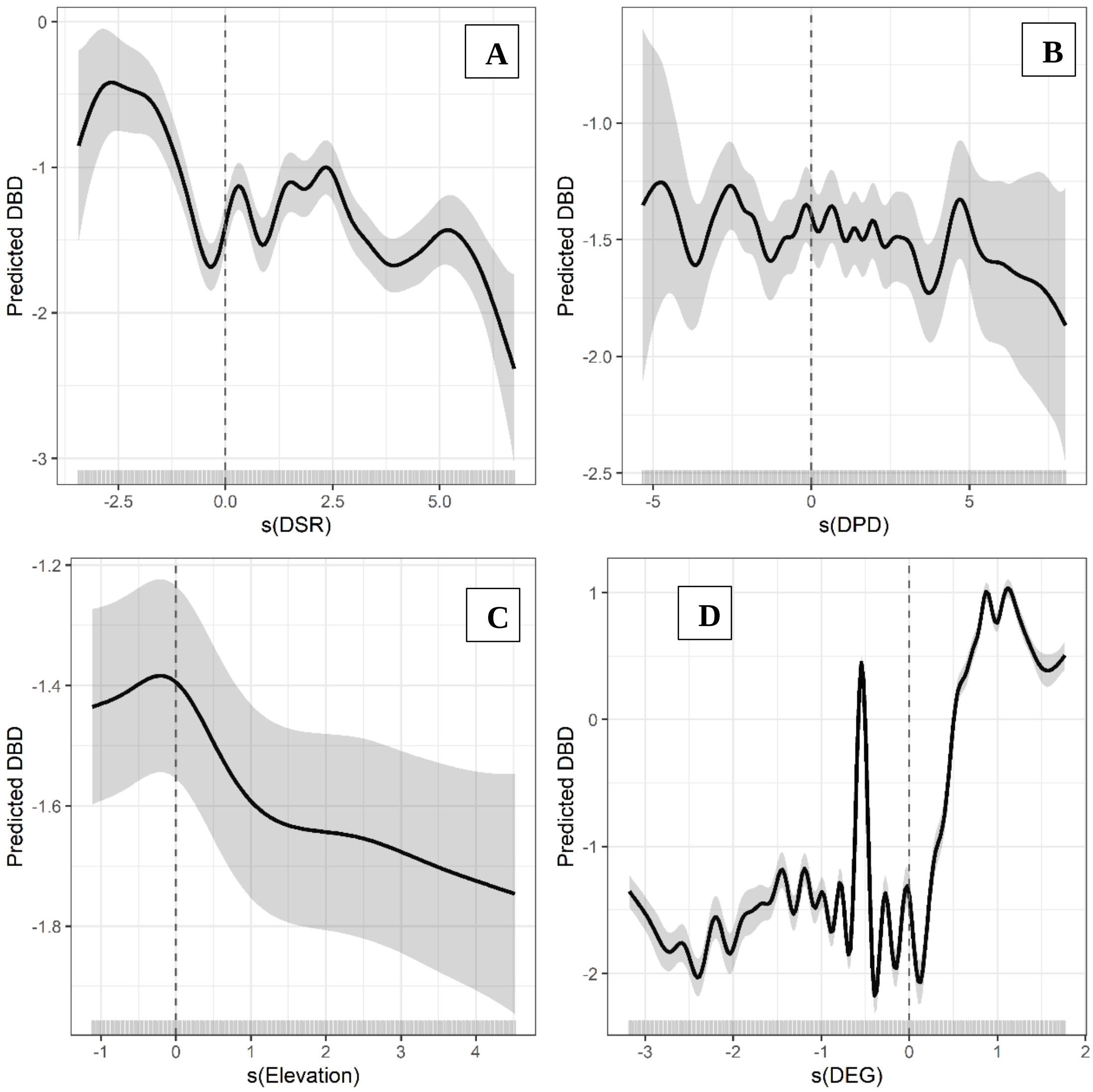


**Figure S40.** Biplots showing the predicted relationships between $\beta_{sim}$ change and the four environmental predictors included in the best generalized additive model (GAM) for the BCC 2.6 GCM/RCP combination. (A). Change in species richness (ΔSR). (B). Change in phylogenetic diversity (ΔPD). (C). Change in elevation. (D). Change in the average level of ecological generalism (ΔEG). Fitted lines show the univariate GAMs with 95% confidence interval (dark grey). Rugs on the x-axes show the predictor values and how they are distributed. Labels on the y-axes indicate the smooth functions for the term of interest (ΔSR, ΔPD, ΔEG and elevation) and the estimated degrees of freedom (following the term). Values above and below the horizontal dashed line indicate heterogenization and homogenization, respectively. Values left and right of the vertical dashed line indicate in (A) species loss and gain, (B) PD decrease and increase and (D) assemblages composed by specialists and generalists, respectively.

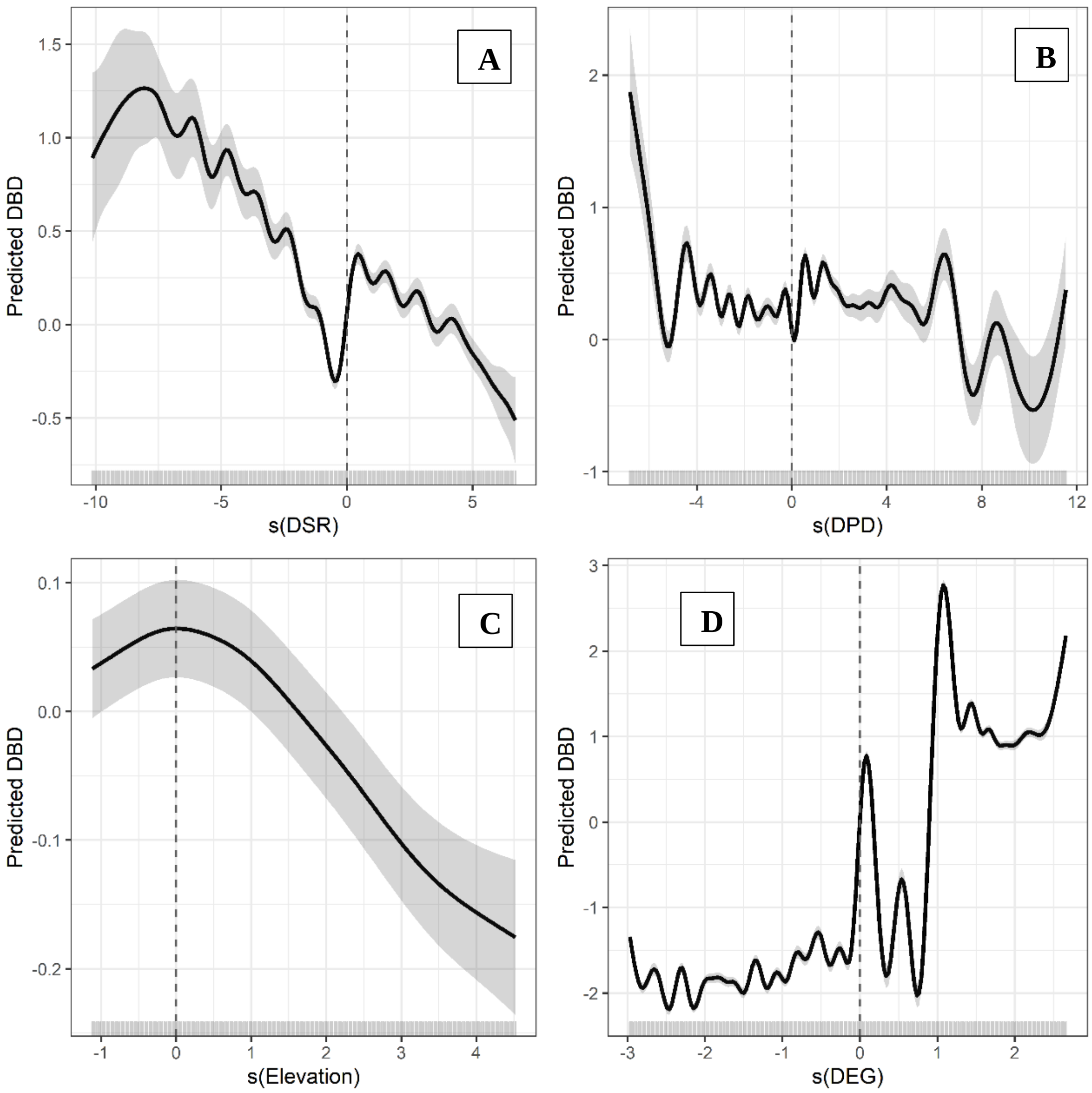


**Figure S41.** Biplots showing the predicted relationships between $\beta_{sim}$ change and the four environmental predictors included in the best generalized additive model (GAM) for the BCC 8.5 GCM/RCP combination. (A). Change in species richness (ΔSR). (B). Change in phylogenetic diversity (ΔPD). (C). Change in elevation. (D). Change in the average level of ecological generalism (ΔEG). Fitted lines show the univariate GAMs with 95% confidence interval (dark grey). Rugs on the x-axes show the predictor values and how they are distributed. Labels on the y-axes indicate the smooth functions for the term of interest (ΔSR, ΔPD, ΔEG and elevation) and the estimated degrees of freedom (following the term). Values above and below the horizontal dashed line indicate heterogenization and homogenization, respectively. Values left and right of the vertical dashed line indicate in (A) species loss and gain, (B) PD decrease and increase and (D) assemblages composed by specialists and generalists, respectively.

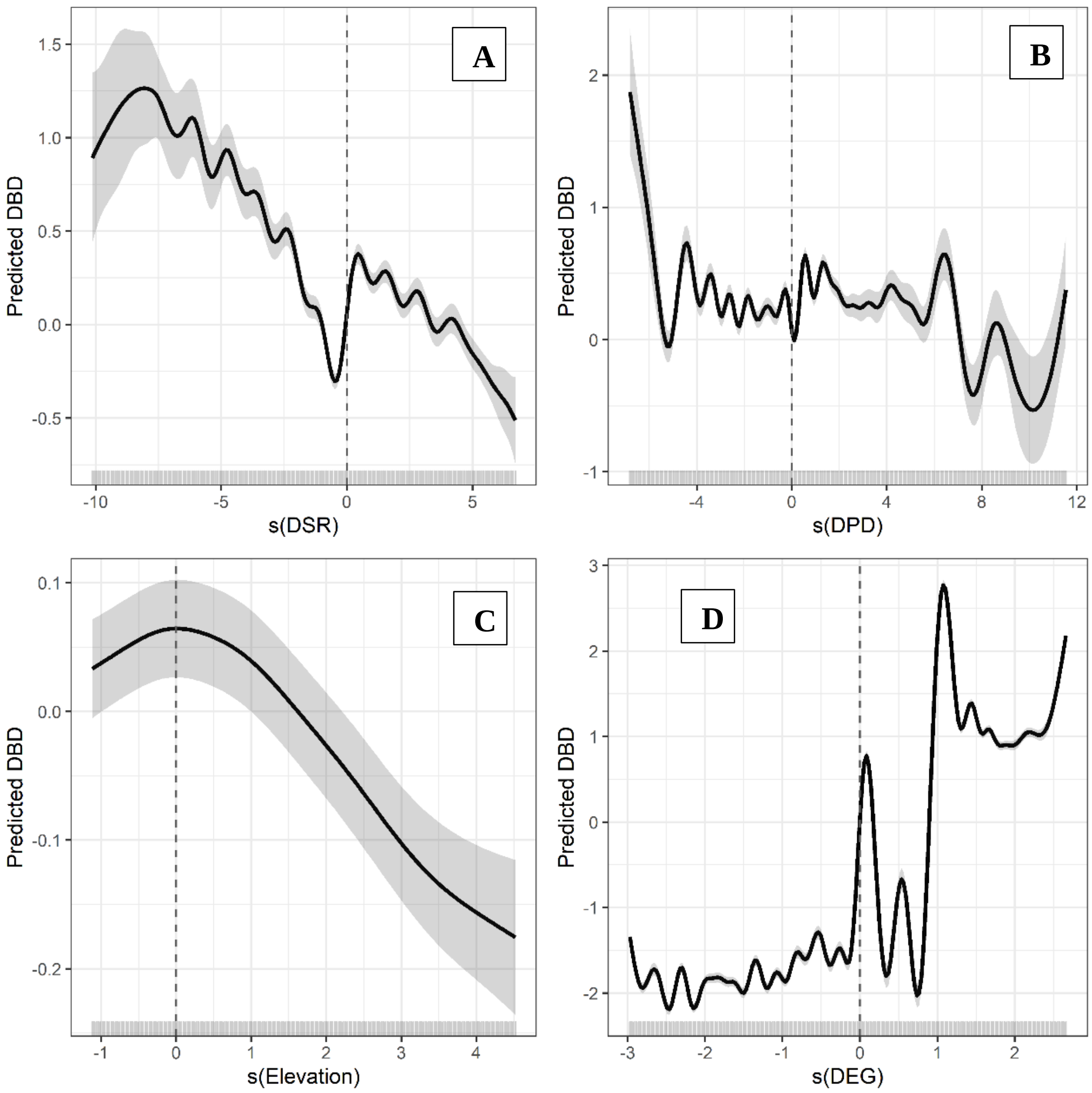


**Figure S42.** Biplots showing the predicted relationships between $\beta_{sim}$ change and the four environmental predictors included in the best generalized additive model (GAM) for the CCSM4 8.5 GCM/RCP combination. (A). Change in species richness (ΔSR). (B). Change in phylogenetic diversity (ΔPD). (C). Change in elevation. (D). Change in the average level of ecological generalism (ΔEG). Fitted lines show the univariate GAMs with 95% confidence interval (dark grey). Rugs on the x-axes show the predictor values and how they are distributed. Labels on the y-axes indicate the smooth functions for the term of interest (ΔSR, ΔPD, ΔEG and elevation) and the estimated degrees of freedom (following the term). Values above and below the horizontal dashed line indicate heterogenization and homogenization, respectively. Values left and right of the vertical dashed line indicate in (A) species loss and gain, (B) PD decrease and increase and (D) assemblages composed by specialists and generalists, respectively.

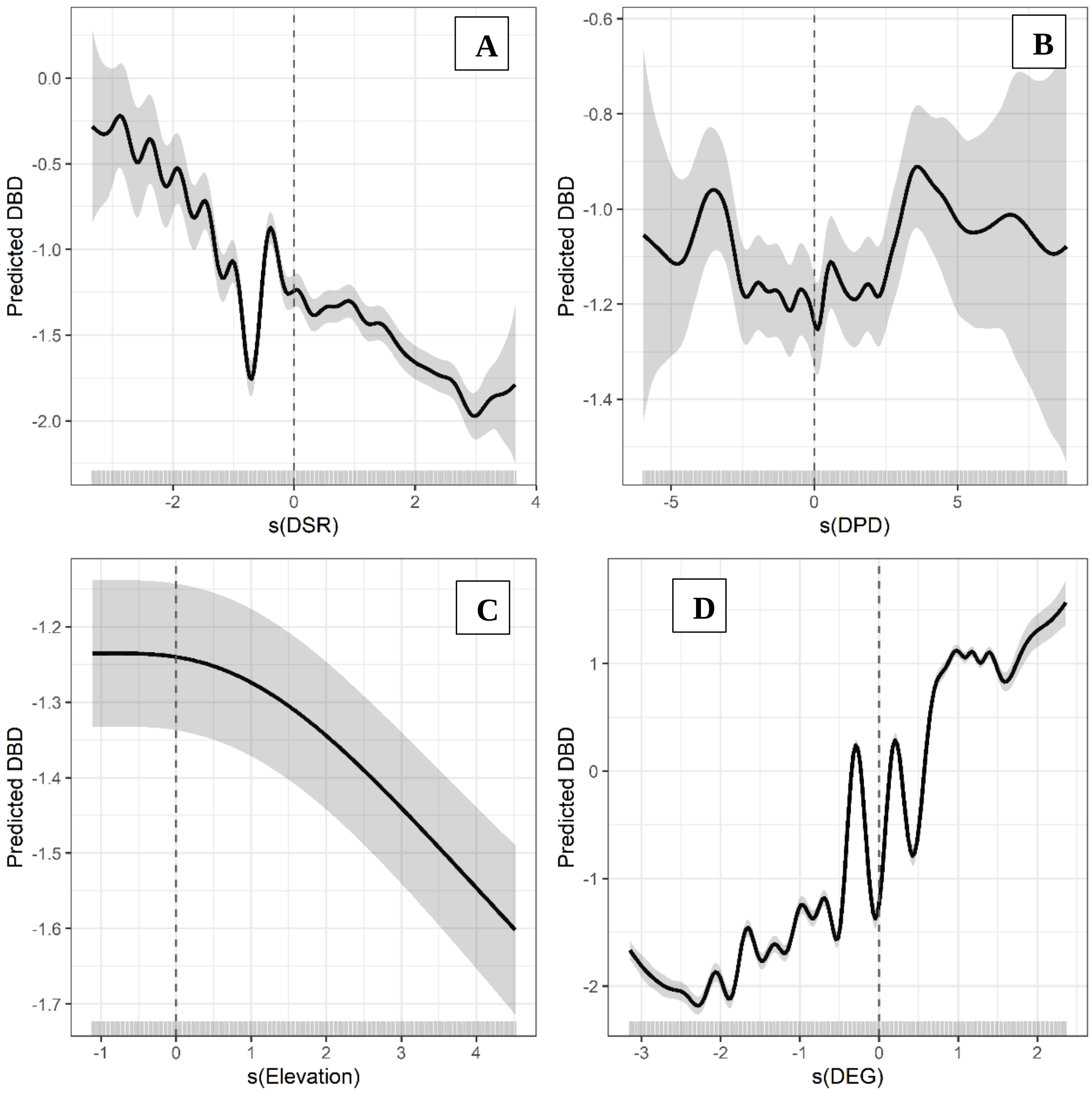


**Figure S 43.** Biplots showing the predicted relationships between $\beta_{sim}$ change and the four environmental predictors included in the best generalized additive model (GAM) for the HadGEM2 2.6 GCM/RCP combination. (A). Change in species richness (ΔSR). (B). Change in phylogenetic diversity (ΔPD). (C). Change in elevation. (D). Change in the average level of ecological generalism (ΔEG). Fitted lines show the univariate GAMs with 95% confidence interval (dark grey). Rugs on the x-axes show the predictor values and how they are distributed. Labels on the y-axes indicate the smooth functions for the term of interest (ΔSR, ΔPD, ΔEG and elevation) and the estimated degrees of freedom (following the term). Values above and below the horizontal dashed line indicate heterogenization and homogenization, respectively. Values left and right of the vertical dashed line indicate in (A) species loss and gain, (B) PD decrease and increase and (D) assemblages composed by specialists and generalists, respectively.

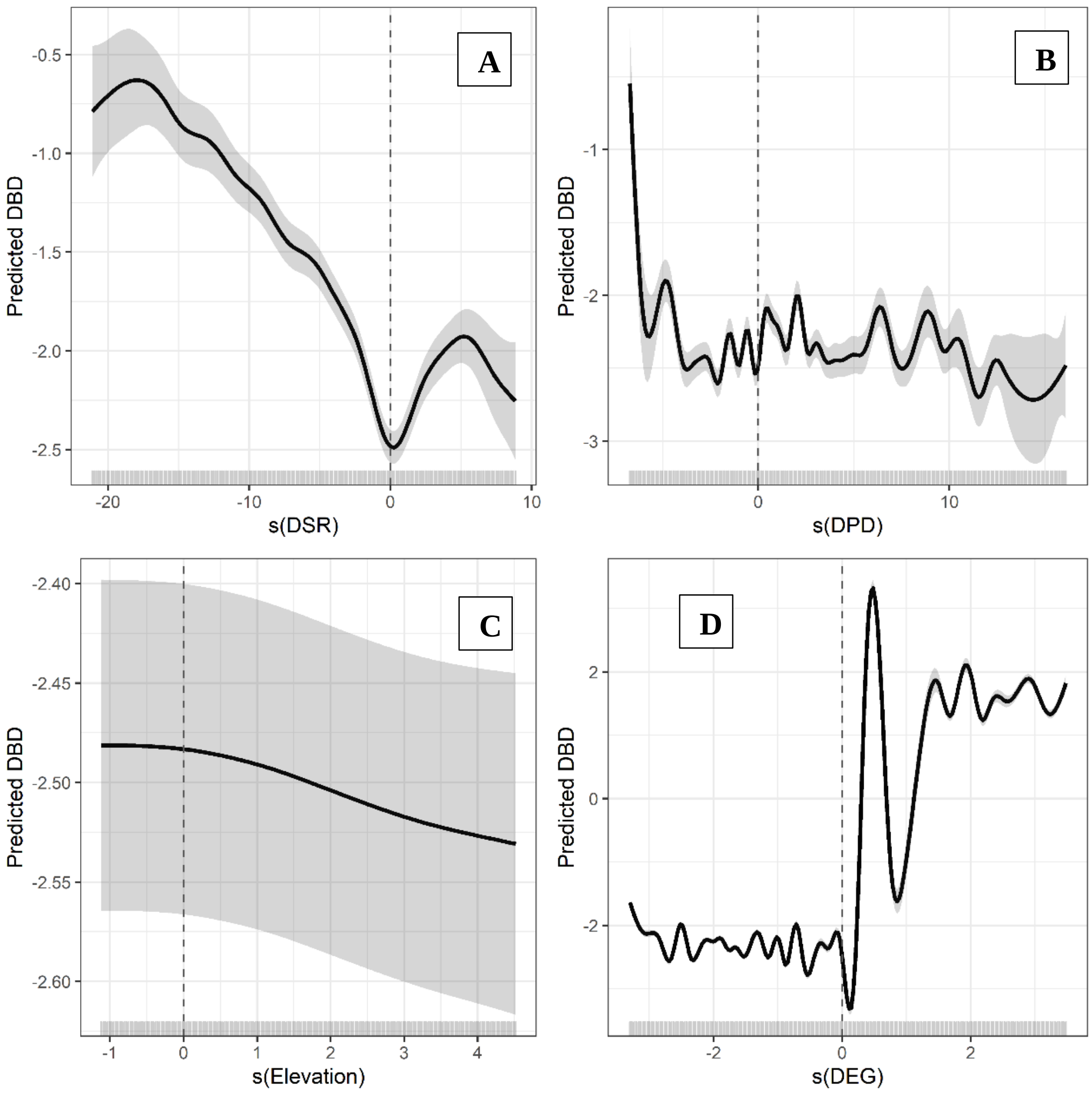


**Figure S44.** Biplots showing the predicted relationships between $\beta_{sim}$ change and the four environmental predictors included in the best generalized additive model (GAM) for the HadGEM2 8.5 GCM/RCP combination. (A). Change in species richness (ΔSR). (B). Change in phylogenetic diversity (ΔPD). (C). Change in elevation. (D). Change in the average level of ecological generalism (ΔEG). Fitted lines show the univariate GAMs with 95% confidence interval (dark grey). Rugs on the x-axes show the predictor values and how they are distributed. Labels on the y-axes indicate the smooth functions for the term of interest (ΔSR, ΔPD, ΔEG and elevation) and the estimated degrees of freedom (following the term). Values above and below the horizontal dashed line indicate heterogenization and homogenization, respectively. Values left and right of the vertical dashed line indicate in (A) species loss and gain, (B) PD decrease and increase and (D) assemblages composed by specialists and generalists, respectively.

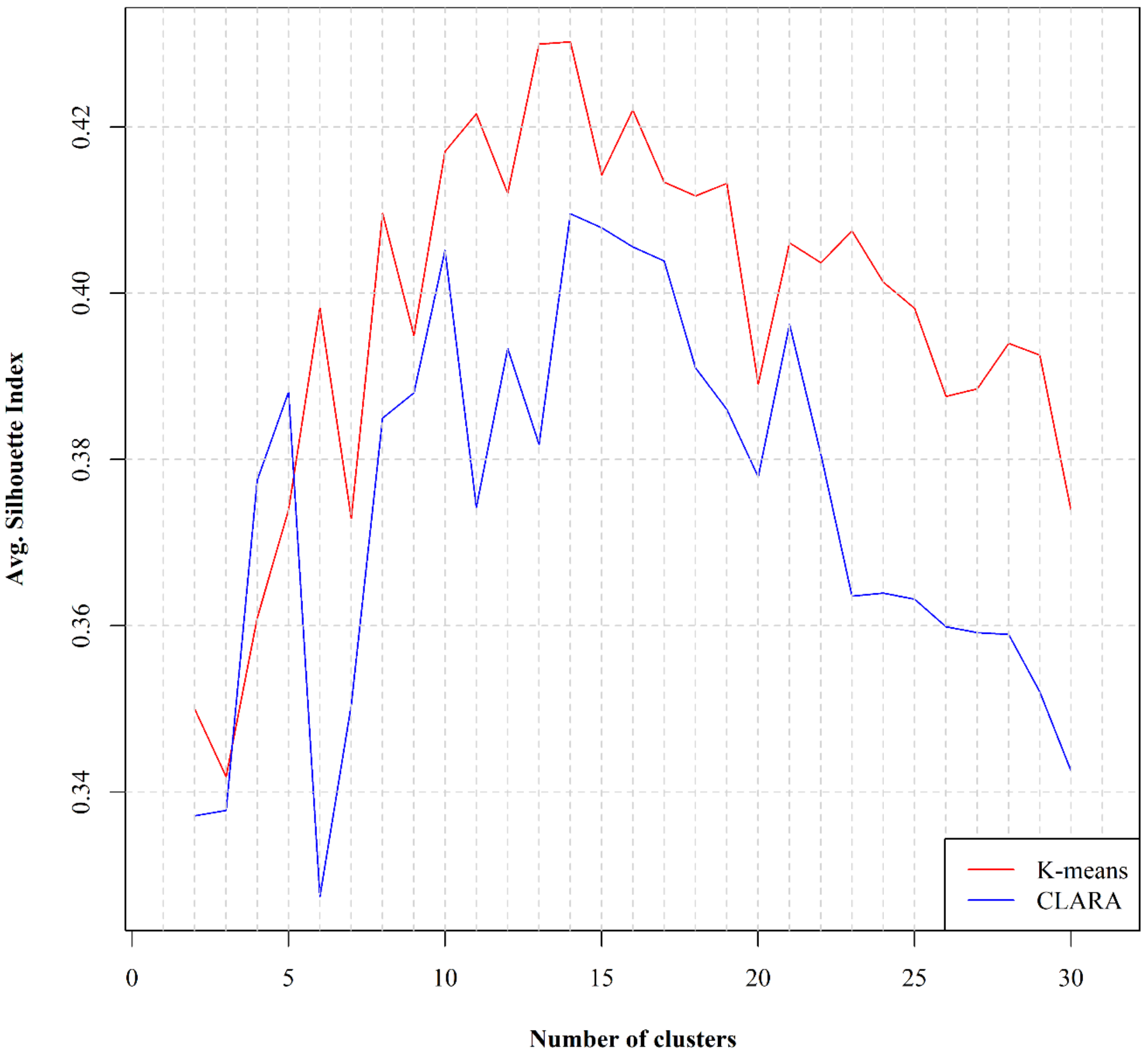


**Figure S45.** The values of the Silhouette index for the k-means and the CLARA unsupervised clustering algorithms regarding the optimal number of biogeographical regions (clusters) currently occurring in Crete.

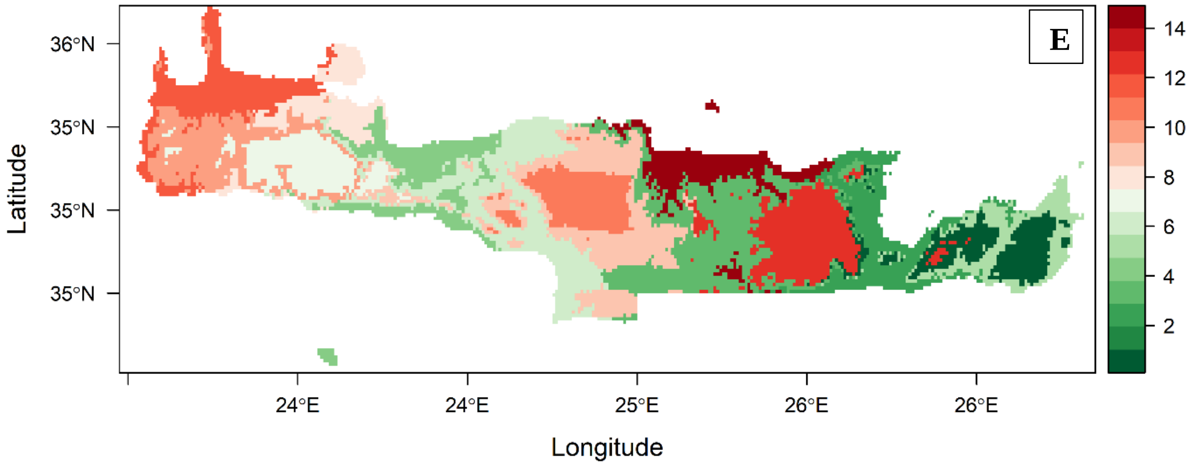


**Figure S46.** Current bioregionalization of Crete. Each colour indicates a different biogeographical region.

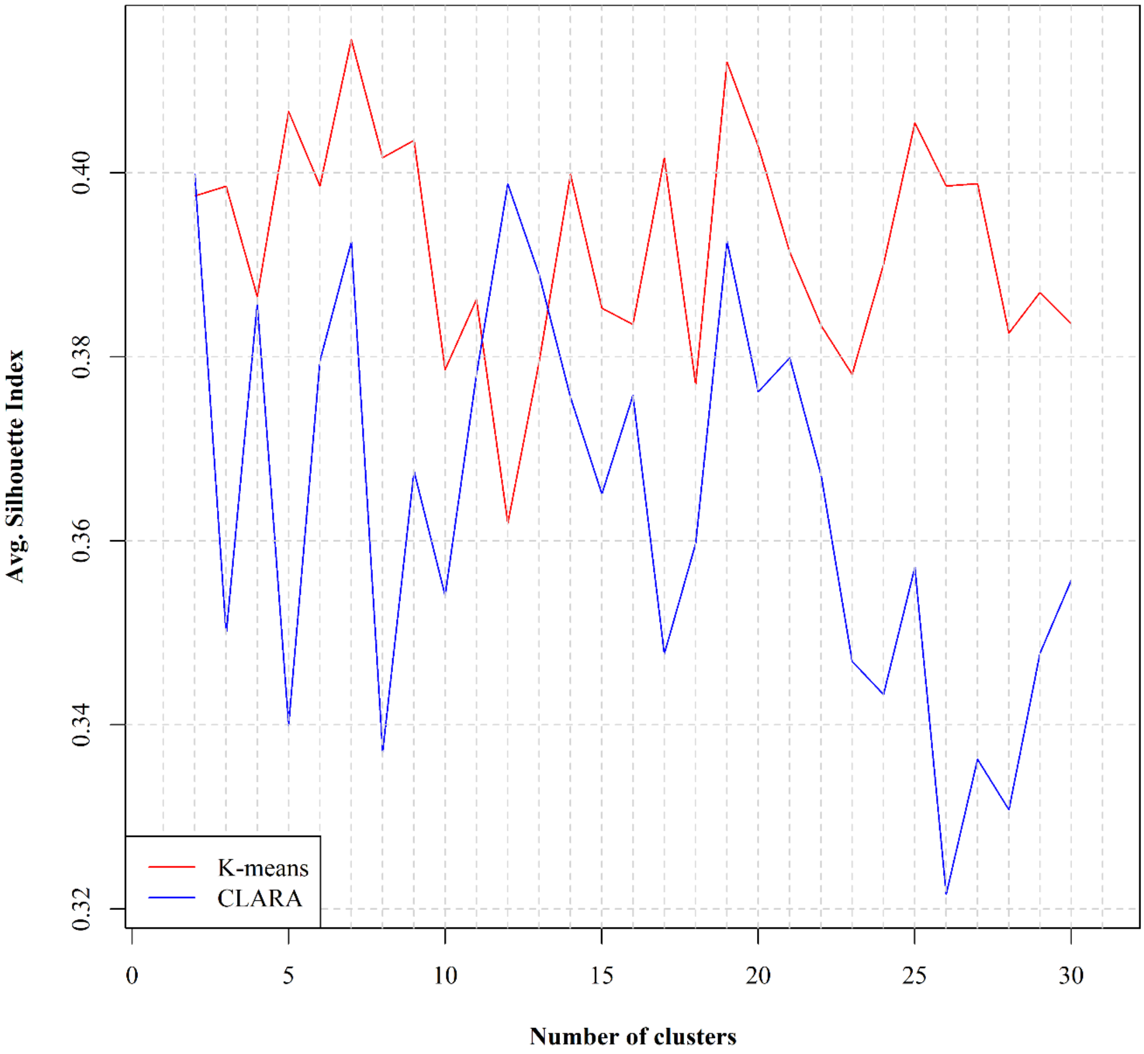


**Figure S47.** The values of the Silhouette index for the k-means and the CLARA unsupervised clustering algorithms regarding the optimal number of biogeographical regions (clusters) predicted to occur for the BCC 2.6 GCM/RCP combination in Crete.

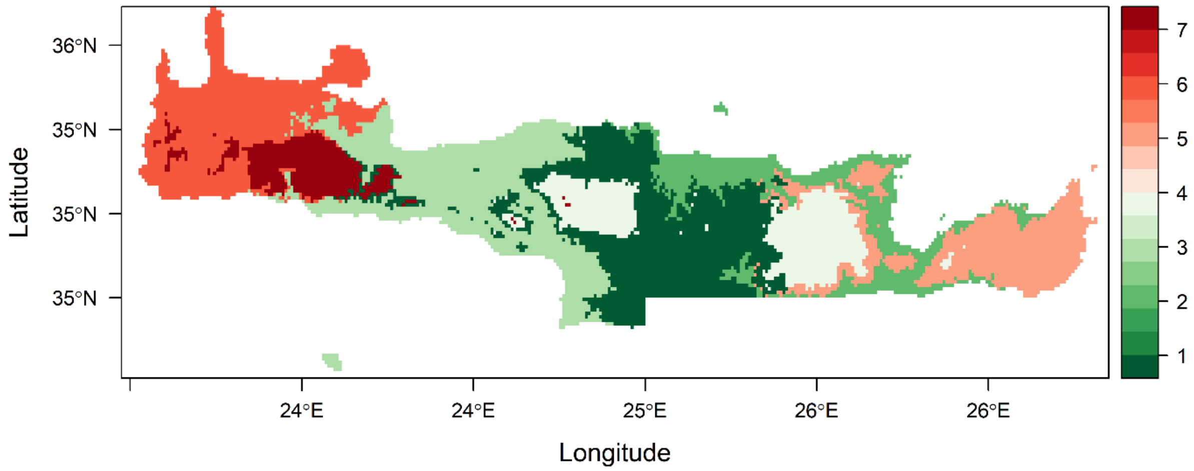


**Figure S48.** Bioregionalization of Crete for the BCC 2.6 GCM/RCP combination. Each colour indicates a different biogeographical region.

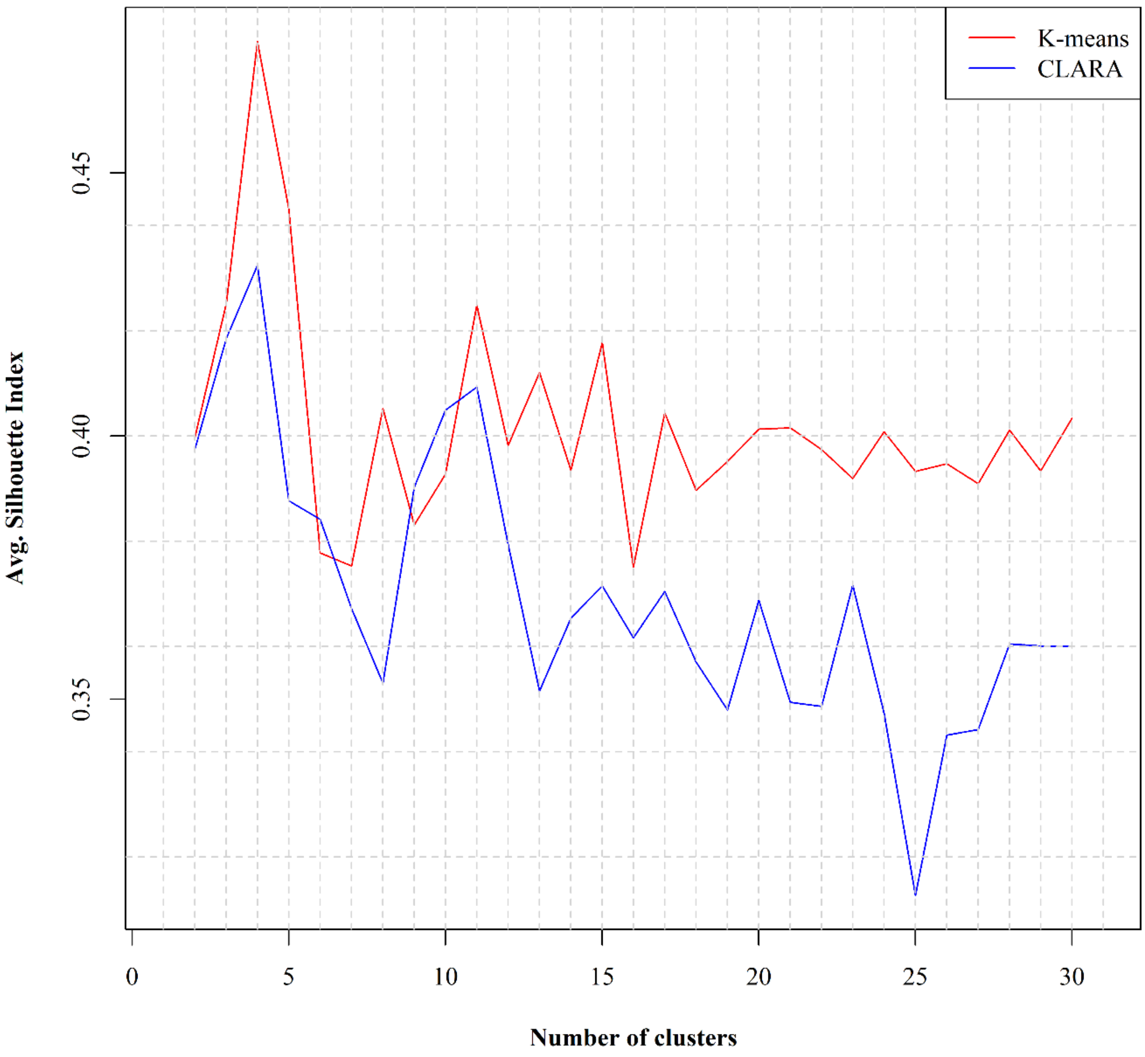


**Figure S49.** The values of the Silhouette index for the k-means and the CLARA unsupervised clustering algorithms regarding the optimal number of biogeographical regions (clusters) predicted to occur for the BCC 8.5 GCM/RCP combination in Crete.

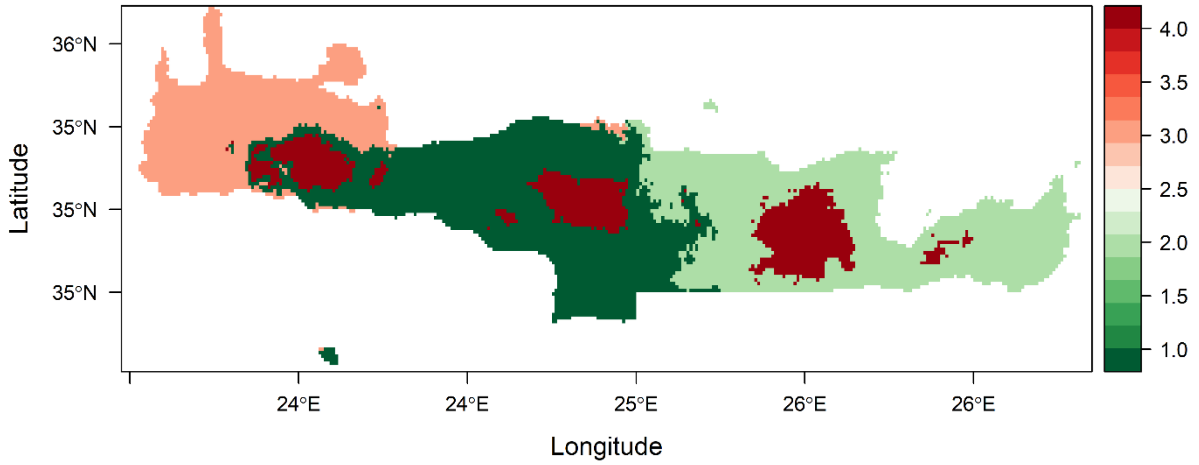


**Figure S50.** Bioregionalization of Crete for the BCC 8.5 GCM/RCP combination. Each colour indicates a different biogeographical region.

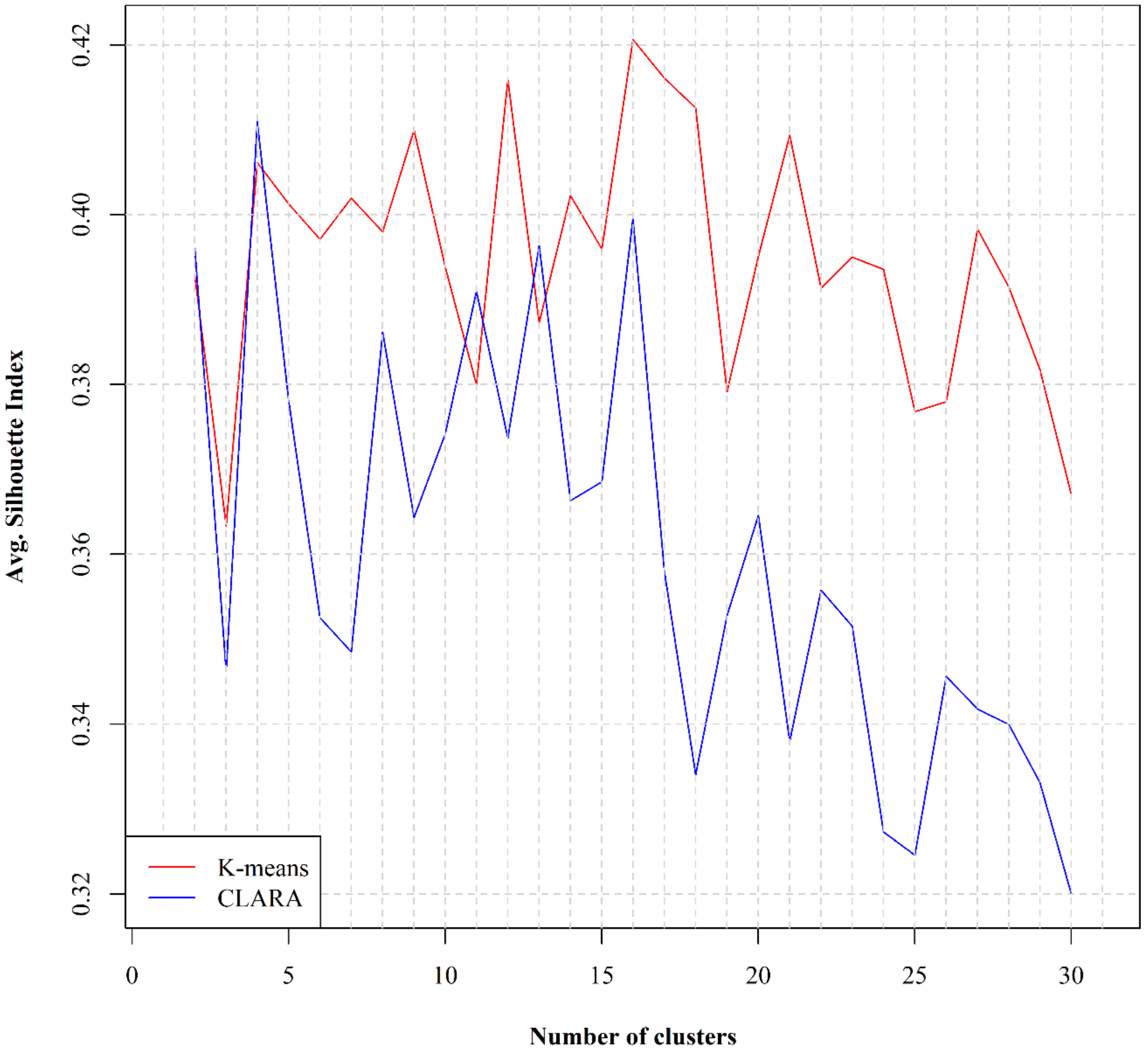


**Figure S51.** The values of the Silhouette index for the k-means and the CLARA unsupervised clustering algorithms regarding the optimal number of biogeographical regions (clusters) predicted to occur for the CCSM4 2.6 GCM/RCP combination in Crete.

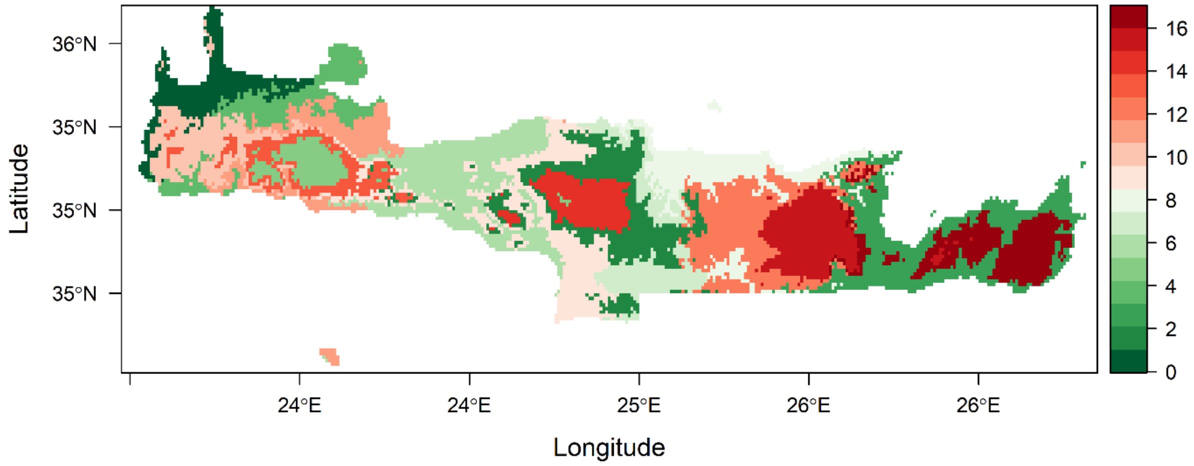


**Figure S52.** Bioregionalization of Crete for the CCSM4 2.6 GCM/RCP combination. Each colour indicates a different biogeographical region.

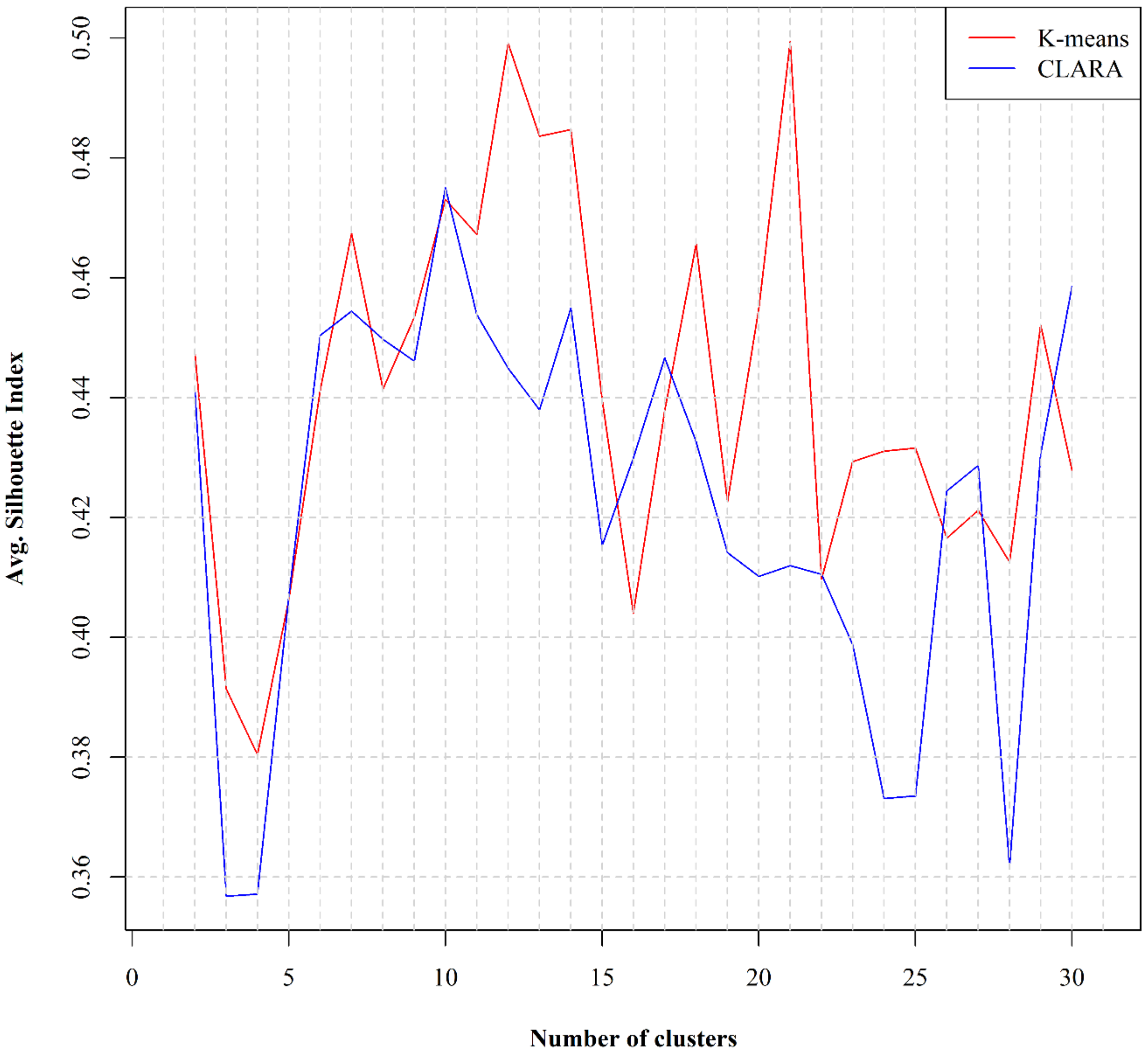


**Figure S53.** The values of the Silhouette index for the k-means and the CLARA unsupervised clustering algorithms regarding the optimal number of biogeographical regions (clusters) predicted to occur for the CCSM4 8.5 GCM/RCP combination in Crete.

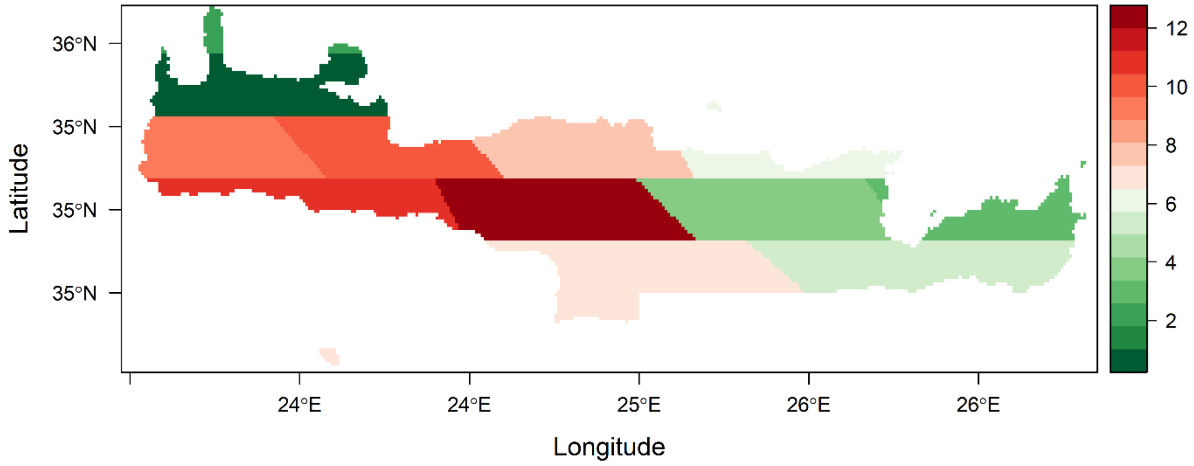


**Figure S54.** Bioregionalization of Crete for the CCSSM4 8.5 GCM/RCP combination. Each colour indicates a different biogeographical region.

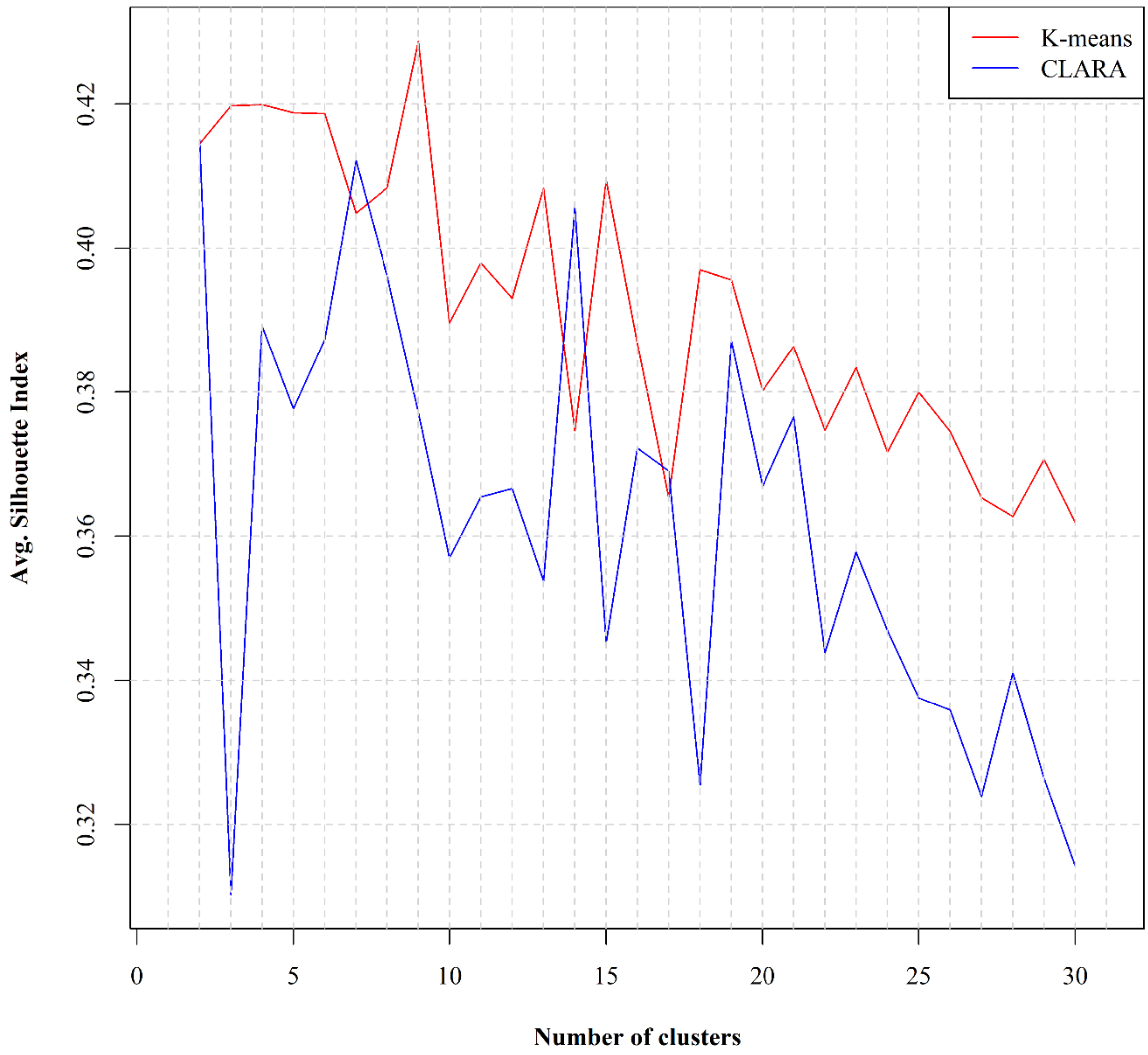


**Figure S55.** The values of the Silhouette index for the k-means and the CLARA unsupervised clustering algorithms regarding the optimal number of biogeographical regions (clusters) predicted to occur for the HadGEM2 2.6 GCM/RCP combination in Crete.

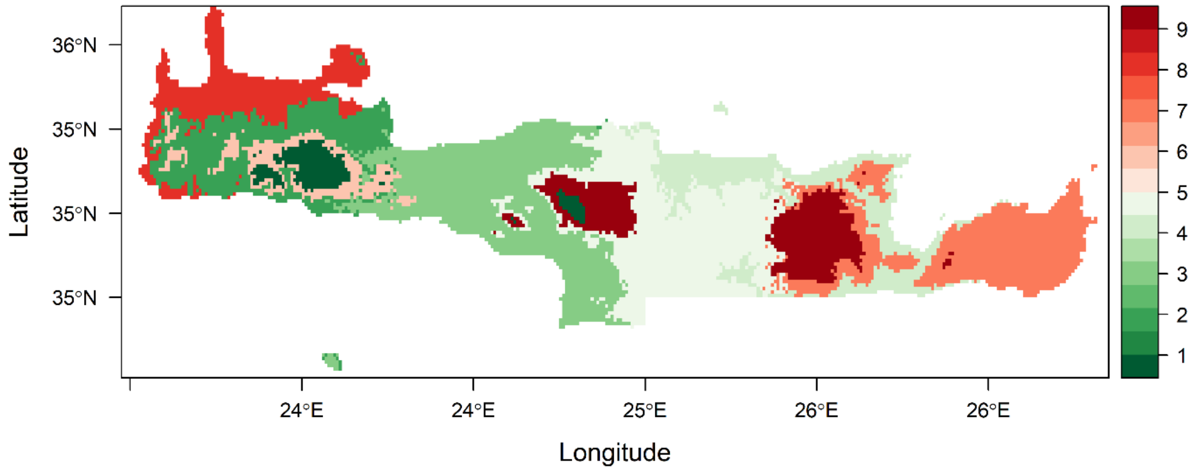


**Figure S56.** Bioregionalization of Crete for the HadGEM2 2.6 GCM/RCP combination. Each colour indicates a different biogeographical region.

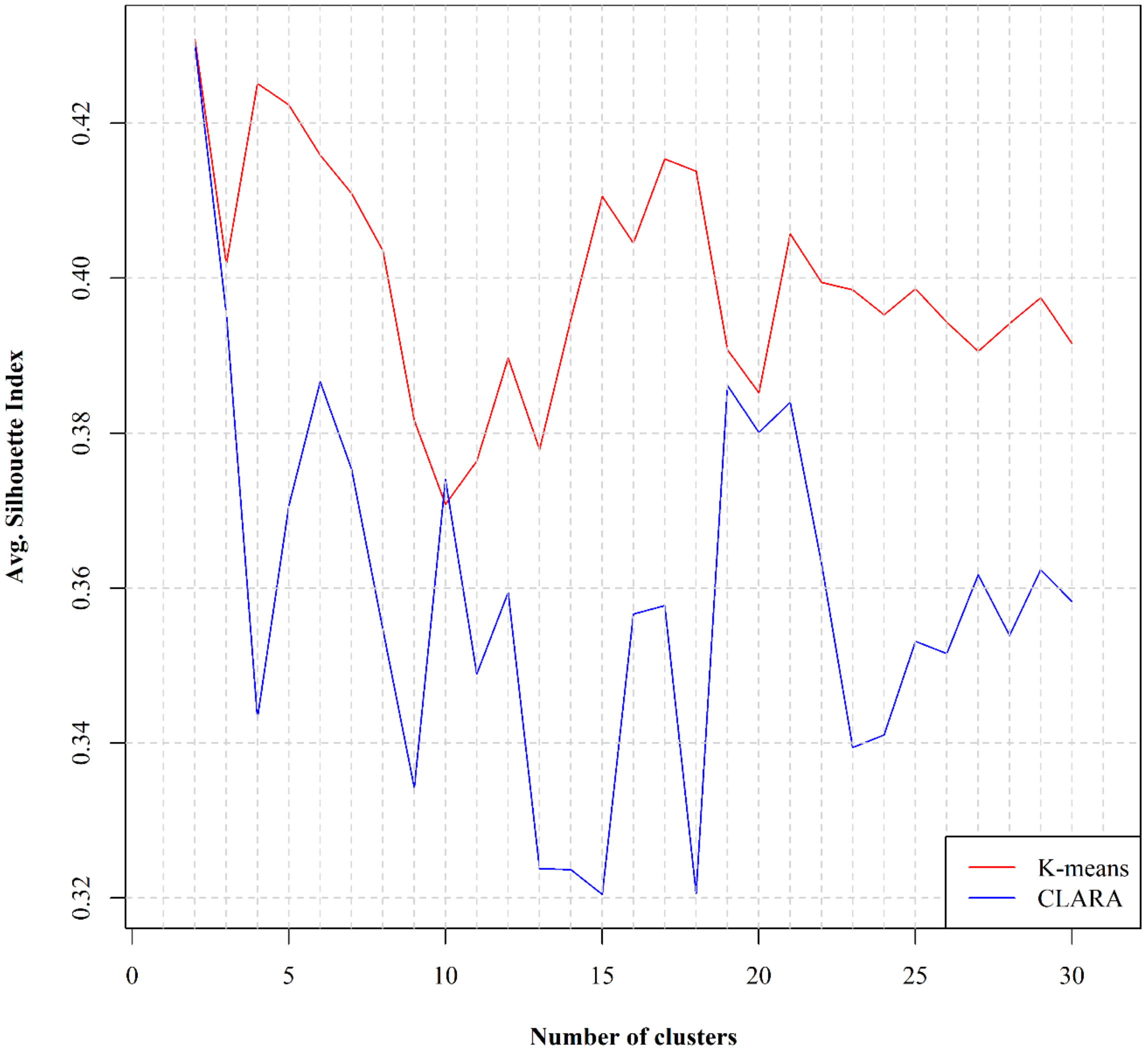


**Figure S57.** The values of the Silhouette index for the k-means and the CLARA unsupervised clustering algorithms regarding the optimal number of biogeographical regions (clusters) predicted to occur for the HadGEM2 8.5 GCM/RCP combination in Crete.

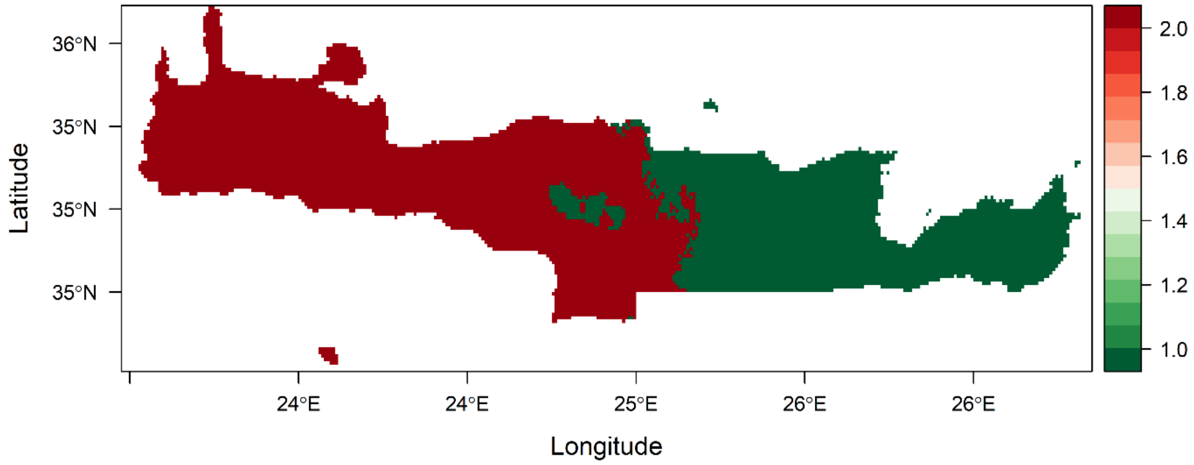


**Figure S58.** Bioregionalization of Crete for the HadGEM2 8.5 GCM/RCP combination. Each colour indicates a different biogeographical region.

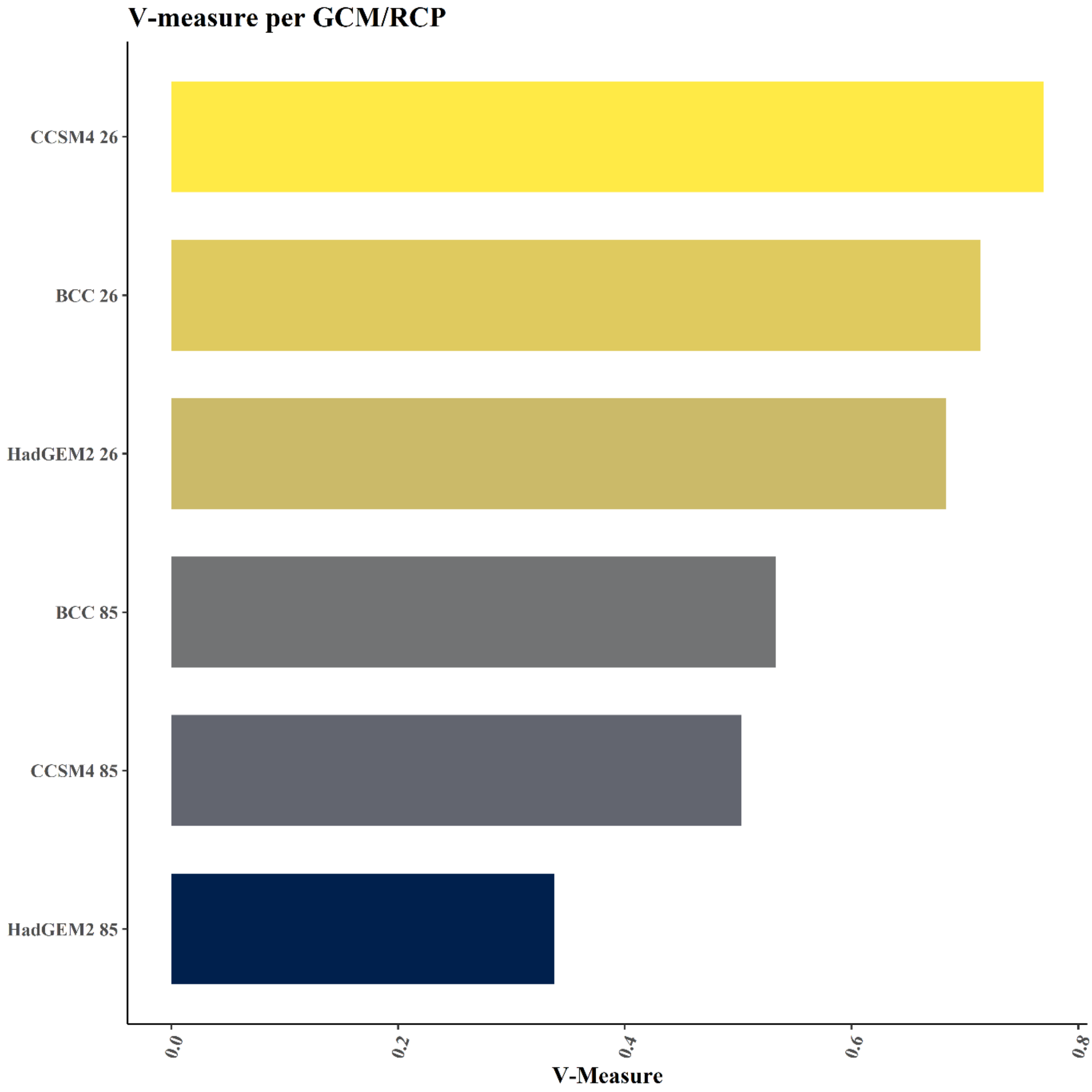


**Figure S59.** Similarity regarding Crete's bioregionalization schema between the present and each Global Circulation Model (GCM) and Representative Concentration Pathway (RCP), based on the V-measure index.

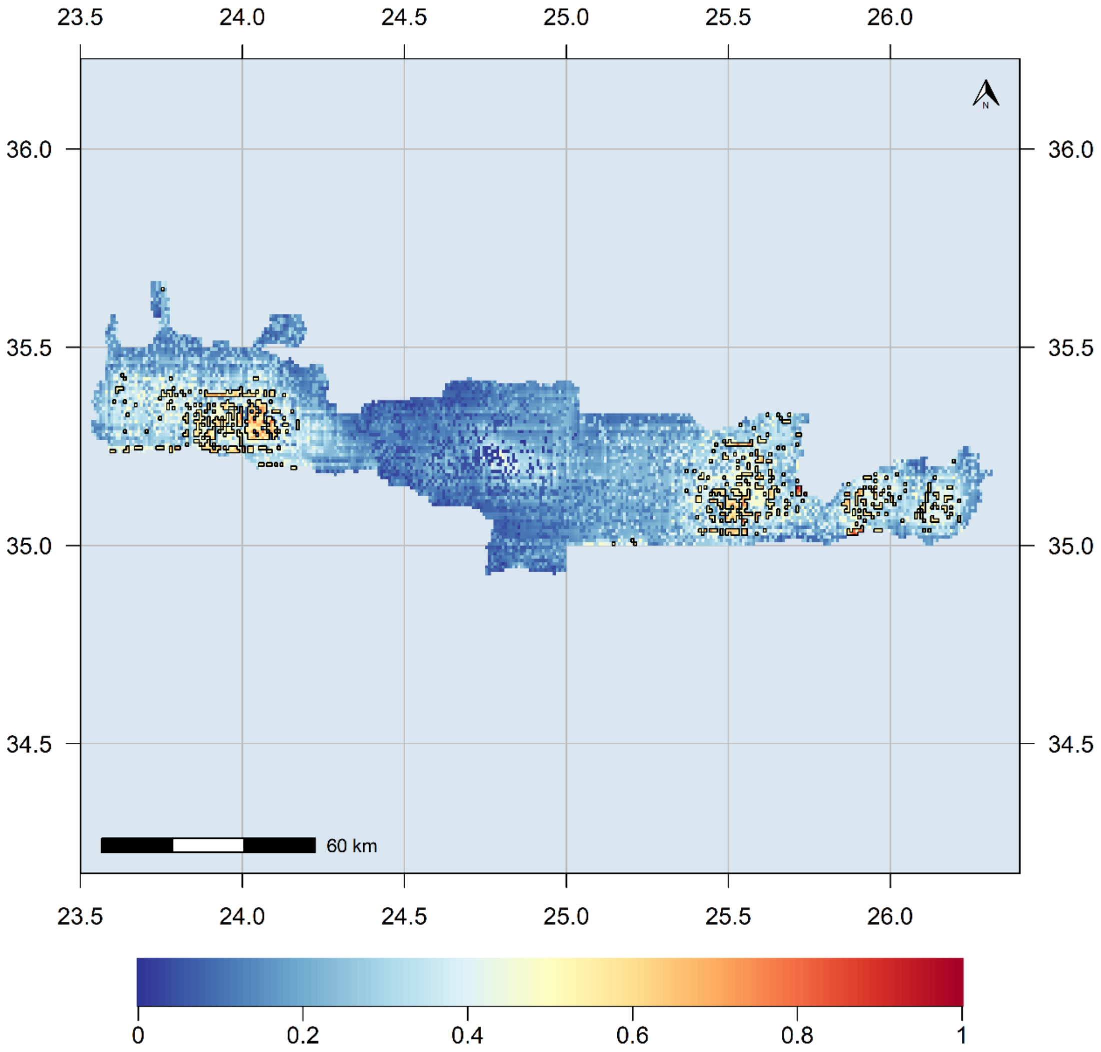


**Figure S60.** Index of climate stability sensu Owens and Guralnick (2019) and areas identified as climate refugia in Crete.

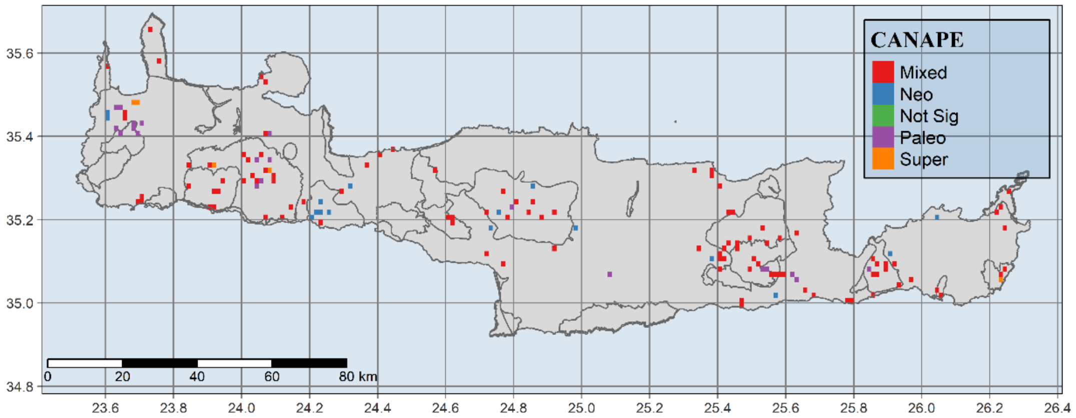


**Figure S61.** Map of the protected areas (PA) network in Crete overlaid onto the Categorical Analysis of Neo- and Paleo-Endemism results.

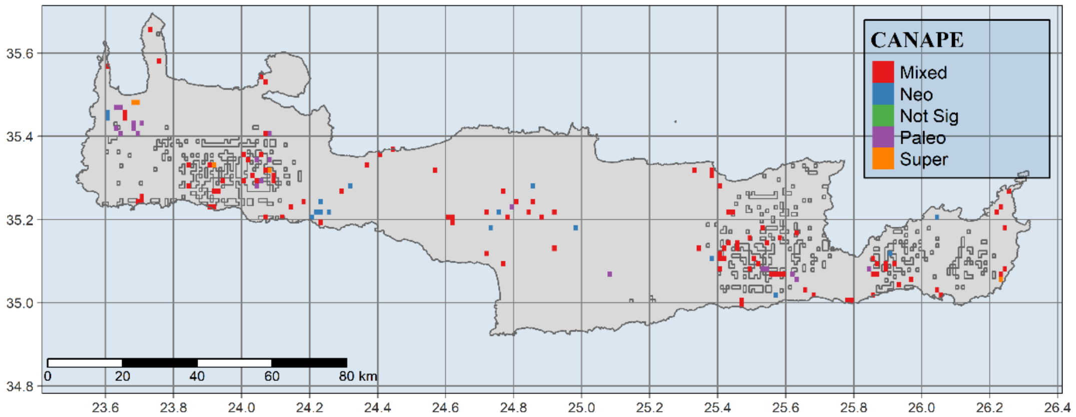


**Figure S62.** Map of the recognised climate refugia in Crete overlaid onto the Categorical Analysis of Neo- and Paleo-Endemism results.

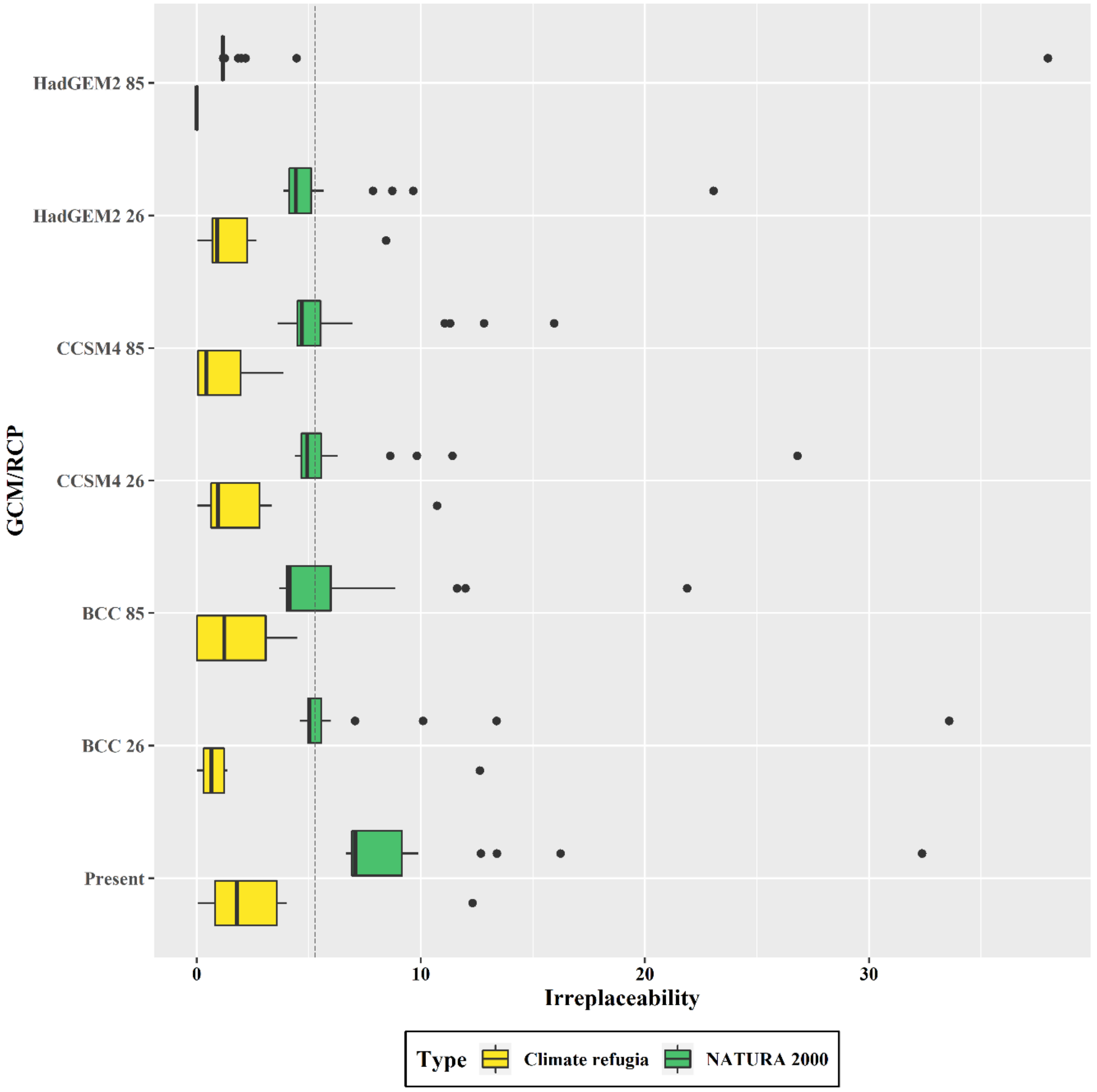


**Figure S63.** Mean irreplaceability index for the protected areas network (NATURA 2000 sites) and climate refugia in Crete, for the current and all the Global Circulation Model (GCM) and Representative Concentration Pathway (RCP) combinations considered in our study. The vertical dashed line represents the median irreplaceability index.

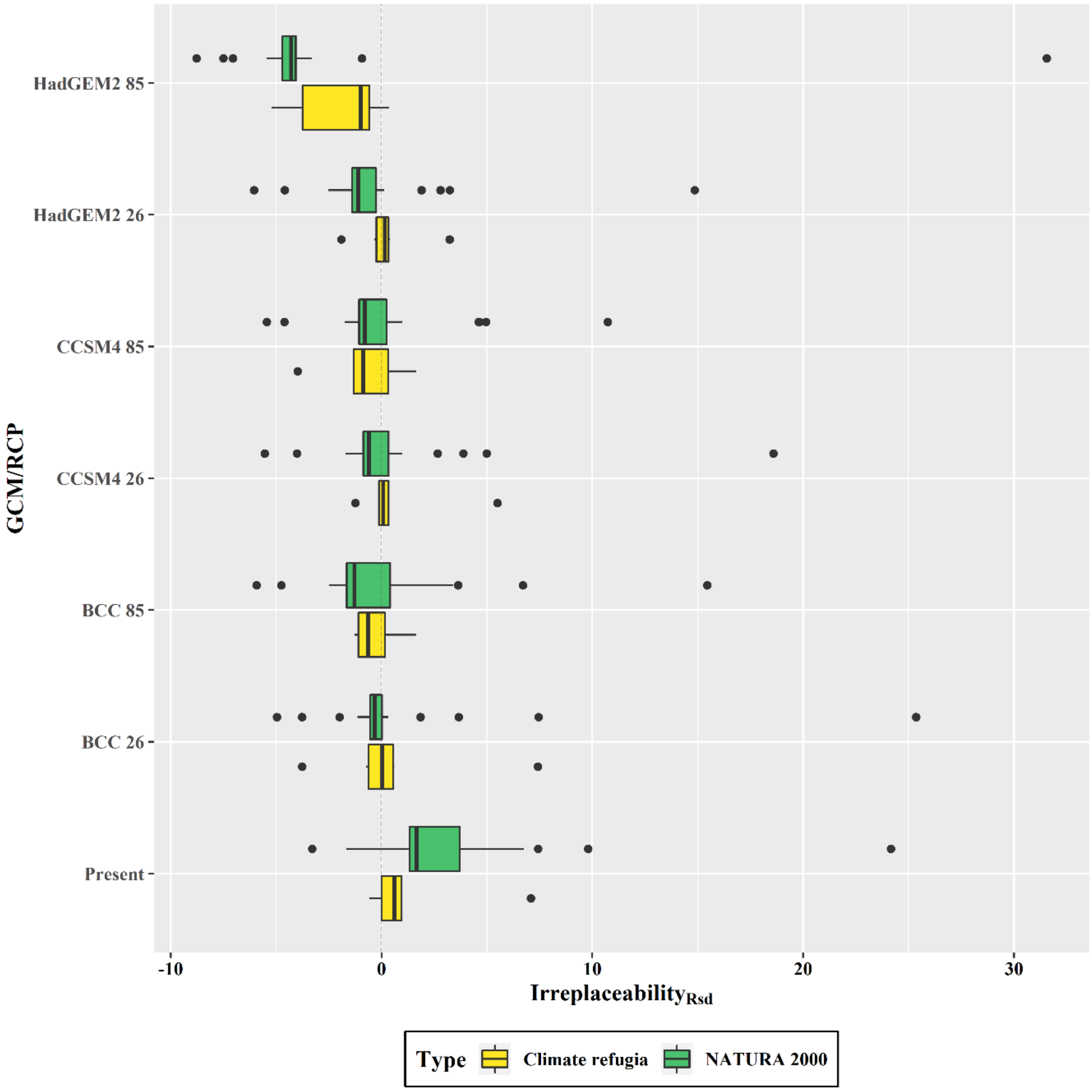


**Figure S64.** Mean irreplaceability index standardized for area, for the protected areas network (NATURA 2000 sites) and climate refugia in Crete, for the current and all the Global Circulation Model (GCM) and Representative Concentration Pathway (RCP) combinations considered in our study. The vertical dashed line represents the median irreplaceability index.
